# Phenotype of mice carrying an NMDA receptor GluN2B protein-truncating variant associated with intellectual disability

**DOI:** 10.1101/2025.10.02.680004

**Authors:** Klevinda Fili, Viktor Kuchtiak, Eni Tomovic, Miriam Candelas Serra, Agnieszka Kubik-Zahorodna, Karel Harant, Paulina Bozikova, Jiri Cerny, Miloslav Korinek, Barbora Hrcka Krausova, Vera Abramova, Mark Dobrovolski, Fatma Elzahraa S. Abdel Rahman, Jan Prochazka, Ales Balik, Tereza Smejkalova, Ladislav Vyklicky

**Author notes:** Co-corresponding authors: Prof. Ladislav Vyklicky, MD, PhD Institute of Physiology, Czech Academy of Sciences Videnska 1083, 142 00 Prague 4, Czech Republic Tereza Smejkalova, PhD Institute of Physiology, Czech Academy of Sciences Videnska 1083, 142 00 Prague 4, Czech Republic.

## Abstract

Pathogenic variants in *GRIN2B*, encoding the NMDA receptor (NMDAR) GluN2B subunit, are linked to intellectual disability (ID) and related neurodevelopmental disorders. While most disease-associated variants are missense, protein-truncating variants (PTVs) may cause haploinsufficiency with less severe phenotypes. Here, we characterize a knock-in mouse model carrying the GluN2B-L825Ffs*15 PTV (*Grin2b*^+/Δ^). Proteomic analysis revealed markedly reduced full-length GluN2B protein and no detectable truncated GluN2B, accompanied by a compensatory increase in GluN2A. Electrophysiology in hippocampal neurons demonstrated reduced NMDA-induced currents, diminished ifenprodil sensitivity, and accelerated NMDAR-mediated EPSC deactivation, consistent with a shift toward GluN2A-containing receptors. AMPAR-mEPSC amplitudes were increased, indicating altered excitatory synaptic function. Behaviorally, *Grin2b*^+/Δ^ mice exhibited hypoactivity, increased anxiety in males, and impaired sensorimotor gating in both sexes, while learning, memory, and social behaviors remained largely intact. These results demonstrate that a monoallelic GluN2B PTV alters NMDAR subunit composition and function, producing moderate behavioral effects, and provide insight into mechanisms underlying *GRIN2B*-associated ID.

## Introduction

Glutamate is the primary excitatory neurotransmitter in the central nervous system, acting on ionotropic NMDA, AMPA, and kainate receptors. NMDA receptors (NMDARs) are characterized by slow gating kinetics and high Ca^2+^ permeability, and they perform key synaptic computations underlying cognitive function and guiding nervous system development (Hansen et al., 2021). NMDARs are heterotetrameric cation channels comprising two obligatory GluN1 and two GluN2 and/or GluN3 subunits (Hansen et al., 2021). There are four types of GluN2 subunits (GluN2A-D), with distinct spatial and temporal expression patterns (Monyer et al., 1994) (Sheng et al., 1994) (Hanson et al., 2024) and with influence on receptor biophysical and pharmacological properties (Monyer et al., 1992; Vicini et al., 1998) (Hansen et al., 2014). The GluN2B subunit, whose expression peaks during embryonic and early postnatal development, is critical for synapse development, and various forms of synaptic plasticity (Kutsuwada et al., 1996) (Hansen et al., 2021).

A significant portion of severe neurodevelopmental disorders likely has *de novo* monogenic causes, with *GRIN* genes, encoding NMDAR subunits, among the implicated genes (López-Rivera et al., 2020). *GRIN* gene variants have been linked to developmental delay (DD), intellectual disability (ID), epilepsy, autism spectrum disorder (ASD) and schizophrenia (Platzer and Lemke, 1993) (Endele et al., 2010) (Tarabeux et al., 2011) (Hu et al., 2016) (Takasaki et al., 2016) (Benke et al., 2021). Over 700 disease-associated *GRIN* gene variants have been identified, predominantly in genes encoding GluN2A (43%), GluN2B (35%), and GluN1 (13%) subunits (Hansen et al., 2021; Korinek et al., 2024) (Perszyk et al., 2021). Most frequent are missense variants, accounting for 65% of the identified disease-associated *GRIN* gene variants, while 23% are protein-truncating variants (PTVs) (Santos-Gomez et al., 2021). Compared to missense variants, PTVs tend to be associated with a less severe DD/ID presentation in carriers (Platzer et al., 2017) (Strehlow et al., 2019) (Santos-Gomez et al., 2021).

Disease mechanisms underlying symptoms in patients with missense variants *vs.* PTVs are not known. At the molecular level, missense variants lead to a single amino acid substitution in the subunit protein that, when incorporated into NMDAR complexes, may alter various aspects of receptor function, including channel gating, allosteric modulation, post-translational modifications, protein-protein interactions, or trafficking (Amin et al., 2021) (Perszyk et al., 2021; Korinek et al., 2024). In contrast, it is thought that the products of most PTVs (with the exception of PTVs affecting the subunit C-terminal domain (Kysilov et al., 2024)) are degraded at the mRNA or the protein stage, or fail to assemble into stable NMDAR complexes, resulting in haploinsufficiency of the affected subunit (Santos-Gomez et al., 2021).

To study the circuit-level functional impact of different disease-causing *GRIN* gene variants animal models carrying clinically relevant variants are needed. In recent years several mouse strains carrying individual patient missense variants have been created and characterized, including two with patient variants in *Grin2b*, encoding GluN2B-C456Y (Shin et al., 2020) or GluN2B-L825V (Candelas Serra et al., 2024). To assess haploinsufficiency, heterozygous individuals of conventional *Grin2b* knock-out strains have been examined (Shin et al., 2020), but such data are very limited so far.

To investigate the functional consequences and possible compensatory mechanisms linked to disease-causing GluN2B PTVs, we generated a knock-in mouse model carrying GluN2B-L825Ffs*15 (GluN2B^Δ^) frameshift variant in the *Grin2b* gene, creating a premature termination codon in the transmembrane domain M4 helix. We observed severely impaired cell surface expression of the GluN2B^Δ^ subunit in heterologous expression systems. In the hippocampal tissue of heterozygous (*Grin2b*^+/Δ^) mice we found decreased mRNA levels of *Grin1* and *Grin2b*, accompanied by reduced GluN2B and slightly increased GluN2A protein levels. Cultured hippocampal neurons from *Grin2b*^+/Δ^ mice exhibited smaller NMDA-induced current amplitudes and accelerated NMDAR-mediated evoked excitatory postsynaptic current (eEPSC) deactivation kinetics. Together, our data suggest that the presence of the *Grin2b*^Δ^ variant leads to altered receptor subunit composition, increasing the relative contribution of GluN2A-containing NMDARs. Behaviorally, *Grin2b*^+/Δ^ mice showed hypoactivity, increased anxiety, altered sensorimotor gating, but no apparent cognitive impairment. This study provides insights into the biological consequences of *Grin2b* PTVs and may inform the development of personalized therapies for patients with different variants in *GRIN2B*.

## Methods

### Ethics statement

Animals were bred and maintained according to the guidelines of the European Union Directive 2010/63/EU and in full compliance with the Czech Act No. 246/1992 Coll. and Decree No. 419/2012 Coll., regarding the protection of animals used for scientific purposes. All experiments were approved by the Animal Care and Use Committee of the Institute of Molecular Genetics, Czech Academy of Sciences, and by the Resort Professional Commission of the Czech Academy of Sciences for Approval of Projects of Experiments on Animals.

### Animals

Heterozygous mice (C57BL/6N background) carrying the GluN2B-L825Ffs*15 mutation in the *Grin2b* gene (*Grin2b*^+/Δ^) were obtained using CRISPR/Cas9 genome-editing system (Candelas Serra et al., 2024). Cas9 protein, gRNA, and ssDNA were transfected into the zygote by electroporation. In the process of generating the mouse strain carrying the GluN2B-L825V variant (Candelas Serra et al., 2024) we also obtained mice with the GluN2B-L825Ffs*15 variant. Results of genome editing were confirmed by PCR amplification in the founder mice and the presence of the frameshift in individual mice was also confirmed by direct Sanger sequencing of PCR amplicons. Heterozygous mice were obtained from breeding *Grin2b*^+/Δ^ with wild-type mice. Newborn *Grin2b*^+/Δ^ pups and their wild-type littermates (*Grin2b*^+/+^) were used for the preparation of primary neuronal cultures. For behavioral experiments, mice were housed under standard 12:12 h light/dark cycle and received food and water ad libitum. Animals underwent behavioral testing at 9–15 weeks of age.

### RNA isolation and RT-qPCR

Total RNA was extracted from the individual hippocampi of 4-month-old male and female mice. Brain tissue was frozen on dry ice immediately after dissection and stored at -80°C until RNA isolation. The tissue samples were homogenized in TRI Reagent (Merck), and total RNA was extracted according to the manufacturer’s protocol. The RNA pellet was resuspended in 20 µl of RNase-free water and the concentration of RNA was measured using a Nano Drop instrument (Thermo Scientific). We used 1 µg of total RNA for cDNA synthesis. We combined 250 ng of random primers (Promega) with total RNA, the mixture was incubated at 70°C for 5 min and then cooled on ice for 2 min. Subsequently, cDNA was synthetized (at 37°C, for 90 min) in a 25 µl reaction containing 200 U of M-MLV Reverse Transcriptase (Promega). The obtained cDNA was additionally diluted by 25 µl of PCR grade water and stored at 4°C until RT-qPCR analysis.

TaqMan qPCR reactions were performed in 20 µl qPCR reaction mixture containing 1.5 µl cDNA samples, 10 µl 2x TaqMan Gene Expression Master Mix, and 2 µl TaqMan Gene Expression Assay (Thermo Fisher Scientific Inc.) containing the probe and the appropriate forward and reverse primers (Table S1). All qPCR reactions were performed in triplicate on a LightCycler 480 instrument (Roche Diagnostics). Relative gene expression levels (RQ values) of target genes were analyzed using the ΔΔCt method (Livak and Schmittgen, 2001; Taylor et al., 2019), where Ct refers to the threshold cycle value. The mean Ct values obtained from qPCR were normalized to the levels of housekeeping genes glyceraldehyde 3-phosphate dehydrogenase (GAPDH) and beta-2-microglobulin (β2M) (Table S1).

### Mass spectrometry analysis

Mass spectrometry analysis was performed as previously described (Candelas Serra et al., 2024). Hippocampal samples from 4-month-old male mice (*n* = 4 for *Grin2b*^+/+^ and *n* = 5 for *Grin2b*^+/Δ^) were lysed by multiple rounds of sonication in 2% sodium deoxycholate detergent solution in 100 mM triethylammonium bicarbonate buffer. Cysteines were reduced with Tris(2-carboxyethyl) phosphine and alkylated with chloroacetamide. Proteins were digested with trypsin, labeled with 16-plex tandem mass tags (TMTpro), and fractionated by reverse-phase chromatography at pH 10. The resulting eight fractions were analyzed using nanoLC coupled with a Thermo Orbitrap Lumos mass spectrometer, with TMT quantification performed in triple-stage mass spectrometry (MS3). Data processing was conducted in ProteomeDiscoverer 2.5 (Thermo Scientific, USA), using TMT reporter ion ratios to estimate relative protein abundance. The database search was performed against the *Mus musculus* UniProt reference database and a common contaminant database. Fixed modifications included TMTpro labeling at the peptide N-terminus and lysine (Unimod #2016) and cysteine carbamidomethylation (Unimod #39), while variable modifications included methionine oxidation (Unimod #1384) and protein N-terminal acetylation (Unimod #1). Proteins and peptides were filtered to a 1% false discovery rate (FDR).

Data were normalized to total peptide amount, and statistical analysis was performed in Perseus software (Tyanova and Cox, 2018). Contaminants were removed, intensity values were log₂-transformed, and proteins with insufficient quantification (fewer than three valid values in at least one experimental group) were excluded. A Student’s t-test with permutation-based FDR correction was applied to identify significantly altered proteins at a 5% FDR threshold. ASD-related genes from the SFARI database (https://gene.sfari.org) were annotated.

To estimate the protein copy number (CN) per cell, the “proteomic ruler” method (Wisniewski et al., 2014) was used. This estimation utilized the mass spectrometry signal of histones, which correlates with DNA content and thus the number of cells in a sample. CN calculations were performed using the Proteomics Ruler Perseus plugin, with the scaling mode set to total protein amount.

### Calculation of proportions of diheteromeric and triheteromeric NMDA receptors

Mass spectrometry provided the CN for the GluN1, GluN2A and GluN2B subunits of NMDA receptors (*CN*_1_, *CN*_2A_, *CN*_2B_). To estimate the relative proportions of NMDA receptor subtypes based on subunit composition, we assumed that each receptor tetramer comprises two obligatory GluN1 subunits and two additional subunits randomly occupied by either GluN2A or GluN2B. The contribution of other GluN2 or GluN3 subunits was considered negligible for the purpose of this analysis. The total number of assembled receptors depends on whether there are fewer GluN1 or GluN2 subunits. It is equal to the smaller number of *CN*_1_/2 or (*CN*_2A_ + *CN*_2B_)/2. Out of all GluN2 subunits (*CN*_2A_ + *CN*_2B_), the proportions of GluN2A and GluN2B are equal to:

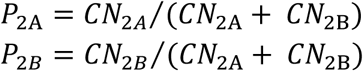

The probability that a receptor with a 1-1-2A-2A subunit composition assembles and therefore the proportion of receptors with a 1-1-2A-2A subunit composition (*P*_2A-2A_) is equal to *P* ^2^:

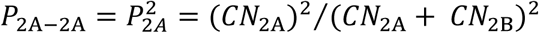

Similarly, out of all assembled receptors, the proportions of 1-1-2B-2B and 1-1-2A-2B receptors, respectively, are given below. The last expression contains the factor of 2 since combinations of 2A-2B and 2B-2A provide receptors with the same subunit composition.

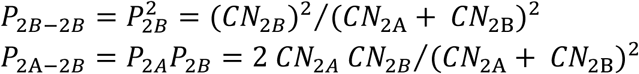

### Homology modeling

The initial models of the murine GluN1/GluN2B complex were constructed based on a previously developed rat NMDAR homology model, which was derived from the available crystal structures 4pe5, 4tll, and 4tlm (Karakas and Furukawa, 2014; Lee et al., 2014). The template receptor geometry corresponded to the ligand-activated state (RAA) as obtained from previous opening/closing simulations (Cerny et al., 2019). Three models of the GluN1/GluN2B complex were generated using the automodel function in MODELLER 9.23 (Sali and Blundell, 1993; Webb and Sali, 2014). Each model was assembled with two GluN1 subunits and all possible combinations of two GluN2B subunits, either incorporating or omitting the truncation mutation. The GluN1 subunit models encompassed residues 23 to 847, in accordance with the UniProt record P35438 (UniProt, 2021), excluding the last 91 residues of the CTD, which are disordered. The GluN2B subunits containing the truncation mutation in the transmembrane helix M4 included residues 30 to 824, followed by the sequence FFGGSHGSQPHHLHL. In contrast, the wild-type GluN2B models incorporated residues 30 to 852. The largely disordered CTD, comprising residues 853 to 1482, was excluded from the modeling process.

### Molecular dynamics simulations

Molecular dynamics (MD) simulations of the murine GluN1/GluN2B receptor model were prepared using CHARMM-GUI (Jo et al., 2008; Lee et al., 2016). The receptor was embedded in a model membrane composed of DOPC and cholesterol (3:1 ratio). The simulation box (160 Å × 160 Å × 220 Å) was solvated with TIP3P water, and Na⁺ and Cl⁻ ions were added to neutralize the system at a final salt concentration of 150 mM. Following the CHARMM-GUI equilibration protocol, 500 ns production MD simulations were conducted using GPU-accelerated GROMACS 2023.2 (Abraham et al., 2015) with the CHARMM36 force field (Klauda et al., 2010). A 2 fs time step was used, with short-range Coulomb and van der Waals cutoffs set to 1.2 nm, while long-range electrostatics was handled using Particle Mesh Ewald (PME) (Essmann et al., 1995). Atomic coordinates were recorded every 10 ps. The NPT ensemble simulation was controlled using the Nosé-Hoover thermostat at 303.15 K, with a coupling parameter of 1.0 ps, applying separate thermal baths for the solute, membrane, and solvent. The Parrinello-Rahman barostat maintained the system at 1.0 atm, with a coupling time constant of 5.0 ps and a compressibility factor of 4.5 × 10⁻⁵. MD geometry snapshots were superposed using the GROMACS trjconv -fit rot+trans command, with the first GluN1 chain as the reference for least-squares fitting. Visualization, graphical representation, and geometric analysis of selected regions were performed using PyMOL version 2.6.

### Analysis of subunit cell surface expression

#### DNA Constructs

The following cDNAs encoding human NMDA receptor subunits were used: GluN1-1a (GluN1; NM_007327), GluN2A (NM_001134407), and GluN2B (NM_000834), all cloned into the pCI-neo expression vector (a generous gift from prof. Stephen Traynelis, Emory University). To analyze receptor surface expression, a green fluorescent protein (GFP) tag was introduced into the extracellular portion of GluN2B directly after the signal peptide using the In Vivo Assembly (IVA) cloning method (Garcia-Nafria et al., 2016). The resulting construct, GluN2B-GFP, was subsequently transferred via restriction enzyme cloning into the pLEX-MCS expression vector (OpenBiosystems, Thermo Scientific) for lentiviral particle production and subsequent GluN2B-GFP expression in neurons (Kysilov et al., 2024) (Candelas Serra et al., 2024) (Kuchtiak et al., 2024). To replicate the genetic modification present in our mouse model, insertion of four thymine bases at the position L825 was achieved using IVA cloning, creating a frameshift mutation identical to the murine model. To ensure the same premature termination codon after 15 amino acids, an additional synonymous mutation at C838 was introduced by IVA mutagenesis, changing the triplet TGC (wild-type sequence) to TGT, thereby generating the mutant construct GluN2B-L825Ffs*15-GFP (GluN2B^Δ^-GFP) in both pCI-neo and pLEX-MCS vectors. Primer design was performed manually, and oligonucleotides were purchased from Eurofins Genomics (Germany). Following PCR amplification using Q5 DNA Polymerase (NEB), DpnI-treated products were transformed into ultracompetent XL10-Gold E. coli cells (Agilent, Santa Clara, USA). Clones were grown overnight, plasmid DNA was extracted, and sequencing (Eurofins Genomics) was performed to confirm the presence of the intended variant. Amino acid numbering follows the full-length protein sequence, including the signal peptide, with the initiating methionine designated as position 1.

#### Cell culture and transfection

Human embryonic kidney 293T (HEK293T) (American Type Culture Collection, ATCC No. CRL-1573, Rockville, MD, USA and monkey COS-7 cells were cultured as described earlier (Kysilov et al., 2024). Surface expression experiments were performed 24 h (microscopy) or 48 h (western blot) after the transfection of HEK293T cells and 48 h after the transfection of COS-7 cells with cDNA encoding GluN1 and GluN2A-GFP, GluN2B-GFP or GluN2B^Δ^-GFP subunits. Transfection was carried out using the GenJet reagent (SignaGen) following the manufacturer’s protocol. Following transfection, cells were maintained in culture medium supplemented with NMDAR antagonists D-(-)-2-amino-5-phosphonopentanoic acid (AP-5, 50 μM), Mg^2+^ (20 mM), and ketamine (1 μg/ml). Electrophysiology experiments were conducted 24–48 h after the transfection of HEK293T cells with cDNA encoding human GluN1 and GluN2B subunits, along with GFP (pQBI 25, Takara, Tokyo, Japan) as a marker of successful transfection, using Matra-A transfection reagent (IBA, Göttingen, Germany) as previously described (Borovska et al., 2012). For the analysis of recombinant subunit cell surface expression, primary neuronal cultures were prepared from mouse hippocampi on postnatal day 1 (P1) and cultured under standard conditions using Neurobasal A medium supplemented with B27 (Gibco/Invitrogen). Neurons were transduced with lentiviruses containing genes for GluN2B-GFP or GluN2B^Δ^-GFP expression, following previously established protocols (Kysilov et al., 2024) (Kuchtiak et al., 2024).

#### Analysis of surface expression by SDS-PAGE and western blot in HEK293T cells

Experiments were performed 48 h after the co-transfection of HEK293T cells with GluN1 and either GluN2B-GFP or GluN2B^Δ^-GFP subunits. NMDAR antagonists (AP-5, 50 μM; Mg^2+^, 20 mM; ketamine, 1 μg/ml) were added to the culture medium to prevent receptor activation. Cells were detached using 12.5 mM ethylenediaminetetraacetic acid (EDTA) at 37°C for 30 min (Kaur and Esau, 2015), collected by centrifugation, resuspended in phosphate-buffered saline (PBS), and fractionated. One quarter of the cell suspension was used as the total cell lysate (T) and incubated in 0.5% NP-40/Tris buffer with Halt Protease Inhibitor Cocktail (Thermo Scientific). The remaining cells were fractionated using the NEXT protocol (Rampado et al., 2022). Cells were first treated with EB1 buffer (0.015% digitonin) to isolate the cytosolic fraction (S1), followed by EB2 buffer (0.5% Triton X-100, Benzonase Nuclease) to obtain the cytoplasmic membrane (S2) and nuclear/organelle (P2) fractions. Samples were mixed with 4× Laemmli buffer, denaturated at 95°C, separated on 4–20% Tris-glycine SDS-PAGE gels, and transferred to nitrocellulose membranes. Membranes were blocked in 5% milk/PBS-Tween and incubated with primary antibodies: anti-GFP (1:2000; 11-476; EXBIO) for GFP-tagged GluN2B, anti-GAPDH (1:2000; 11-942; EXBIO) as a cytosolic marker, and anti-histone H3 (1:3000; ab1791; Abcam) as a nuclear marker. After washing, a horseradish peroxidase (HRP)-conjugated secondary antibody (1:6000, Thermo Scientific) was applied, signals were detected using SuperSignal West Pico substrate (Thermo Scientific) and imaged with a CCD camera system (Vilber Loumart). GAPDH verified cytosolic enrichment of the S1 fraction and histone H3 confirmed nuclear enrichment of the P2 fraction.

### Analysis of surface expression by a colorimetric assay in COS-7 cells

The colorimetric surface expression assay was performed as previously described (Kysilov et al., 2024) using COS-7 cells co-transfected with GluN1 and either GluN2B-GFP or GluN2B^Δ^-GFP subunits and maintained in culture medium supplemented with NMDAR antagonists (AP-5, 50 μM; Mg^2+^, 20 mM; ketamine, 1 μg/ml). To assess surface and total receptor expression, two 24-well plates were used for each experiment, with each condition run in quadruplicate. Cells transfected with GluN1 alone served as background controls. Cells were washed and fixed in 4% paraformaldehyde (PFA) before treatment with either the surface antigen blocking solution (PBS containing 50 mM NH_4_Cl, 10% goat serum, and 2% fish gelatin) or the permeabilizing blocking solution (PBS with 50 mM NH₄Cl, 10% goat serum, 2% fish gelatin, 0.1% Triton X-100, and 0.1% Tween-20). After blocking, cells were incubated with a rabbit anti-GFP primary antibody (1:500; 11-476-C100; EXBIO) followed by a goat anti-rabbit HRP-conjugated secondary antibody (1:500; AQ2P; Sigma-Aldrich). Detection was performed using o-phenylenediamine dihydrochloride (SIGMAFAST OPD tablet set; P9187; Sigma-Aldrich), and absorbance was measured at 492 nm using an Infinite PRO 200 multifunction modular reader (Tecan). Surface and total receptor expression values were obtained after background subtraction. The average value of surface labelling was subtracted from the average value of total labelling thus obtaining the value of intracellular labelling. Surface expression was determined as the ratio of surface to intracellular values.

### Analysis of surface expression by fluorescence microscopy in HEK293T cells

Fluorescence microscopy-based surface expression analysis was performed as previously described (Kysilov et al., 2024). HEK293T cells were co-transfected with GluN1 along with GluN2A and/or GluN2B-GFP or GluN2B^Δ^-GFP subunits and maintained in culture medium supplemented with NMDAR antagonists (AP-5, 50 μM; Mg^2+^, 20 mM; ketamine, 1 μg/ml). Experiments were performed 24 h post-transfection. Cells were washed with PBS, fixed in 4% PFA, and blocked using the surface antigen blocking solution (PBS containing 50 mM NH₄Cl, 10% goat serum, and 2% fish gelatin). Surface expression was assessed by incubating cells with a rabbit anti-GFP primary antibody (1:1000; AB3080P; Merck) followed by a goat anti-rabbit Alexa Fluor 647 secondary antibody (1:1000; A-21244; Invitrogen). Intracellular receptor expression was evaluated following blocking and permeabilization (PBS containing 50 mM NH₄Cl, 10% goat serum, 2% fish gelatin, 0.1% Triton X-100, and 0.1% Tween-20) using the same primary antibody and a goat anti-rabbit Alexa Fluor 555 secondary antibody (1:1000; A-21428; Invitrogen). 4′,6-diamidino-2-phenylindole (DAPI) was used to visualize cell nuclei. Images were acquired using a Leica DMi8 S microscope equipped with an sCMOS camera (Leica DFC9000 GTC), captured as z-stack images (z-step size: 0.18 μm) using a PlanAPO 63× (1.47 NA) oil-immersion objective, and analyzed with ImageJ software. Surface expression was quantified as the ratio of surface-to-intracellular fluorescence signals (Kuchtiak et al., 2024).

### Analysis of surface expression by fluorescence microscopy in neurons

The analysis of surface expression in neurons was performed as previously described (Kysilov et al., 2024) (Candelas Serra et al., 2024) (Kuchtiak et al., 2024). At 4 days *in vitro* (DIV), lentiviral particles encoding GluN2B-GFP or GluN2B^Δ^-GFP subunits were introduced into primary neuronal cultures. The surface and intracellular expression of GluN2B-GFP subunits was assessed at 14 DIV, following a protocol similar to that used for HEK293T cells, with minor modifications. Antibodies were diluted to 1:500, and intracellular GluN2B staining was performed using a goat anti-rabbit Alexa Fluor 488 secondary antibody (1:500; AA-11034; Invitrogen). Synapses were labeled using a primary mouse anti-postsynaptic density protein 95 (PSD-95) antibody (1:500; K28/43; NeuroMab), followed by a goat anti-mouse Alexa Fluor 555 secondary antibody (1:500; A-21422; Invitrogen). Images were acquired and analyzed as described for the HEK293T cell surface expression analysis, with an additional deconvolution step performed using Huygens software with the Classic Maximum Likelihood Estimation algorithm.

#### Analysis of endogenous NMDAR surface expression in neurons by fluorescence microscopy

At DIV 14, surface GluN2A and GluN2B subunits were labeled in primary hippocampal neurons from *Grin2b*^+/+^ or *Grin2b*^+/Δ^ mice by live staining with a rabbit anti-GluN2A antibody (1:250; AGC-002; Alomone) or a rabbit anti-GluN2B (1:250; AGC-003; Alomone), both targeting extracellular epitopes. Antibodies were diluted in Neurobasal medium (Gibco™), and cells were incubated at room temperature for 30 min. Following incubation, neurons were washed with PBS, fixed in 4% PFA, and blocked using a surface antigen blocking solution (PBS containing 50 mM NH_4_Cl, 10% goat serum, and 2% fish gelatin). Cells were then incubated for 1 h with goat anti-rabbit Alexa Fluor 488 secondary antibody (1:500; AA-11034; Invitrogen). After mounting, images were acquired as described for HEK293T cells. Image analysis was performed using ImageJ with a custom macro. Following maximum z-stack projection, a 20 μm dendritic region of interest (ROI) was selected and duplicated. The ROI was then thresholded, and receptor puncta were identified using Analyze Particles (ImageJ plugin). The mean fluorescence intensity of receptor clusters was measured within the 20 μm dendrite after projection.

### Primary hippocampal microisland cultures

For electrophysiological experiments primary hippocampal microisland cultures were used and prepared as previously described (Smejkalova et al., 2021). Glass coverslips (24 mm; Glaswarenfabrik Karl Hecht) were coated with 0.15% agarose (Serva) and allowed to dry. Microdots (∼200 µm in diameter, spaced ∼200 µm apart) were stamped using a growth-permissive substrate containing 0.1 mg/ml poly-D-lysine (Sigma) and 0.2 mg/ml collagen (Serva). Following decapitation, cortices and hippocampi were dissected from postnatal day 0–1 (P0–P1) mouse pups in Hank’s Balanced Salt Solution (HBSS; Gibco/Invitrogen). Primary cortical astrocytes were prepared from wild-type mice, dissociated using trypsin (Gibco/Invitrogen) and mechanical trituration, and cultured for two weeks in Dulbecco’s Modified Eagle Medium (DMEM) with GlutaMAX (Gibco/Invitrogen), supplemented with 10% fetal bovine serum (FBS; Gibco/Invitrogen) and 1% penicillin-streptomycin (Sigma), at 37°C in a humidified atmosphere containing 5% CO₂. Two days before hippocampal dissection, astrocytes were trypsinized and replated onto the microdot-stamped coverslips at a density of ∼6000 cells/cm^2^. Primary hippocampal neurons were then prepared from pups obtained from *Grin2b*^+/+^ and *Grin2b*^+/Δ^ crossings. Hippocampi from each pup (P0–P1; male and female) were processed individually. A sample of brain tissue was collected and stored at -20°C for subsequent genotyping. Hippocampi were digested using papain (Worthington) at 37°C for 40 min, followed by mechanical dissociation. Neurons were plated at a density of ∼300 cells/cm² on top of the astrocyte microislands in Neurobasal A medium supplemented with B27 (Gibco/Invitrogen), GlutaMAX (Gibco/Invitrogen), and penicillin-streptomycin (Sigma). Microisland cultures were maintained at 37°C in a humidified 5% CO₂ incubator until used for electrophysiological recordings.

### Electrophysiology

Whole-cell voltage-clamp recordings were obtained from one-neuron hippocampal microisland cultures (7–21 DIV) at -70 mV using an Axopatch 200B amplifier (Axon Instruments/Molecular Devices). Data were sampled at 10 kHz and filtered at 2 kHz. Series resistance was compensated by 70–80%. Recordings were excluded if series resistance exceeded 20 MΩ or increased by more than 20% during the session. Patch electrodes (4–6 MΩ) pulled from borosilicate glass capillaries were filled with intracellular solution containing (in mM): 125 K^+^ gluconate, 15 KCl, 10 HEPES, 5 EGTA, 0.5 CaCl_2_, 2 ATP Mg salt, 0.3 GTP Na salt, and 10 creatine phosphate, with pH adjusted to 7.2 using KOH. For agonist-evoked whole-cell responses, the extracellular solution contained (in mM): 160 NaCl, 2.5 KCl, 10 HEPES, 10 glucose, 0.2 EDTA, and 0.7 CaCl_2_, with pH adjusted to 7.3 using NaOH. Kainate (100 µM) or NMDA (100 µM) were applied to evoke AMPAR or NMDAR currents, respectively, while tetrodotoxin (TTX, 0.5 µM) was used to block action potentials.

Miniature excitatory postsynaptic currents (mEPSCs) were recorded in extracellular solution containing (in mM): 160 NaCl, 2.5 KCl, 10 glucose, 10 HEPES, 2 CaCl_2_, and 1 MgCl_2_, with pH adjusted to 7.3 using NaOH. Evoked EPSCs (eEPSCs) were elicited by brief depolarization (1 ms at +10 mV) from the holding potential of -70 mV every 10 s and recorded in Mg²⁺-free extracellular solution supplemented with 10 µM glycine. All extracellular solutions contained bicuculline methochloride (10 µM) to block GABA_A_ receptors. To pharmacologically isolate NMDAR-or AMPAR-EPSCs, 2,3-dioxo-6-nitro-1,2,3,4-tetrahydrobenzo[f]quinoxaline-7-sulfonamide (NBQX, 3 µM) or AP-5 (50 µM) were used, respectively. Ifenprodil (3 µM) was used to inhibit responses mediated by NMDARs containing GluN2B subunits. To assess short-term plasticity, paired-pulse ratio (PPR) was calculated from AMPAR-EPSCs by dividing the peak amplitude of the second eEPSC by that of the first, using an interpulse interval of 50 ms. The amplitudes, the PPR, and the deactivation kinetics of eEPSCs were analyzed from 4–5 consecutive individual sweeps and the obtained values were averaged. Data were analyzed with respect to the genotype and the sex of the animals to examine potential sex differences in the studied parameters.

### Animal behavioral tests

Behavioral testing of *Grin2b*^+/Δ^ animals and their *Grin2b*^+/+^ littermates took place at 9–15 weeks of age, with males and females tested separately on different days. The analyses were conducted in the following sequence: open field test, elevated plus maze, novel object recognition, social preference test, nestlet shredding, prepulse inhibition of the acoustic startle (PPI) test, and contextual and cued fear conditioning. A separate cohort of animals was tested in the IntelliCage system. The details of the open field test, the PPI test, contextual and cued fear conditioning, and the IntelliCage system have been previously described (Candelas Serra et al., 2024).

#### Open field test

Each mouse was placed in the corner of a cubic (42 x 42 cm) uniformly illuminated (200 lux) open-top box and allowed 10 min of free exploration. Mouse movement was monitored and analyzed using a video tracking system (Viewer software, Biobserven GmbH, Germany).

#### Prepulse inhibition test

Animals were acclimated to soundproof cabinets (Med Associates Inc., USA) for 10 min before the start of the test. Four prepulse acoustic stimuli (70, 77, 82, 85 dB; 20 ms duration) were used alone and in combination with a startle stimulus (110 dB, 50 ms duration). White noise was used as the stimulus, and all delivery and recordings were made against a constant background noise of 65 dB. Ten trials of startle, prepulse, and combined prepulse with startle were performed in a randomized order, with intertrial intervals randomly varied between 10–20 s. All animal responses to the stimuli were detected, quantified, graphically represented, and automatically stored using a sensor platform, amplifier, and computer software.

#### Cued and contextual fear conditioning test

Mice were placed in a conditioning cage inside a soundproof cabinet (Ugo Basile, Gemonio, Italy) with a stainless-steel rod floor for shock delivery. The test starts with an adaptation period followed by two pairings of a conditioned stimulus (20 s of 4 kHz pure tone at 77 dB) and the unconditioned stimulus (1 s of 0.5 mA constant current to the cage floor). The unconditioned stimulus coincided with the cessation of the conditioned stimulus, with a 2-min interval between pairings. Contextual memory was assessed 24 h after learning in the same environment, with freezing behavior automatically recorded during a 6-min trial using ANY-maze software (Ugo Basile, Gemonio, Italy). For the delayed cued memory test, mice were introduced to a novel context 3 h after the contextual trial. The freezing response to the conditioned stimulus was monitored for 2 min.

#### IntelliCage tests

IntelliCage system is a fully automated device that monitors the cognitive abilities and emotional states of group-housed laboratory mice. One week before transferring to the IntelliCage, all animals received subcutaneous transponders (ISO-compliant). The IntelliCages were equipped with four operant corners, each containing two door-guided openings for drinking bottles. Each corner was equipped with a radiofrequency identification antenna to detect implanted transponders, allowing identification of animals upon entry. Access to water was contingent upon correct nose pokes at the door of the selected corner. Animals were tested in several phases, with water availability in specific corners regulated to assess spatial learning, spatial reversal learning, learning extinction, and working memory (Candelas Serra et al., 2024).

#### Elevated plus maze test

The elevated plus maze apparatus consisted of two closed arms and two open arms, with a light intensity of 60 lux at the center of the maze. Animals were placed in the center and allowed to explore the maze for 5 min. The total time spent in the open arms, closed arms, and the center was automatically recorded and analyzed using the Viewer software (Biobserve GmbH, Germany).

#### Novel object recognition test

The novel object recognition test was employed to evaluate exploratory behavior towards a novel object as a measure of recognition memory. This fully automated test utilized a video tracking system (Viewer, Biobserve GmbH, Bonn, Germany). During a two-day habituation phase the animals were allowed to explore the testing arena for 10 min each day. On the third day, the animals were exposed to two identical objects during a familiarization phase. Following a 3-h retention interval, a probe trial was conducted in which one of the familiar objects was replaced with a novel object. All trials were performed under the illumination of 70 lux at the center of the maze. Objects were placed 7.5 cm from the walls, with their positions alternating across trials to control for side preference. Test subjects were consistently introduced from the wall opposite the object locations to maintain positional balance. The system automatically recorded object visits, exploration time for each object, and the corresponding percentages.

#### Social preference test

The social preference test was performed using two cylindrical wire cages placed in the open-field arena. Each cage (10.5 cm in diameter, 15 cm in height) was positioned equidistantly from two diagonally opposite corners of the arena, 10 cm from the walls. All procedures were conducted under low-light conditions (70 lux at the center of the open field). The experimental and juvenile target animals were sex-matched. Target mice were pre-habituated to the empty wire cages in the open field for 10 min on the day before testing. On the test day, the target animal was placed in one of the wire cages for 10 min without the experimental mouse present; a Duplo block was placed in the other as a non-social control. The experimental mouse was then introduced into the open field and allowed to explore both cages freely for 10 min. Latency to approach, number of visits, and interaction time with the cage containing the target animal were automatically recorded and analyzed using the Viewer software (Biobserve GmbH, Germany).

#### Nestlet shredding

Mice were housed individually for one week for acclimation. Following this period, 12 g of nesting material (Animalab) was provided in each cage for five consecutive days. To evaluate repetitive nestlet shredding behavior, the unused nesting material was collected and weighed at the same time each day to determine the amount utilized by the mice.

### Statistical analysis

Data are presented in bar graphs showing mean ± standard error of the mean (SEM), or in scatter plots showing the distribution of individual cells overlaid on bar graphs showing mean ± SEM. Unless specified otherwise, the sample size *n* refers to the number of cells or animals (independent values) per group. Statistical analysis was performed with STATGRAPHICS Centurion 18 (Statgraphics Technologies, Inc., USA). If the original data did not have a symmetric distribution and constant variance, they were power transformed to attain symmetric distribution and homoscedasticity (constant variance). We used absolute values of studentized residuals greater than 3 to identify outliers. Data were further analyzed by Student’s t-test or the Analysis of variance (ANOVA) to compare more than two groups, using the General Linear Model function in STATGRAPHICS. When statistically significant differences were found, *post-hoc* analyses were conducted using the Dunnett method (for comparisons to the control), the Duncan method (for all pairwise comparisons), or the Least Significant Difference (LSD) method (for selected pairwise comparisons). A *p-value* ≤ 0.05 was considered statistically significant throughout the study. For experiments where data were collected with respect to cell age *in vitro*, we divided the DIV into four age groups with approximately the same number of data points in each group, to be able to see the trend with respect to age. Afterwards, we analyzed these groups using the ANOVA (using the General Linear Model function in STATGRAPHICS).

## Results

### Generation of mice with a protein-truncating variant in the *Grin2b* gene

Understanding the interplay between genetic variation and possible compensatory mechanisms is critical for developing targeted interventions for *GRIN*-related disorders (Quatraccioni et al., 2024). Recently, we have generated a knock-in mouse strain carrying a patient missense variant in the *Grin2b* gene (*Grin2b*^+/L825V^) to explore the consequences of this variant for NMDAR function, synaptic transmission, and behavior (Candelas Serra et al., 2024). A male patient heterozygous for the GluN2B-L825V variant presented with severe ID and ASD (Awadalla et al., 2010) (Tarabeux et al., 2011). We have also generated a mouse line with a frameshift variant introduced at the same location in *Grin2b*, leading to a premature termination codon (*Grin2b*^+/Δ^) (Fig. 1A). In the process of CRISPR-Cas9 genome editing, four thymidines were inserted in the *Grin2b* gene at the triplet encoding L825. This leads to a leucine to phenylalanine substitution and a frameshift, altering the sequence of the subsequent 15 amino acids and introducing a stop codon (GluN2B-L825Ffs*15). The mutation is predicted to produce a truncated GluN2B subunit lacking part of the M4 membrane domain and the entire CTD (GluN2B^Δ^; Fig. 1A-B; see Fig. 1C for Sanger validation of the GluN2B-L825Ffs*15 mutation in *Grin2b*⁺^/Δ^ and *Grin2b*^Δ/Δ^ mice and confirmation of its absence in *Grin2b*^+/+^ mice). A female patient heterozygous for a PTV GluN2B-E839* with a premature termination codon at the same position as in the GluN2B-L825Ffs*15 variant exhibits mild/moderate ID, fine motor skill deficits, abnormal EEG, sleep problems, and early puberty (Dr. Xavier Altafaj, Fac. de Medicina i Ciències de la Salut, Barcelona, personal communication) (Santos-Gomez et al., 2021). To elucidate possible consequences of a PTV at this location we decided to examine the cellular and neurobehavioral effects of the GluN2B-L825Ffs*15 PTV in our mouse model and assess potential compensatory changes at the mRNA and protein levels.

**Figure 1.**
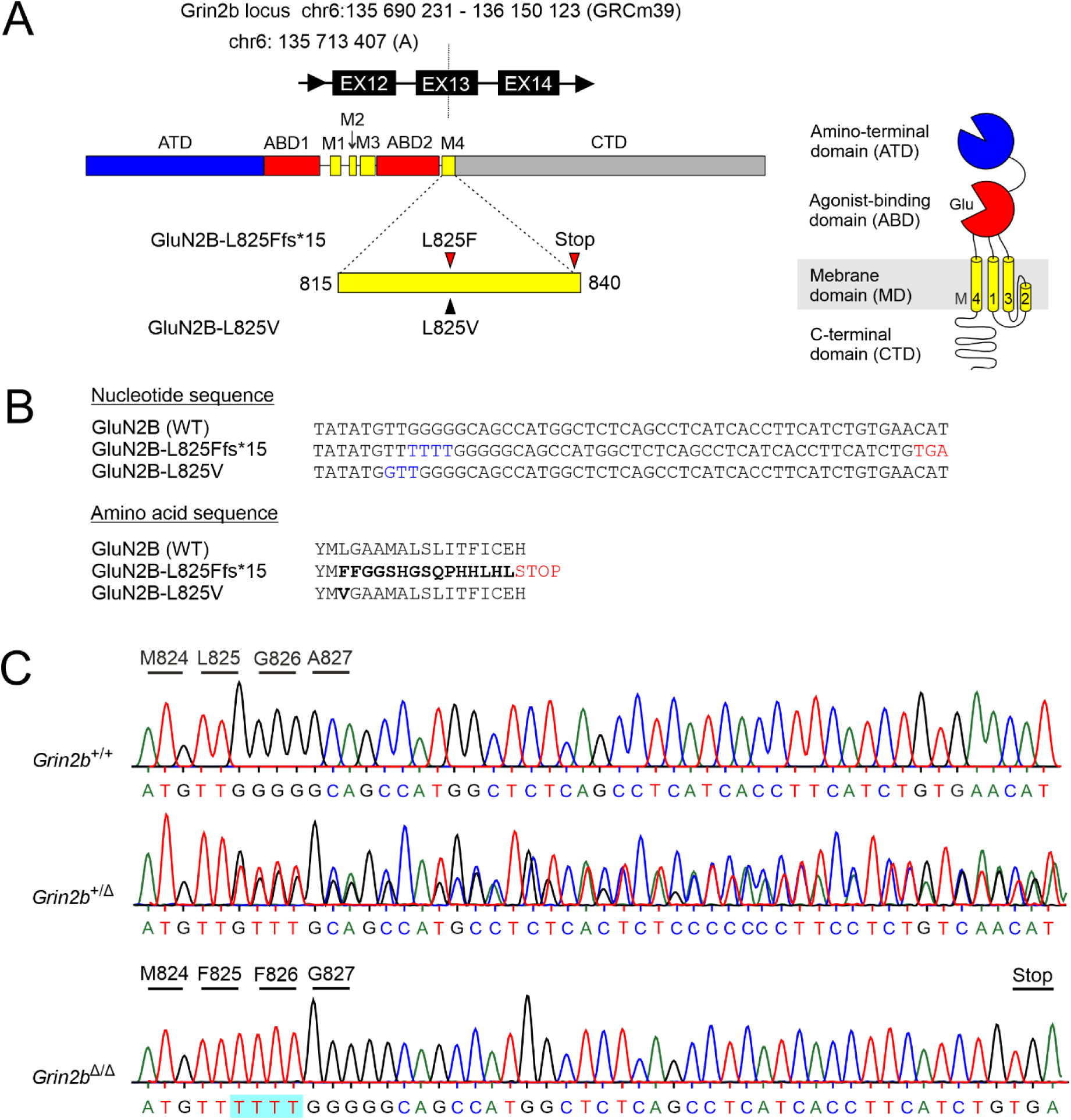
Generation of *Grin2b*⁺^/Δ^ mice using CRISPR-Cas9. (A) Schematic representation of the GluN2B subunit structure, showing the extracellular amino-terminal domain (ATD); agonist-binding domain (ABD; comprising S1 and S2 lobes), the transmembrane domain (TMD; consisting of M1, M3, M4, and the re-entrant M2 loop), and the intracellular CTD. The M4 segment contains the target site used to generate both the GluN2B^Δ^ and the GluN2B-L825V variants. (B) *Top:* Nucleotide sequences of the wild-type and the two *Grin2b* variants. Note the insertion of four thymidines (blue) resulting in a frameshift and a premature termination codon (TGA; red), corresponding to the GluN2B-L825Ffs*15 variant. *Bottom:* Amino acid sequences of the wild-type and the two GluN2B subunit variants. (C) Chromatograms represent the Sanger sequencing validation of the GluN2B-L825Ffs*15 variant in heterozygous (*Grin2b*⁺^/Δ^) and homozygous (*Grin2b*^Δ/Δ^) mice, with confirmation of its absence in wild-type (*Grin2b*^+/+^) controls.

Homozygous *Grin2b*^Δ/Δ^ mice, similar to mice with loss-of-function missense mutations *Grin2b*^C456Y/C456Y^ or *Grin2b*^L825V/L825V^ (Shin et al., 2020) (Candelas Serra et al., 2024) die shortly after birth. Our heterozygous mutant mice (*Grin2b*^+/Δ^) obtained from breeding *Grin2b*^+/Δ^ with wild-type mice were born at the expected Mendelian ratios and exhibited normal survival and growth. No differences in weight were observed when adult *Grin2b*^+/Δ^ and *Grin2b*^+/+^ mice were compared (*Grin2b*^+/Δ^ *vs. Grin2b*^+/+^; male: 24.4 ± 0.59 g, *n* = 11 *vs.* 25.4 ± 0.97 g, *n* = 10 (*p* = 0.48; Student’s t-test); female: 20.1 ± 0.30 g, *n* = 10 *vs.* 20.4 ± 0.39 g, *n* = 12 (*p* = 0.64; Student’s t-test).

### Expression of *Grin* mRNA in the hippocampus of *Grin2b*^+/Δ^ mice

Organisms have developed genetic robustness, defined as the ability to minimize the effects of pathogenic nonsense variants at the phenotypic level (de Visser et al., 2003). First we proceeded to investigate whether the monoallelic PTV in *Grin2b*^+/Δ^ mice could introduce compensatory mechanisms resulting in transcriptional downregulation of mutant mRNA and the upregulation of homologous genes (Howe and Patani, 2023). The mRNA expression levels of NMDAR subunits were determined in the hippocampus of adult male and female *Grin2b*^+/+^ and *Grin2b*^+/Δ^ mice by RT-qPCR. In male *Grin2b*^+/Δ^ mice, *Grin1* and *Grin2b* mRNA levels were significantly lower by 20.1 ± 2.5% (*p* = 0.017; Student’s t-test) and 5.8 ± 2.3% (*p* = 0.042; Student’s t-test), respectively, compared to samples from male *Grin2b*^+/+^ controls, while *Grin2a* expression remained unchanged (*p* = 0.964; Student’s t-test) (Fig. 2A). In female *Grin2b*^+/Δ^ mice, *Grin1* mRNA levels tended to be lower by 17.8 ± 6.4% (p = 0.079; Student’s t-test) and *Grin2b* mRNA levels were significantly reduced by 19.3 ± 4.8% (*p* = 0.004; Student’s *t*-test), compared to samples from female *Grin2b*^+/+^ controls, while *Grin2a* mRNA expression was unchanged (p = 0.552; Student’s t-test), (Fig. 2B).

**Figure 2.**
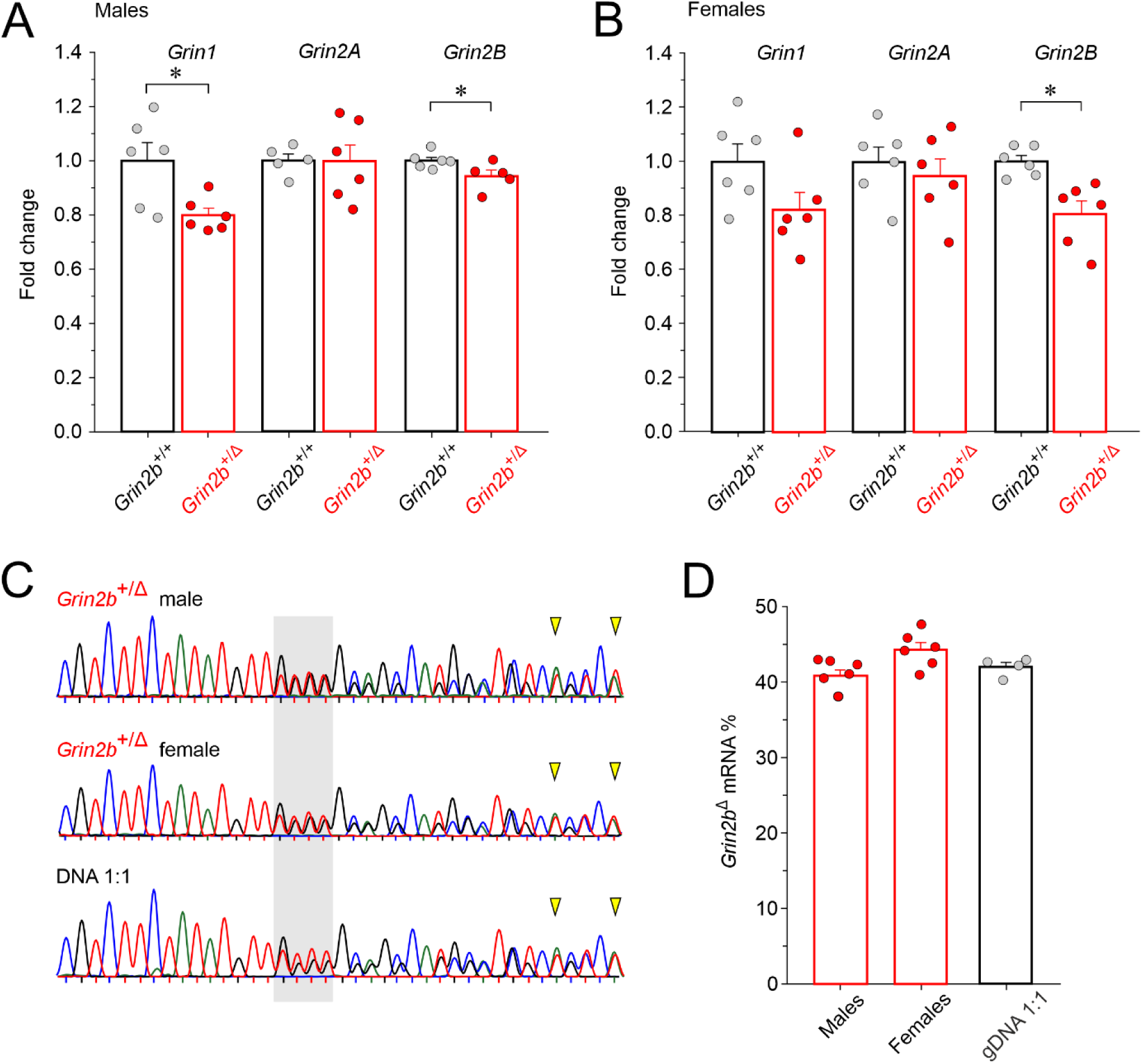
The effect of the *Grin2b*^Δ^ mutation on *Grin1*, *Grin2a,* and *Grin2b* mRNA expression. Relative mRNA expression levels of *Grin1*, *Grin2a*, and *Grin2b* in the hippocampus of adult male (A) and female (B) *Grin2b*^+/+^ and *Grin2b*⁺^/Δ^ mice, as determined by RT-qPCR. Data were normalized to the corresponding mean *Grin2b*^+/+^ value and are presented as mean ± SEM and analyzed using Student’s t-test to compare the expression in *Grin2b*⁺^/Δ^ versus *Grin2b*^+/+^ mice; asterisks (*) indicate statistically significant differences. (C) Representative chromatograms of Sanger sequencing of PCR amplicons spanning the region of the frameshift mutation (gray box) in male and female *Grin2b*⁺^/Δ^ mice and the plasmid DNA mixture of *Grin2b*⁺:*Grin2b*^Δ^ at a 1:1 ratio; yellow arrowheads indicate two of the five analyzed nucleotides. (D) The relative expression levels of *Grin2b* mRNA show comparable expression of wild-type and mutant alleles. Data are presented as mean ± SEM and were analyzed using one-way ANOVA (no significant differences were found).

It is important to note that *Grin2b* mRNA levels in *Grin2b*^+/Δ^ mice reflect transcripts from both the wild-type and the mutant allele. To determine their relative expression, we performed classical end-point PCR using primers spanning multiple exon–exon boundaries to exclude genomic DNA (agarose gel of RT-PCR 1200 nt amplicons from hippocampal RNA using primers spanning exon 7 to 13 – see Table S1 for primer sequences), followed by direct sequencing of the resulting amplicons. Chromatogram analysis revealed that the peak amplitude ratio of five different nucleotides downstream of the GluN2B-L825Ffs*15 mutation (selected because they are not surrounded by identical nucleotides) was 40.8 ± 0.8% in males (*n* = 7) and 44.3 ± 1.0% in females (*n* = 6), indicating near-equal presence of the mutant and wild-type alleles (Fig. 2C, D). These ratios were not significantly different from control amplicons generated from a 1:1 gDNA mixture of *Grin2b*^+^ and *Grin2b*^Δ^ alleles (42.0 ± 0.6%, *n* = 4; male *p* = 0.328; female *p* = 0.121). This ratio between wild type and variant mRNA indicates that the variant mRNA does not undergo extensive degradation. This suggests that nonsense-mediated decay (NMD)—a mechanism that typically eliminates faulty mRNA—is not fully engaged in this case (Howe and Patani, 2023).

### Protein expression in the hippocampus of *Grin2b*^+/Δ^ mice

To assess the impact of the *Grin2b*^Δ^ mutation on the brain proteome, we performed mass spectrometry-based proteomic analysis on hippocampi from adult male *Grin2b*^+/+^ (*n* = 4) and *Grin2b*⁺^/Δ^ (*n* = 5) mice (see Methods). After applying filtering criteria, we identified 5,454 proteins for further analysis. The presence of the *Grin2b*^Δ^ allele in *Grin2b*⁺^/Δ^ mice significantly altered the expression of 23 proteins (4 down-regulated and 19 up-regulated), each showing more than a twofold change compared to *Grin2b*^+/+^ controls (Fig. 3A). Among these, three up-regulated proteins were neuron specific: ACTL6B (implicated in neurodevelopmental deficits), BACE1 (involved in amyloid precursor protein processing), and BAX (essential for developmental neuronal apoptosis). The remaining proteins lacked a clear link to brain function, including SEC62 (endoplasmic reticulum (ER)-associated protein), VMP1 (involved in zymogen granule degradation in pancreatitis), ARMH3 (Golgi organization regulator), H2BU2 (nucleosome component), MT1 (metal homeostasis and detoxification), and GAS8 (structural component of ciliary movement) (Fig. 3A; Table S2 lists proteins with altered expression levels, the corresponding gene names, and the protein biological functions). Notably, several mildly dysregulated proteins were associated with tumorigenesis.

**Figure 3.**
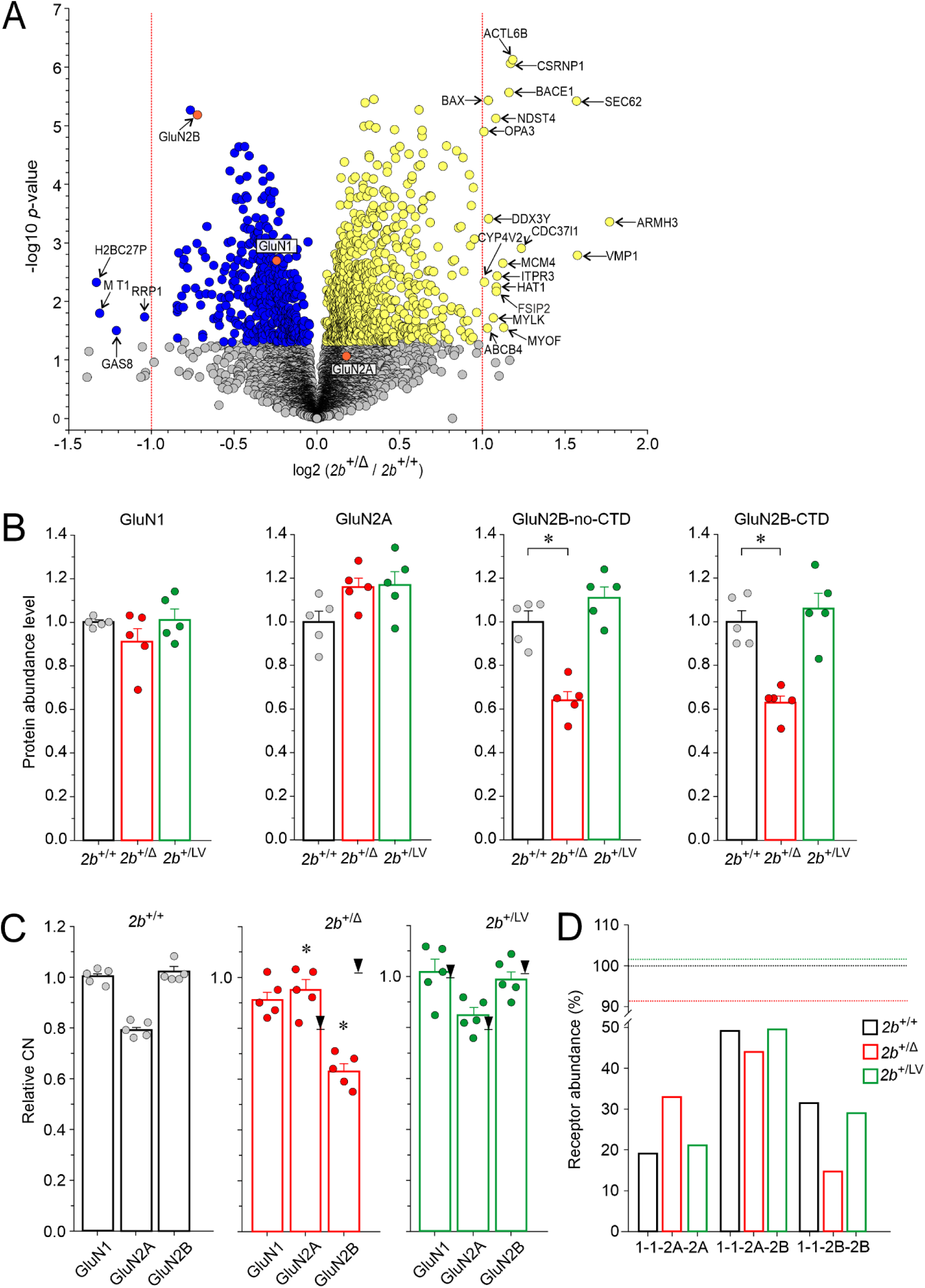
Protein expression analysis in the hippocampus of *Grin2b*^+/+^ and *Grin2b*⁺^/Δ^ mice. (A) Volcano plot of proteomics analysis comparing protein levels in hippocampal tissue from adult *Grin2b*^+/+^ and *Grin2b*⁺^/Δ^ mice. The x-axis shows log₂ fold-change (negative: downregulated; positive: upregulated), and the y-axis shows –log₁₀ p-values. The horizontal change of color marks the *p* = 0.05 significance threshold, and vertical dashed lines indicate a two-fold change. Proteins significantly downregulated in *Grin2b*⁺^/Δ^ mice are shown in blue; significantly upregulated proteins are shown in yellow; non-significant changes are in gray. Statistical significance was determined using a two-sample test (see Methods). (B) Protein abundance levels of GluN1, GluN2A, GluN2B-no-CTD, and GluN2B-CTD in *Grin2b*^+/+^ (*2b*^+/+^), *Grin2b*⁺^/Δ^ (*2b*⁺^/Δ^), and *Grin2b*⁺^/L825V^ (*2b*⁺^/LV^) mice, normalized to the wild-type *Grin2b*^+/+^ reference. Data are presented as mean ± SEM; asterisks (*) indicate statistically significant differences (one-way ANOVA followed by the Dunnett method). (C) Relative CN calculated as the CN of GluN1, GluN2A, and GluN2B subunits in *Grin2b*^+/+^, *Grin2b*⁺^/Δ^, and *Grin2b*⁺^/L825V^ mice, normalized to the CN of GluN1 subunit in *Grin2b*^+/+^ mice. Arrows indicate the reference GluN subunit levels in *Grin2b*^+/+^ mice. Data are presented as mean ± SEM; asterisks (*) indicate significant differences in CN between genotypes (one-way ANOVA followed by the Dunnett method). (D) Estimated relative abundance of GluN1/GluN2B diheteromers (1-1-2B-2B), GluN1/GluN2A/GluN2B triheteromers (1-1-2A-2B), and GluN1/GluN2A diheteromers (1-1-2A-2A) in *Grin2b*^+/+^, *Grin2b*⁺^/Δ^, and *Grin2b*⁺^/L825V^ mice, based on the CN data in (C) and calculation methods described Methods. Dotted lines indicate total NMDAR content.

Since disease-associated GluN2B variants have been shown to impact embryonic and early postnatal neurodevelopment, including neuron differentiation, dendritogenesis, synaptogenesis, and synaptic plasticity (Sabo et al., 2022) we sought to determine whether the presence of the *Grin2b*^Δ^ allele in *Grin2b*⁺^/Δ^ mice may affect the expression of specific synaptic proteins in the hippocampal tissue. The results of the analysis showed that the expression levels of proteins involved in vesicle exocytosis and synaptic function were not significantly altered in the hippocampus of *Grin2b*⁺^/Δ^ mice compared to their *Grin2b*^+/+^ siblings (*p* = 0.767; Student’s t-test). The average ratio for the analyzed synaptic proteins in *Grin2b*⁺^/Δ^ mice relative to *Grin2b*^+/+^ controls was 1.08 ± 0.016 (see Table S3 for gene and protein names of the selected synaptic proteins, along with their ratios in *Grin2b*^+/Δ^ *vs*. *Grin2b*^+/+^ hippocampal tissue), suggesting that the development of synaptic contacts in the hippocampus remains largely unchanged in *Grin2b*^+/Δ^ mice.

To assess potential changes in the relative expression of NMDAR subunits, we quantified the protein abundance levels of GluN subunits in *Grin2b*^+/Δ^ mice, normalizing them to the *Grin2b*^+/+^ reference level (Fig. 3B). Protein abundance levels of GluN1 and GluN2A were quantified using total peptide signal intensities spanning the entire protein and did not differ significantly between *Grin2b*^+/Δ^ and *Grin2b*^+/+^ mice. For GluN2B, two separate protein abundance measurements were obtained: (*i.*) GluN2B-no-CTD, determined by the total intensity of unique GluN2B peptides from the N-terminal domain to Asp768, representing the combined abundance of both wild-type and GluN2B^Δ^ subunits and (*ii.*) GluN2B-CTD, determined by the total intensity of unique CTD peptides, reflecting only the wild-type GluN2B subunit. The protein abundance level of GluN2B-no-CTD was significantly reduced in *Grin2b*^+/Δ^ mice (0.64 ± 0.04-fold; *n* = 5) compared to *Grin2b*^+/+^ (Fig. 3B). Similarly, the GluN2B-CTD abundance was significantly reduced in *Grin2b*^+/Δ^ mice (0.63 ± 0.03-fold; *n* = 5) compared to *Grin2b*^+/+^. The small difference (1.0%) between GluN2B-CTD and GluN2B-no-CTD abundance suggests that the truncated GluN2B^Δ^ subunit protein is present at negligible levels in the proteome (Fig. 3B).

To determine whether the observed changes in the abundance of the GluN2B subunit were specific to the frameshift variant, we performed proteomic analysis on hippocampi from heterozygous mice carrying the missense GluN2B-L825V mutation at a similar site (*Grin2b*^+/L825V^) (see (Candelas Serra et al., 2024), for electrophysiological, molecular, and behavioral characterization). The protein abundance levels of GluN1, GluN2A, GluN2B-no-CTD, and GluN2B-CTD in *Grin2b*^+/L825V^ mice did not significantly differ from the *Grin2b*^+/+^ reference levels (Fig. 3B), indicating that, unlike the frameshift *Grin2b*^Δ^ allele, the missense *Grin2b*^L825V^ allele likely produces significant quantities of the protein product.

### Possible consequences of the *Grin2b*^Δ^ allele on the NMDAR subunit composition

In *Grin2b*^+/+^ mice, the mean relative CN (see Methods) of GluN2A and GluN2B subunits was 0.79 and 1.02, respectively, of the CN of the GluN1 subunit (Fig. 3C). Given the tetrameric structure of NMDARs—containing two GluN1 and two GluN2 subunits—it may be the abundance of GluN1, rather than the abundance of GluN2A/B subunits, that primarily determines the total number of NMDARs in the adult hippocampus. The relative CN of NMDAR subunits allows for an estimation of the distribution of diheteromeric and triheteromeric NMDARs, given several assumptions: First, NMDARs consist of four subunits: two GluN1 and either two GluN2A, two GluN2B (diheteromeric receptors), or one of each (triheteromeric receptors) (Al-Hallaq et al., 2007) (Luo et al., 1997) (Tovar et al., 2013) (Rauner and Kohr, 2011). Second, GluN1 subunits randomly associate with GluN2 subunits with equal preference. Third, the majority of the available subunits assemble into functional NMDARs (Luo et al., 1997; Luo et al., 2002). In *Grin2b*^+/Δ^ mice, assuming all GluN1 subunits are incorporated into NMDARs, together 44.8% of GluN2A and GluN2B subunits remain unassembled. The estimated relative distribution of GluN1/GluN2A, GluN1/GluN2B, and GluN1/GluN2A/GluN2B receptors in *Grin2b*^+/+^ mice is 19.2%, 31.5%, and 49.2%, respectively (Fig. 3D).

In the hippocampus of *Grin2b*^+/Δ^ mice, the mean relative CN of GluN1, GluN2A, and GluN2B subunits was 0.91, 0.95, and 0.63, respectively, of the relative CN of the GluN1 subunit in *Grin2b*^+/+^ mice (Fig. 3C). Based on protein abundance estimates, the GluN2B signal predominantly represents the wild-type GluN2B subunit (∼99%), with only a ∼1% contribution of GluN2B^Δ^. In *Grin2b*^+/Δ^ mice, again assuming all the available GluN1 subunits are incorporated into NMDARs, 42.2% of GluN2A and GluN2B subunits remain unassembled. Calculations indicate an 8.6% reduction in the total number of NMDARs in *Grin2b*^+/Δ^ mice compared to *Grin2b*^+/+^ controls. Notably, this reduction is likely driven by a small decline in GluN1 levels rather than a decrease in GluN2B. On the other hand, the distribution of diheteromeric and triheteromeric NMDARs was affected by the relative increase in GluN2A and decrease in GluN2B levels in *Grin2b*^+/Δ^ mice compared to *Grin2b*^+/+^ controls. In *Grin2b*^+/Δ^ mice, the estimated relative distribution of diheteromeric GluN1/GluN2A, GluN1/GluN2B, and triheteromeric GluN1/GluN2A/GluN2B receptors was 32.9%, 14.6%, and 43.9%, respectively (Fig. 3D).

Next, we analyzed the CN of NMDAR subunits in *Grin2b*^+/L825V^ mice (Candelas Serra et al., 2024). In these mice, the relative CN of GluN1, GluN2A, and GluN2B were 1.02, 0.85, and 0.99, respectively, of the CN of the GluN1 subunit in *Grin2b*^+/+^ mice (Fig. 3C). Since proteomic analysis cannot distinguish between wild-type GluN2B and the GluN2B-L825V variant, the value reflects the combined abundance of both wild-type and variant subunits. Minor CN variations in *Grin2b*^+/L825V^ resulted in modest changes in the total number of NMDARs and the relative distribution diheteromeric of GluN1/GluN2A, GluN1/GluN2B, and triheteromeric GluN1/GluN2A/GluN2B receptors compared to *Grin2b*^+/+^ mice (Fig. 3D). Notably, while all NMDARs in *Grin2b*^+/Δ^ mice should exhibit normal functional properties, a subset of NMDARs in *Grin2b*^+/L825V^ mice is expected to have reduced function due to a reduced probability of opening of receptors containing the GluN2B-L825V subunit (Candelas Serra et al., 2024).

Together, these findings highlight the possible role of the GluN1 subunit availability in determining the total number of NMDARs in hippocampal neurons. Our results suggest that the reduced availability of the GluN2B subunit in *Grin2b*^+/Δ^ mice may result in an altered distribution of GluN2A-containing *vs.* GluN2B-containing NMDARs without affecting the overall NMDAR content.

### Surface expression of the GluN2B^Δ^ subunit

While our RT-qPCR results show no evidence of *Grin2b*^Δ^ mRNA degradation (Fig. 2D), our proteomic analysis strongly suggests that the truncated GluN2B^Δ^ subunit is virtually absent in the hippocampi of adult *Grin2b*⁺^/Δ^ mice (Fig. 3B). To better understand possible cellular fates of the GluN2B^Δ^ protein, we sought to determine whether NMDARs containing the GluN2B^Δ^ subunit would be able to assemble, reach the cell surface, and retain any functional activity. In addition we aimed to elucidate whether GluN2B^Δ^ subunit could affect surface expression of wild type GluN2B subunits. NMDARs containing the GluN2B subunit truncated before or within the M4 domain, or with mutations in the four amino acids following the M4 segment (HLFY(840-843)), exhibit defects in surface expression (Horak et al., 2008) (Hawkins et al., 2004; Kysilov et al., 2024). We performed patch-clamp recordings from GFP-labeled HEK293T cells transfected with GluN1 and GluN2B^Δ^ and found no detectable responses to 1 mM glutamate (0/10; response >10 pA was considered the minimum threshold), whereas all GFP-labeled HEK293T cells transfected with GluN1 and GluN2B exhibited responses to 1 mM glutamate (10/10; mean peak response -596 ± 134 pA). There are several possible explanations for the absence of responses in GluN2B^Δ^-transfected cells: GluN2B^Δ^ subunits may not be expressed, or GluN2B^Δ^ subunits may fail to assemble with GluN1, or NMDARs containing GluN2B^Δ^ subunits may assemble but are not trafficked to the cell surface, of the GluN2B^Δ^ subunit-containing NMDA receptors are non-functional.

To determine whether NMDARs containing the GluN2B^Δ^ subunit are expressed and trafficked to the cell surface, we performed SDS-PAGE followed by western blot analysis on HEK293T cells co-transfected with GluN1 and either wild-type GluN2B-GFP or GluN2B^Δ^-GFP. Figure 4A shows a western blot probed with anti-GFP to detect GluN2B subunits, and anti-GAPDH or anti-histone H3 antibodies to confirm appropriate protein enrichment in cytosolic (S1), cytoplasmic membrane (S2) and nuclear/organelle (P) fractions (see Methods). In the cytoplasmic membrane fraction (S2), a band at ∼194 kDa corresponding to wild-type GluN2B fused to GFP (167 + 27 kDa) was observed in HEK293T cells expressing GluN2B-GFP, although this band was most prominent in the nuclear/organelle (P) fraction. Similarly, a distinct band at ∼122 kDa corresponding to GluN2B^Δ^-GFP (95 + 27 kDa) was detected in HEK293T cells expressing the truncated subunit, suggesting that in the heterologous expression system significant amounts of the truncated subunit protein are produced. Since NMDARs are trafficked to the membrane only as assembled complexes (Wenthold et al., 2003), the presence of GluN2B^Δ^-GFP in the cytoplasmic membrane (S2) fraction supports the existence of NMDARs containing the truncated subunit and their trafficking to the cell surface in HEK293T cells.

**Figure 4.**
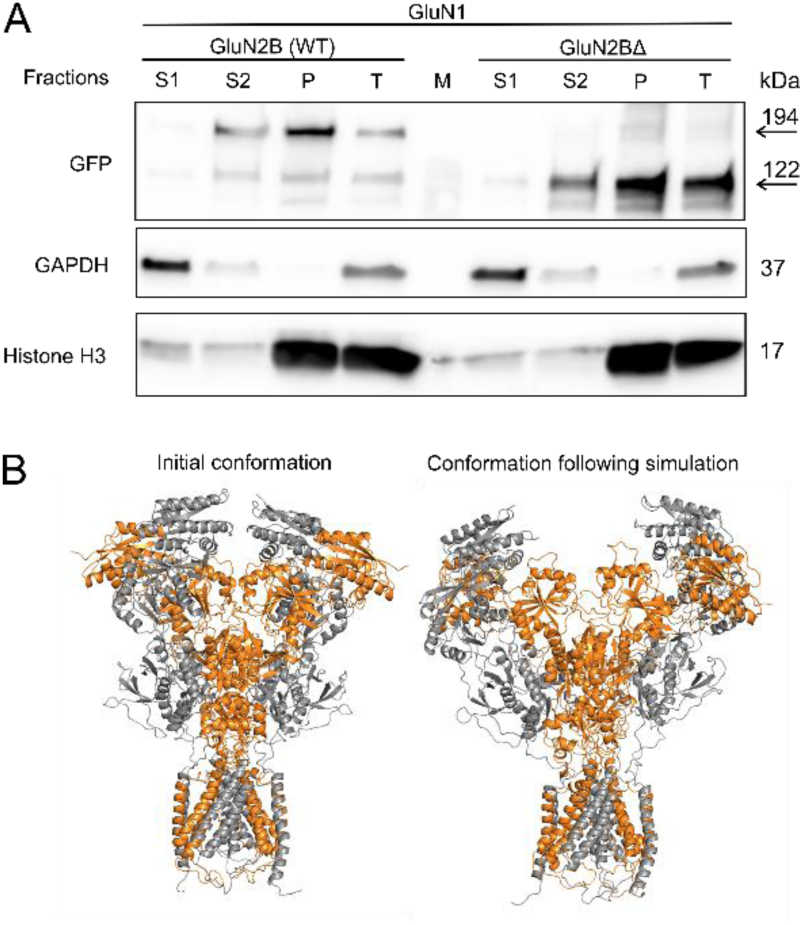
Subcellular localization and structure of GluN2B and GluN2B^Δ^. (A) Western blot analysis of cytosolic (S1), cytoplasmic membrane (S2), and nuclear/organelle (P) fractions prepared from HEK293T cells co-transfected with GluN1 and either wild-type GluN2B-GFP or mutant GluN2B^Δ^-GFP subunits. Membranes were probed with antibodies against GFP (to detect GFP-tagged GluN2B), GAPDH (cytosolic marker), and histone H3 (nuclear marker). The expected molecular weights are indicated on the right. (B) Structural model of the GluN1/GluN2B receptor with both GluN2B subunits truncated. The left panel shows the initial geometry and the right panel displays a snapshot after 500 ns of MD simulation. While the TMD remained relatively stable, the ATD and ABD regions exhibited substantial separation and loss of inter-domain contacts, resembling desensitized-like states.

Despite the truncation of the M4 helix and introduction of polar residues within the membrane bilayer, MD simulations of GluN1/GluN2B complexes carrying one or two truncated GluN2B subunits, suggest that the receptors remain stable and embedded in the model membrane. The interactions between the remaining transmembrane helices and the intact portion of M4 are preserved. During the 500 ns simulation—typically sufficient to capture TMD rearrangements—no significant structural changes associated with receptor closure were observed. In contrast, the ATD and ABD exhibited significant separation and loss of inter-domain contacts, potentially resembling desensitized states of related ionotropic glutamate receptors (Schauder et al., 2013). Structural snapshots of the GluN1/GluN2B receptor with both GluN2B subunits truncated at 500 ns are shown in Figure 4B.

To evaluate the impact of the GluN2B^Δ^ variant on NMDAR surface expression more quantitatively, we performed immunofluorescence microscopy in HEK293T cells transfected with equal amounts of expression vectors encoding GluN1 and either wild-type or mutant GFP-tagged GluN2B subunits. Compared to NMDARs containing GluN2B, receptors incorporating the GluN2B^Δ^ variant exhibited a 78% reduction in surface expression (*p* < 0.001, Student’s t-test) (Fig. 5A). This reduction was further confirmed by a colorimetric assay in COS-7 cells, which revealed a 90% decrease in surface expression (*p* < 0.001, Student’s t-test) (Fig. S2).

**Figure 5.**
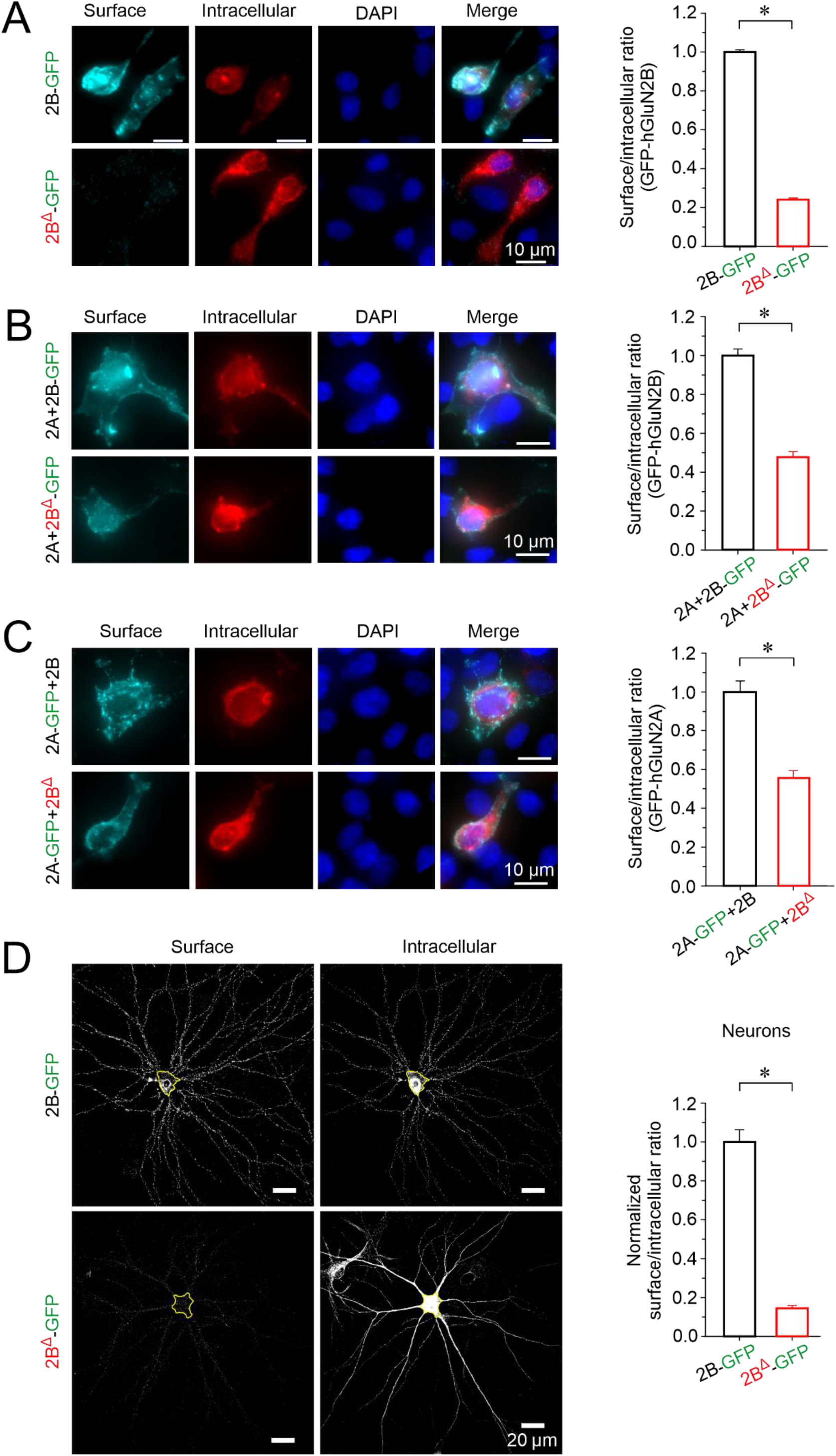
The GluN2B^Δ^ variant disrupts receptor cell surface expression. (A) *Left:* Representative immunofluorescence images showing surface and intracellular expression of GluN2B-GFP subunits in HEK293T cells co-transfected with GluN1 and either wild-type (2B-GFP) or mutant (2B^Δ^-GFP) constructs. DAPI staining was used as a cell viability marker. *Right:* Quantification of the normalized surface/intracellular fluorescence ratio in HEK293T cells (*n* ≥ 100 cells per group, four independent experiments). (B) *Left:* Representative images of surface and intracellular expression of GluN2B-GFP or GluN2B^Δ^-GFP in HEK293T cells also co-transfected with GluN1 and GluN2A (2A). *Right:* Quantification of the normalized surface/intracellular fluorescence ratio (*n* ≥ 180 cells per group, four independent experiments). (C) *Left:* Representative fluorescence images showing surface and intracellular localization of GluN2A-GFP (2A-GFP) in HEK293T cells also co-transfected with GluN1 and either GluN2B (2B) or GluN2B^Δ^ (2B^Δ^). *Right:* Quantification of the normalized surface/intracellular fluorescence ratio for GluN2A-GFP (*n* ≥ 100 cells per group, four independent experiments). (D) *Left:* Representative images of surface and intracellular immunostaining of GluN2B-GFP and GluN2B^Δ^-GFP in hippocampal neurons from *Grin2b*^+/+^ mice. Yellow outlines indicate neuronal somata used for quantification. *Right:* Normalized surface/intracellular fluorescence ratios for GluN2B-GFP and GluN2B^Δ^-GFP in the somata of hippocampal neurons (*n* ≥ 20 cells per group, four independent experiments). Data are shown as mean ± SEM. Asterisks (*) indicate statistically significant differences (*p* ≤ 0.05, Student’s t-test).

Given that NMDARs can form triheteromeric complexes (Al-Hallaq et al., 2007) (Luo et al., 1997) (Tovar et al., 2013) (Rauner and Kohr, 2011), we next assessed whether co-expression of GluN2A could enhance the surface localization of GluN2B^Δ^. Co-transfection of HEK293T cells with GluN1, GluN2B^Δ^-GFP, and GluN2A resulted in a twofold increase in GluN2B^Δ^-GFP surface expression compared to cells expressing GluN1 and GluN2B^Δ^-GFP alone (*p* < 0.001, Student’s t-test) (Fig. 5A, B). However, co-expression of GluN2B^Δ^ negatively affected the surface expression of GluN2A-GFP (Fig. 5C), suggesting impaired assembly and trafficking of NMDARs containing the GluN2B^Δ^ compared to the wild-type GluN2B subunit. Since protein overexpression in HEK293T cells can engage trafficking and quality-control pathways that differ from those operating in neurons, we also examined cell surface expression of the GluN2B^Δ^-GFP subunit introduced by lentiviral vector transduction in wild-type primary hippocampal neurons. In this system the GluN2B^Δ^-GFP subunit also showed a markedly lower surface expression on the soma compared to the wild-type GluN2B-GFP (86% reduction, *p* < 0.001, Student’s t-test) (Fig. 5D). Despite the fact that our cell surface expression analyses in heterologous cells and wild-type neurons involve high levels of exogenous GluN2B-GFP expression driven by the CMV promoter, our results demonstrate a substantial defect of the GluN2B^Δ^ subunit cell surface expression, which may lead to GluN2B^Δ^ protein degradation in *Grin2b*^+/Δ^ mice.

### Diminished NMDA-induced whole-cell responses in *Grin2b*^+/Δ^ hippocampal neurons

To evaluate the functional impact of the GluN2B^Δ^ subunit in its native context, primary hippocampal microisland cultures were prepared from newborn *Grin2b*^+/+^ and *Grin2b*^+/Δ^ mice and maintained for 7–21 DIV. It is widely accepted that cultured neurons undergo developmental processes similar to those observed *in vivo*, encompassing both morphological and functional aspects (Downes et al., 2012). Developmental increase in dendritic arborization is evident in hippocampal neurons immunostained for β3-tubulin at 7 and 21 DIV (Fig. 6A). Using whole-cell voltage-clamp recordings, we examined the amplitude of responses induced by 100 μM kainic acid or 100 μM NMDA (with 10 μM glycine) at a holding potential of -70 mV. Recordings were restricted to excitatory neurons, identified by the occurrence of autaptically evoked AMPAR-eEPSCs. To account for variations in neuronal size, agonist-induced peak currents—reflecting synaptic and extrasynaptic receptor activation—were normalized to cell capacitance. The AMPA/kainate receptor current density was not consistently affected by genotype; in different DIV age groups, mean current density in *Grin2b*^+/Δ^ neurons ranged from -44% to +12% compared to *Grin2b*^+/+^ neurons (ANOVA: *p* = 0.011 for genotype, *p* = 0.015 for age, *p* = 0.02 for genotype × age interaction; *n* = 57 and *n* = 40 for *Grin2b*^+/Δ^ and *Grin2b*^+/+^, respectively) (Fig. 6B). In contrast, the analysis of NMDAR current densities showed a consistent significant reduction in *Grin2b*^+/Δ^ neurons, ranging from -31% to -59% compared to *Grin2b*^+/+^ neurons in individual age groups. The ANOVA revealed significant effects of genotype (*p* < 0.001) and age (*p* = 0.002), while the genotype × age interaction was not significant (*p* = 0.253); *n* = 59 for *Grin2b*^+/Δ^ and *n* = 54 for *Grin2b*^+/+^ neurons (Fig. 6C).

**Figure 6.**
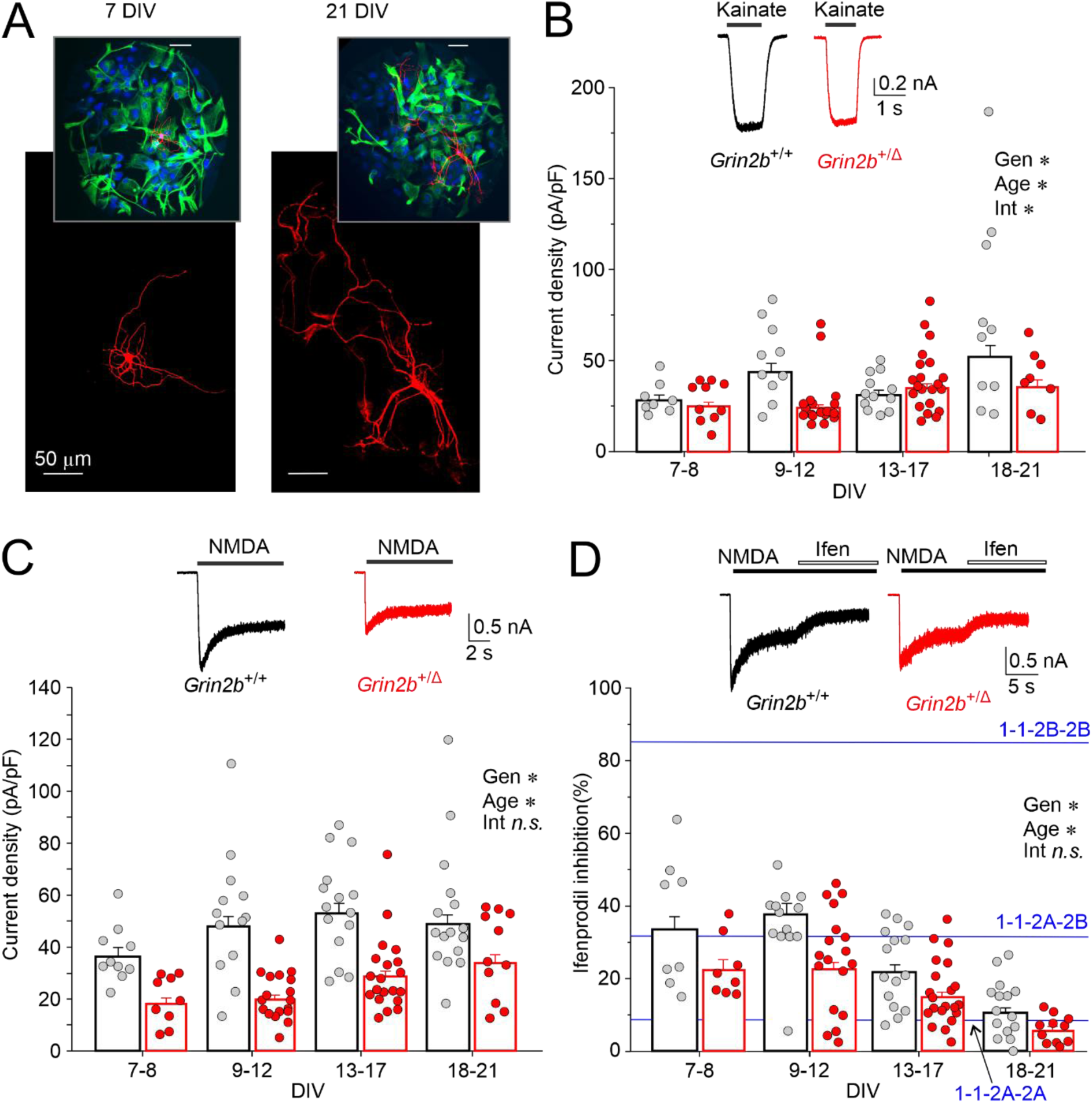
NMDAR whole-cell currents are reduced in *Grin2b*^+/Δ^ neurons. (A) Primary hippocampal microisland cultures stained with β3-tubulin. Representative images show neuronal morphology at 7 DIV (left) and 21 DIV (right). Insets display the same microislands at a lower magnification, with neurons labeled by β3-tubulin (red), astrocytes by glial fibrillary acidic protein (GFAP) (green), and nuclei by DAPI (blue). Scatter plots show the current density distribution in individual hippocampal neurons by DIV. Currents were induced by 100 µM kainate (B) or 100 µM NMDA in the presence of 10 µM glycine (C). Insets show representative whole-cell responses evoked by kainate or NMDA in neurons prepared from *Grin2b*^+/+^ (black) or *Grin2b*^+/Δ^ (red) animals and cultured for 21 DIV. (D) The scatter plot shows the distribution of ifenprodil (3 µM) inhibition of NMDA-evoked currents recorded in individual neurons by DIV. Inset shows the effect of 3 µM ifenprodil on responses to 100 μM NMDA in neurons prepared from *Grin2b*^+/+^ (black) or *Grin2b*^+/Δ^ (red) animals cultured for 21 DIV. Blue lines indicate the expected sum of voltage-dependent and voltage-independent inhibition of diheteromeric GluN1/GluN2B (1-1-2B-2B; 85.3%), and GluN1/GluN2A (1-1-2A-2A; 7.9%), and triheteromeric GluN1/GluN2A/GluN2B (1-1-2A-2B; 31.7%) receptors by 3 µM ifenprodil at -60 mV (with IC_50_ values and Hill slopes from (Hansen et al., 2014)) Asterisks (*) Indicate significant differences with respect to: Gen, genotype; Age, age (DIV); Int, genotype × age interaction. Data were power transformed and tested using ANOVA, followed by pairwise comparisons (LSD method).

Subsequently, we used ifenprodil, a potent inhibitor of NMDARs containing the GluN2B subunit (Williams, 1993), to investigate the source of the reduced NMDAR current densities observed in *Grin2b*⁺^/Δ^ neurons (Fig. 6D). In *Grin2b*⁺^/Δ^ neurons, mean NMDAR current responses were inhibited by ifenprodil (3 μM) by an average of -31% to -47% compared to *Grin2b*^+/+^ neurons across different age groups (ANOVA: *p* < 0.001 for genotype, *p* < 0.001 for age, *p* = 0.792 for genotype × age interaction; *n* = 58 and *n* = 50 cells for *Grin2b*⁺^/Δ^ and *Grin2b*^+/+^, respectively) (Fig. 6D). These findings indicate that *Grin2b*⁺^/Δ^ hippocampal neurons exhibit smaller NMDA-induced current densities and a reduced contribution of GluN2B-containing NMDARs. Additionally, the degree of ifenprodil block of NMDA-induced currents declines with age in both genotypes, consistent with the developmental switch from GluN2B to GluN2A subunits (McKay et al., 2018). We also analyzed sex differences in AMPA/kainate and NMDAR current densities, as well as the degree of NMDAR current inhibition by ifenprodil. No significant differences were observed between female and male neurons (data are summarized in Table S4).

### Accelerated deactivation of NMDAR eEPSC in *Grin2b*^+/Δ^ mice

To evaluate the impact of the *Grin2b*^Δ^ variant on excitatory synaptic transmission, we recorded AMPAR- and NMDAR-mediated components of eEPSCs, pharmacologically isolated using AP-5 or NBQX (see Methods), in hippocampal microisland cultures. We first reliably detected eEPSCs at 6–7 DIV. AMPAR-eEPSC peak current densities increased with age in both *Grin2b*⁺^/Δ^ and *Grin2b*^+/+^ neurons, with no significant genotype effect (ANOVA: genotype *p* = 0.774, age *p* < 0.001, genotype × age interaction *p* = 0.589; *n* = 54 and *n* = 58 for *Grin2b*⁺^/Δ^ and *Grin2b*^+/+^, respectively; Fig. 7A). NMDAR-eEPSC peak amplitudes also increased with age in both genotypes, with no statistically significant overall difference between genotypes (ANOVA: genotype *p* = 0.051, age *p* < 0.001, genotype × age interaction *p* = 0.062; *n* = 50 and *n* = 55 for *Grin2b*⁺^/Δ^ and *Grin2b*^+/+^, respectively; Fig. 7B). Notably, the mean deactivation time course of NMDAR-eEPSCs was significantly faster in *Grin2b*⁺^/Δ^ neurons—by 22% to 41%— compared to *Grin2b*^+/+^ neurons across age groups, consistent with a reduced contribution of GluN2B-containing NMDARs. Deactivation kinetics accelerated with age in both genotypes, as expected during synaptic maturation (ANOVA: genotype *p* < 0.001, age *p* = 0.015, genotype × age interaction *p* = 0.438; *n* = 50 and *n* = 57 for *Grin2b*⁺^/Δ^ and *Grin2b*^+/+^, respectively; Fig. 7C). Results summarized in Table S4 revealed no significant differences between sexes.

**Figure 7.**
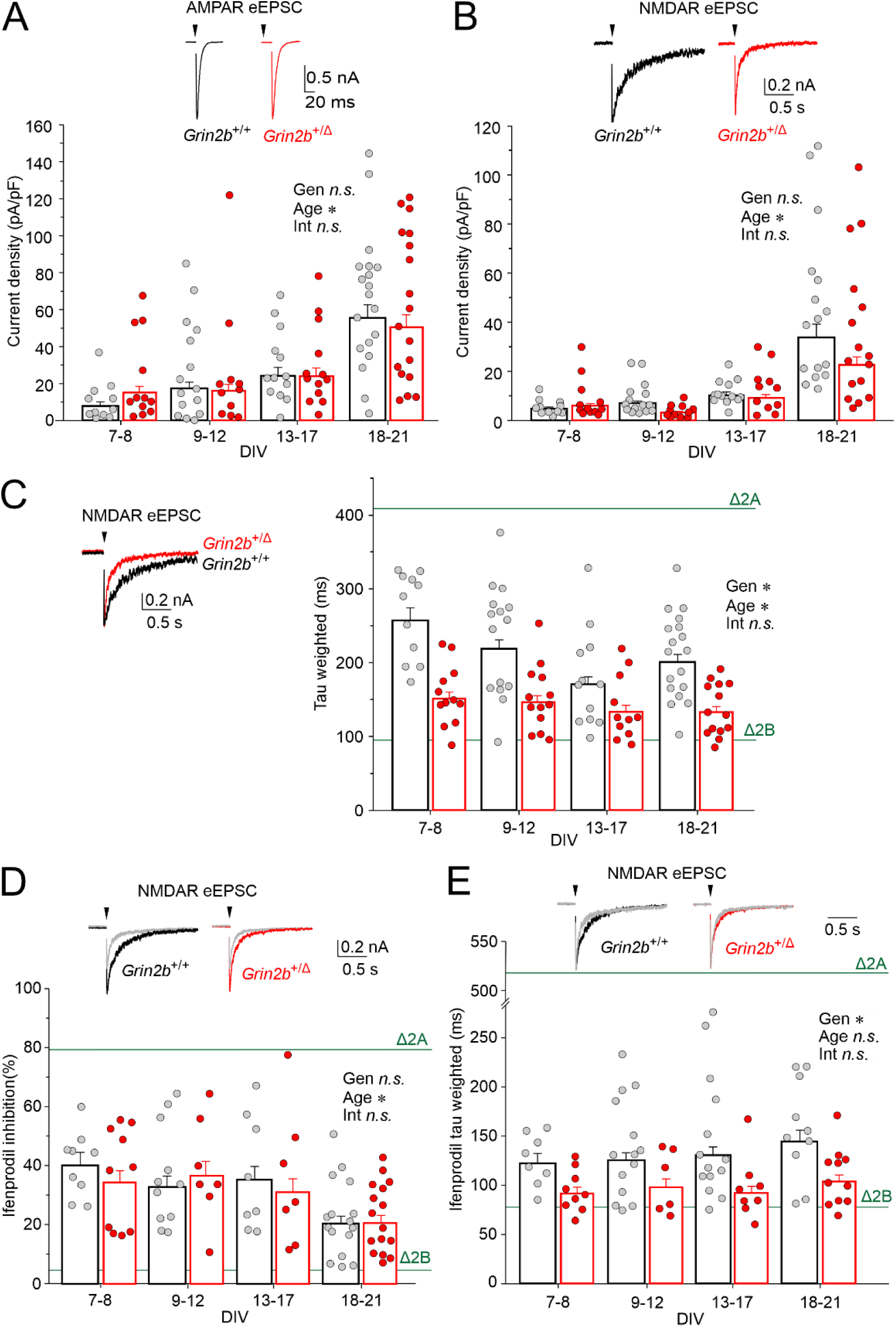
NMDAR-eEPSCs in *Grin2b*^+/Δ^ neurons have faster deactivation. (A–B) Scatter plots show the distribution of peak current densities for AMPAR-eEPSCs (A) and NMDAR-eEPSCs (B) recorded from individual *Grin2b*^+/+^ (gray) and *Grin2b*⁺^/Δ^ (red) neurons, plotted as a function of DIV. Insets: representative traces from *Grin2b*^+/+^ (black) and *Grin2b*⁺^/Δ^ (red) neurons at 8 DIV. (C) Weighted deactivation time constants of NMDAR-eEPSCs plotted by DIV. Inset: scaled NMDAR-eEPSCs from *Grin2b*^+/+^ (black) and *Grin2b*⁺^/Δ^ (red) neurons at 8 DIV. (D) Ifenprodil (3 µM) inhibition of NMDAR-eEPSC peak amplitudes plotted by DIV. (C) Weighted deactivation time constants of NMDAR-eEPSCs recorded in the presence of ifenprodil (3 µM) and plotted by DIV. Inset: example traces from *Grin2b*^+/+^ (black) and *Grin2b*⁺^/Δ^ (red) neurons recorded at 15 DIV, showing the effect of ifenprodil (gray). The green lines in C, E, and D indicate mean weighted deactivation time constants of NMDAR-eEPSCs recorded from mouse hippocampal pyramidal neurons at -60 mV (C), mean ifenprodil (3 µM) inhibition of NMDAR-eEPSCs at +40 mV(D), and mean weighted deactivation time constants of NMDAR-eEPSCs recorded in the presence of ifenprodil (3 µM) after conditional knockout of GluN2A (Δ2A) or GluN2B (Δ2B) subunits (Gray et al., 2011). Asterisks (*) indicate significant differences for: Gen, genotype; Age, age (DIV); Int, genotype × age interaction. Data were power transformed and analyzed by ANOVA followed by pairwise comparisons (LSD method).

To further probe the contribution of GluN2B-containing receptors, we applied ifenprodil (3 µM), a selective GluN2B antagonist. Ifenprodil inhibited NMDAR-eEPSC peak amplitudes to a similar degree in both genotypes, with inhibition decreasing with age (ANOVA: genotype *p* = 0.755, age *p* = 0.001, genotype × age interaction *p* = 0.843; *n* = 40 and *n* = 44 for *Grin2b*⁺^/Δ^ and *Grin2b*^+/+^, respectively; Fig. 7D). These findings are consistent with previous reports showing developmental decrease in ifenprodil inhibition of EPSCs likely reflecting increasing synaptic incorporation of GluN2A subunits and a corresponding reduction in GluN2B subunits (Hestrin, 1992) (Flint et al., 1997) (Gray et al., 2011). The decay kinetics of NMDAR-eEPSC recorded in the presence of ifenprodil were on average accelerated by 30% in *Grin2b*⁺^/Δ^ neurons and by 33% in *Grin2b*^+/+^ neurons when compared to corresponding controls recorded without ifenprodil. The mean deactivation time course of NMDAR-eEPSCs recorded in the presence of ifenprodil was significantly faster in *Grin2b*⁺^/Δ^ neurons than in *Grin2b*^+/+^ neurons across all age groups (ANOVA: genotype *p* < 0.001, age *p* = 0.459, genotype × age interaction *p* = 0.965; *n* = 33 and *n* = 46 for *Grin2b*⁺^/Δ^ and *Grin2b*^+/+^, respectively; Fig. 7E). Our eEPSC data suggest that the overall synaptic NMDAR content may be largely preserved in *Grin2b*⁺^/Δ^ neurons (Fig. 7B), despite the observed reduction in whole-cell NMDAR current density (Fig. 6C). However, NMDAR subunit composition is likely altered in both the synaptic and the extrasynaptic compartment, with a decrease in GluN2B contribution to NMDAR signaling in *Grin2b*⁺^/Δ^ compared to *Grin2b*⁺^/^⁺ neurons.

### Increased amplitude of AMPAR-mEPSC in *Grin2b*^+/Δ^ mice

NMDARs regulate glutamatergic transmission by limiting AMPAR incorporation at developing synapses (Alvarez et al., 2007). *Grin2b* deletion increases synaptic AMPAR localization and AMPAR-mEPSC amplitudes (Hall et al., 2007) (Kelsch et al., 2014) or increases the number of functional synapses as suggested by increased AMPAR-mEPSC frequency (Gray et al., 2011).

To investigate possible alterations of AMPAR-mediated synaptic transmission in a disease-relevant model, we recorded action potential-independent AMPAR-mEPSCs from *Grin2b*⁺^/Δ^ and *Grin2b*^+/+^ single-neuron hippocampal microisland cultures at different DIV (Candelas Serra et al., 2024). The amplitude of AMPAR-mEPSCs in *Grin2b*⁺^/Δ^ neurons was 32% higher than in age-matched *Grin2b*^+/+^ neurons from littermates (ANOVA: *p* < 0.001 for genotype, *p* = 0.841 for age, *p* = 0.243 for genotype × age interaction; *n* = 52 and *n* = 43 for *Grin2b*⁺^/Δ^ and *Grin2b*^+/+^, respectively) (Fig. 8A, B). The frequency of AMPAR-mEPSCs in *Grin2b*⁺^/Δ^ neurons was not consistently different from *Grin2b*^+/+^ neurons (ANOVA: *p* = 0.161 for genotype, *p* < 0.001 for age, *p* = 0.024 for genotype × age interaction; *n* = 52 and *n* = 43 for *Grin2b*⁺^/Δ^ and *Grin2b*^+/+^, respectively; see Fig. 8C).

**Figure 8.**
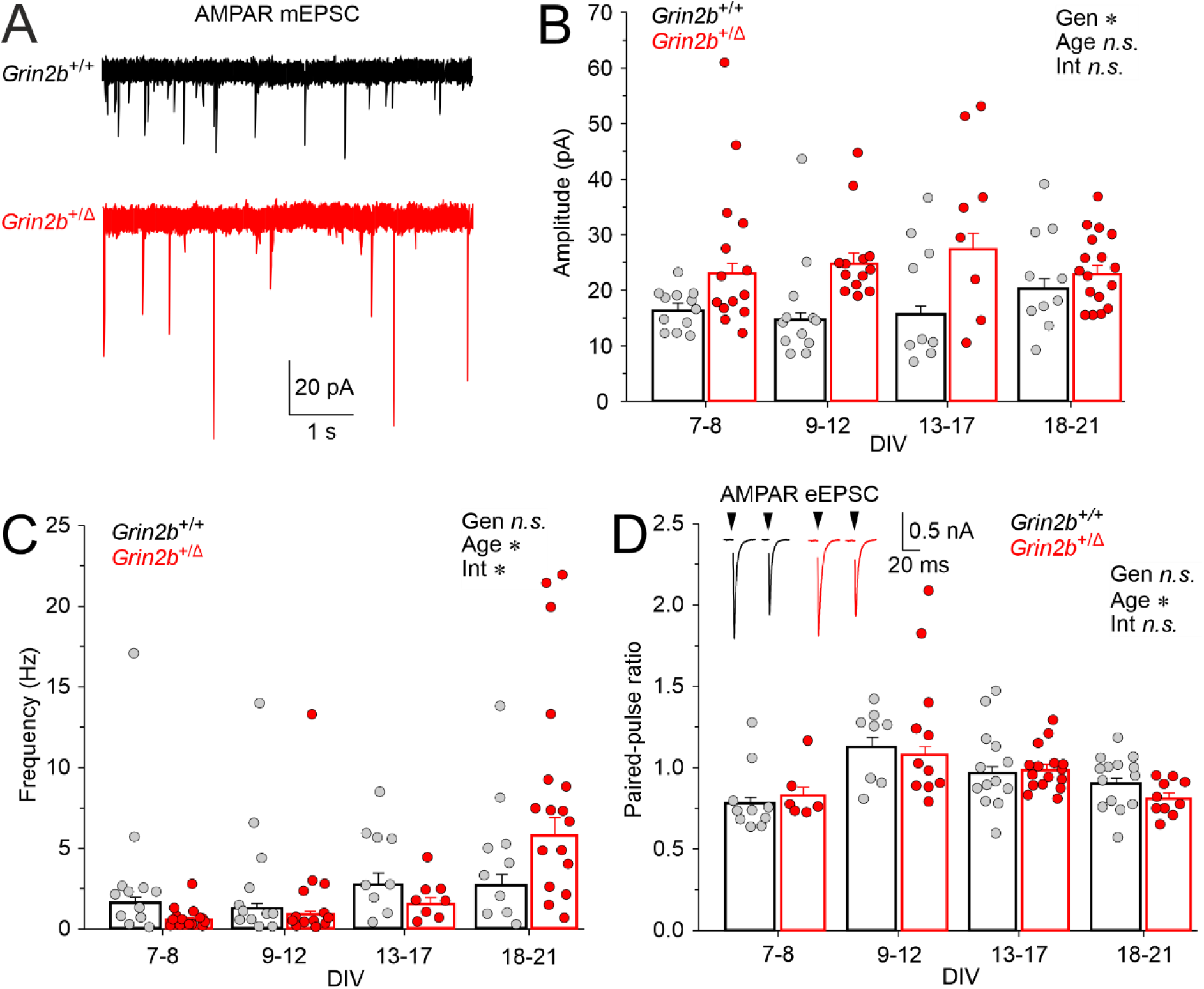
Properties of AMPAR-mEPSCs and glutamate release in *Grin2b*^+/Δ^ and *Grin2b*^+/+^ neurons. (A) Representative recordings of AMPAR-mEPSCs in neurons from *Grin2b*^+/+^ and *Grin2b*⁺^/Δ^ animals (DIV = 8). (B) Scatter plot showing the distribution of mean AMPAR-mEPSC amplitudes recorded from individual *Grin2b*^+/+^ and *Grin2b*⁺^/Δ^ neurons (each point represents the mean AMPAR-mEPSC amplitude recorded over a 2-min interval). (C) Scatter plot showing the distribution of AMPAR-mEPSC frequencies recorded from individual *Grin2b*^+/+^ and *Grin2b*⁺^/Δ^ neurons. (D) Scatter plot showing the distribution of the AMPAR PPR in individual *Grin2b*^+/+^ and *Grin2b*⁺^/Δ^ neurons. Asterisks (*) indicate significant differences for: Gen, genotype; Age, age (DIV); Int, genotype × age interaction. Data were power transformed and analyzed using ANOVA followed by pairwise comparisons (LSD method).

Furthermore, the PPR of AMPAR-eEPSCs was similar between *Grin2b*⁺^/Δ^ and *Grin2b*^+/+^ neurons (ANOVA: *p* = 0.679 for genotype, *p* = 0.016 for age, *p* = 0.560 for genotype × age interaction) (Fig. 8D) (Velasco and Llobet, 2020). Together these findings indicate that that a monoallelic PTV in *Grin2b*, and the consequent reduction in GluN2B-mediated NMDAR signaling, promotes an increase in the density or function of synaptic AMPARs without affecting the number of functional AMPAR-containing glutamatergic synapses in excitatory hippocampal neurons or the properties of presynaptic glutamate release.

### Behavioral impact of GluN2B^Δ^ in *Grin2b*^+/Δ^ mice

Reduced NMDAR function in mice is associated with a range of behavioral abnormalities, including motor symptoms, altered anxiety levels, impaired sensorimotor gating, changes in social behavior, and cognitive deficits (Lee and Zhou, 2019). Mice heterozygous for the conventional *Grin2b* deletion (Kutsuwada et al., 1996) show a relatively mild behavioral phenotype characterized by impaired prepulse inhibition of the acoustic startle (Takeuchi et al., 2001), hypoactivity in the open field, and reduced anxiety (Shin et al., 2020). To assess the behavioral effects of a monoallelic *Grin2b* PTV, we conducted a series of behavioral tests comparing adult *Grin2b*⁺^/Δ^ mice with their *Grin2b*^+/+^ littermates.

### Open field and PPI tests in *Grin2b*⁺^/Δ^ mice

In the open field test, which evaluates general activity and anxiety in a novel environment, adult male *Grin2b*⁺^/Δ^ mice exhibited reduced activity, covering 22% less total distance and spending significantly more time resting compared to *Grin2b*^+/+^ males. In contrast, no activity differences were observed between female *Grin2b*⁺^/Δ^ and *Grin2b*^+/+^ mice, whose activity levels were similar to *Grin2b*^+/+^ males (Fig. 9A, Fig. S3, Table S5). The analysis of thigmotaxis also revealed a sex-dependent behavioral phenotype. Male *Grin2b*⁺^/Δ^ mice exhibited 27% fewer entries into the center zone and covered 36% less distance in the center zone compared to *Grin2b*^+/+^ males, indicating increased anxiety. No significant differences were observed between female *Grin2b*⁺^/Δ^ and *Grin2b*^+/+^ mice (Fig. 9A, Fig. S3, Table S5). In contrast, no genotype-dependent differences in anxiety were observed in the elevated plus maze test, as both male and female *Grin2b*⁺^/Δ^ mice spent similar time in the closed and open arms as their *Grin2b*^+/+^ littermates (Fig. S4, Table S5).

**Figure 9.**
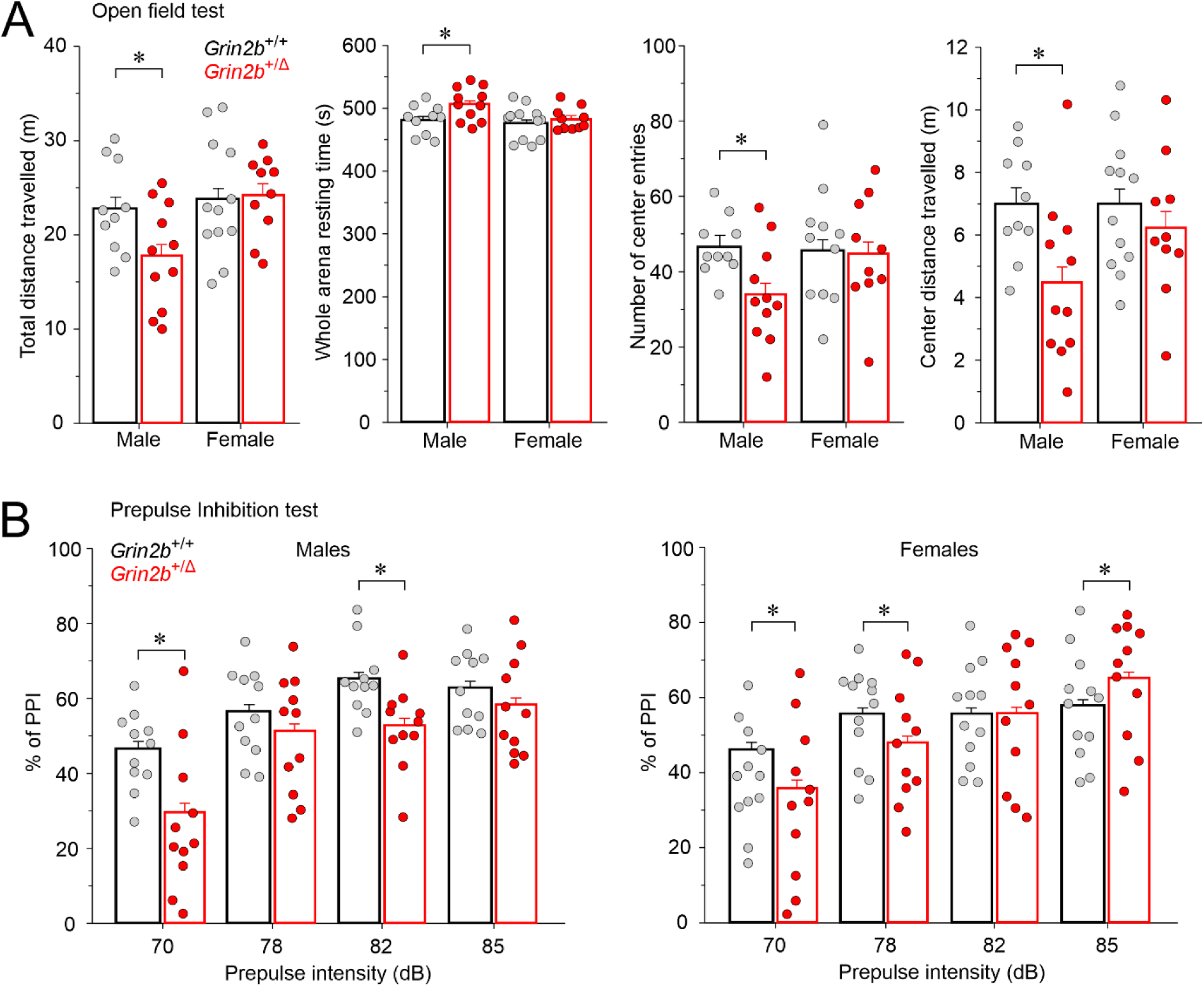
Adult male *Grin2b*⁺^/Δ^ mice exhibit reduced activity and increased anxiety, while both sexes show impaired sensorimotor gating. (A) Quantification of total distance traveled, whole-arena resting time, number of center entries, and center distance traveled in *Grin2b*⁺^/Δ^ and *Grin2b*^+/+^ mice (9–15 weeks old) assessed in the open field test. Data are presented as mean ± SEM. Sample sizes (*Grin2b*^+/+^/*Grin2b*⁺^/Δ^): open field test, *n* = 10/11 males, *n* = 12/10 females. Asterisks (*) indicate significant genotype differences, as analyzed by Student’s t-test. (B) Quantification of the percentage of the inhibition of the startle response as a function of prepulse intensity in the PPI test. Sample sizes (*Grin2b*^+/+^/*Grin2b*⁺^/Δ^) PPI, *n* = 11/11 males, *n* = 12/11 females. Asterisks (*) indicate significant genotype differences, as analyzed by ANOVA followed by post-hoc tests using the LSD method. Behavioral assays were performed using age-matched animals. The numerical data underlying this figure can be found in Table S5.

The PPI test was used to assess sensorimotor gating, a process often impaired in connection with NMDA hypofunction in animal models of schizophrenia and ASD (Perry et al., 2007), obsessive-compulsive disorder (Swerdlow et al., 1993) and other neuropsychiatric conditions (Lee and Zhou, 2019). PPI measures the suppression of the startle reflex when a weak, non-startling prepulse precedes a strong startle stimulus. The PPI ratio was lower in *Grin2b*⁺^/Δ^ males compared to *Grin2b*^+/+^ males across all prepulse intensities. However, significant reductions were observed only at 70 and 82 dB. Similarly, *Grin2b*⁺^/Δ^ females showed a significantly lower PPI ratio at 70 and 78 dB but an increase at 85 dB (Fig. 9B, Table S5).

### Cognitive performance in *Grin2b*⁺^/Δ^ mice

Homozygous mice with a conditional postnatal deletion of *Grin2b* in the forebrain exhibit deficits in learning and memory (von Engelhardt et al., 2008). To determine whether heterozygous *Grin2b*⁺^/Δ^ mice show an impairment of cognitive function, we evaluated their performance in associative memory and recognition tasks. In the contextual and cued fear conditioning test, *Grin2b*⁺^/Δ^ mice exhibited normal activity levels during the baseline period and showed equivalent foot shock reactivity compared to *Grin2b*^+/+^ controls. During the post-training memory test, *Grin2b*⁺^/Δ^ mice displayed context freezing, with no significant difference from *Grin2b*^+/+^ mice (Fig. 10A, Table S5). Cognitive performance was further assessed using the novel object recognition test. The preference for the novel object did not differ significantly between *Grin2b*⁺^/Δ^ males and *Grin2b*^+/+^ controls; however, it was reduced in *Grin2b*⁺^/Δ^ compared to *Grin2b*^+/+^ females (Fig. 10B, Table S5).

**Figure 10.**
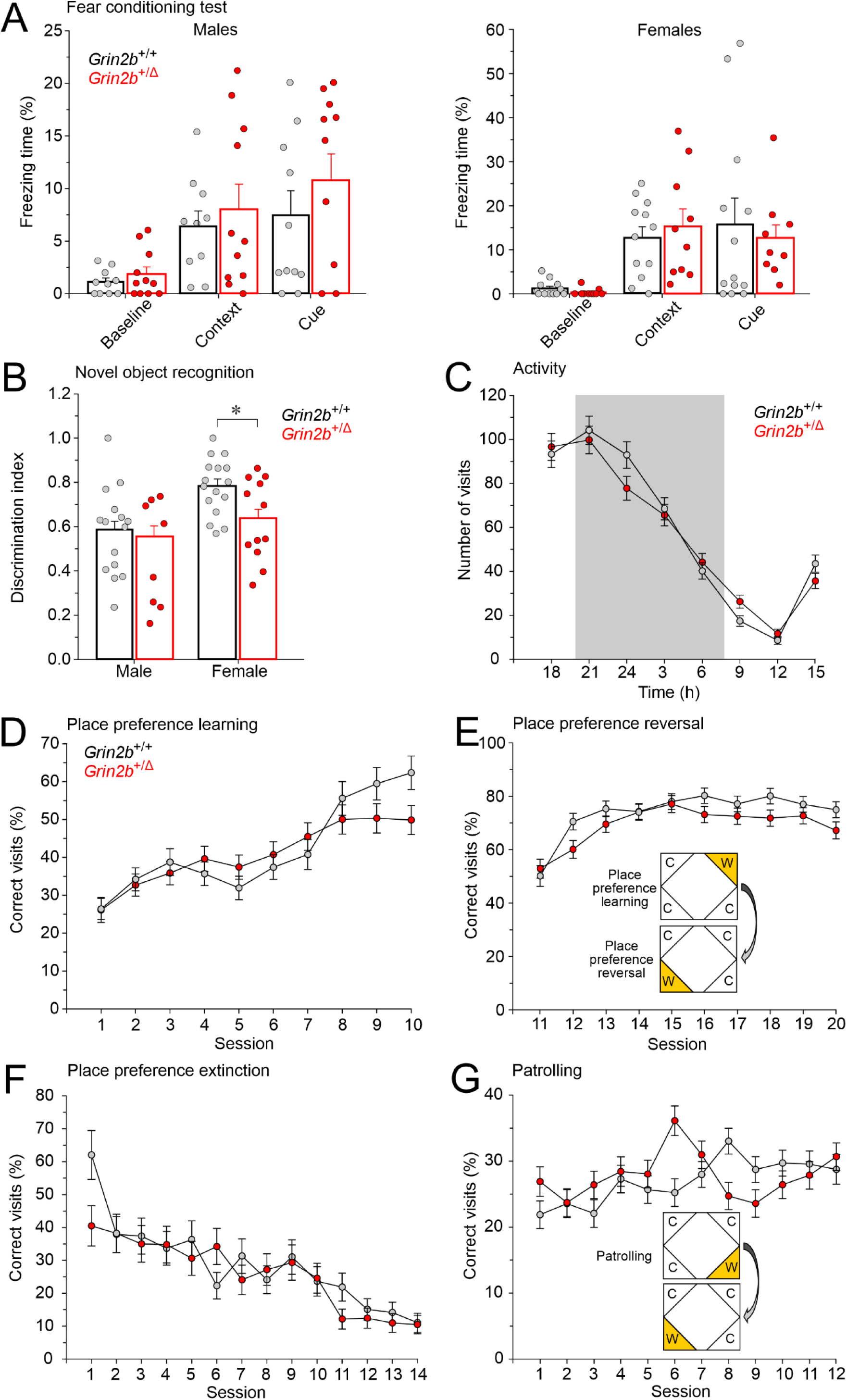
Normal learning and memory in *Grin2b*^+/Δ^ mice. (A) Fear conditioning test showing normal baseline, context, and cue freezing responses in male and female *Grin2b*^+/Δ^ mice. Group sizes *Grin2b*^+/+^/*Grin2b*^+/Δ^: *n* = 10/11 males; *n* = 12/10 females. (B) Novel object recognition performance was reduced in *Grin2b*^+/Δ^ females. Group sizes *Grin2b*^+/+^/*Grin2b*^+/Δ^: *n* = 15/8 males; *n* = 16/12 females. (C–G) IntelliCage-based behavioral testing of female *Grin2b*^+/Δ^ and *Grin2b*^+/+^ mice. (C) Number of visits during the 4-day free adaptation phase (all corners open, unrestricted water access). (D–G) Session scores (% visits to the correct corner) for place preference (D), place preference reversal (E), place preference extinction (F), and patrolling behavior (G). Trial design: C = unconditionally closed; W = water available at designated corners during drinking sessions (9–11 PM and 3–5 AM). Data are shown as mean ± SEM and were analyzed by Repeated Measures ANOVA. Group sizes *Grin2b*^+/+^/*Grin2b*^+/Δ^: *n* = 9/8 females. No significant genotype differences were observed in any test. See Table S5 for numerical data.

To further assess cognitive performance, we tested *Grin2b*⁺^/Δ^ mice in IntelliCages, an automated system that evaluates behavior in group-housed animals without human intervention (Candelas Serra et al., 2024). The IntelliCages use door-guarded water reservoirs, with individual drinking responses tracked via subcutaneous transponders. With the start of restricted access to water, male *Grin2b*⁺^/Δ^ mice developed aggressive behavior, necessitating the termination of their participation for animal welfare reasons. Therefore, only females were tested in IntelliCages. Circadian rhythmicity assessed during the free adaptation phase remained intact in *Grin2b*⁺^/Δ^ females, as their visits to the drinking corners followed the normal day/night cycle, with no genotype differences (*p* = 0.927, Repeated Measures ANOVA) (Fig. 10C, Table S5). *Grin2b*⁺^/Δ^ females performed similarly to *Grin2b*^+/+^ females in place preference learning (*p* = 0.443, Repeated Measures ANOVA) and place preference reversal learning, which measures cognitive flexibility (*p* = 0.257, Repeated Measures ANOVA) (Fig. 10D, E). Place preference extinction and patrolling behavior, an indicator of working memory, also showed no significant genotype differences (*p* = 0.113 and *p* = 0.314, Repeated Measures ANOVA, respectively) (Fig. 10F, G, Table S5).

### Social behavior in *Grin2b*⁺^/Δ^ mice

To assess social and repetitive behavior in *Grin2b*⁺^/Δ^ mice, we conducted the social interaction test and the nestlet shredding test (Silverman et al., 2010; Sonzogni et al., 2020). *Grin2b*⁺^/Δ^ males exhibited normal social behavior, with no significant differences in time spent sniffing the juvenile target compared to *Grin2b*^+/+^ controls. In contrast, *Grin2b*⁺^/Δ^ females spent relatively more time exploring the juvenile compared to the object-containing chamber (Fig. S5, Table S5). Nestlet shredding performance did not differ significantly between *Grin2b*⁺^/Δ^ and *Grin2b*^+/+^ mice (Fig. S6, Table S5).

While homozygous *Grin2b* deletion is lethal (Kutsuwada et al., 1996), heterozygous conventional *Grin2b* deletion is associated with relatively moderate behavioral phenotype characterized by hypoactivity and impaired sensorimotor gating (Takeuchi et al., 2001) (Shin et al., 2020). Similarly, *Grin2b*⁺^/Δ^ mice exhibit only moderate behavioral changes. These include hypoactivity and increased anxiety in males, and impaired sensorimotor gating in both sexes. Learning and memory, and social behavior remain largely unaffected.

## Discussion

In this study, we used *in vitro* and *in vivo* methods to analyze mice heterozygous for a frameshift *Grin2b* mutation, with the goal to uncover molecular, cellular, and behavioral aspects relevant to human GluN2B PTVs. The GluN2B^Δ^ subunit truncated from the M4 transmembrane helix shows severely impaired surface expression in heterologous expression systems and may be largely degraded in *Grin2b*⁺^/Δ^ brain tissue. Proteomic analysis revealed reduced GluN2B levels predicting a shift toward an increased proportion of diheteromeric GluN1/GluN2A receptors. Hippocampal neurons from *Grin2b*⁺^/Δ^ mice showed reduced NMDA-induced currents less sensitive to ifenprodil, faster NMDAR-eEPSC deactivation, and increased AMPAR-mEPSC amplitudes. Behaviorally, *Grin2b*⁺^/Δ^ males exhibited reduced locomotion in the novel environment and increased anxiety-like behavior, while both sexes showed impaired sensorimotor gating. Learning, memory, and social behavior was not affected. Together our results suggest that a monoallelic *Grin2b* PTV alters NMDAR subunit composition and leads to relatively moderate behavioral effects.

### Aberrant *Grin2b* mRNA is not selectively downregulated in *Grin2b*⁺^/Δ^ mice

The RT-qPCR analysis of hippocampal tissue from *Grin2b*⁺^/Δ^ mice revealed only a modest reduction in *Grin2b* mRNA levels (Fig. 2A, B). In many prokaryotic and eukaryotic systems, the presence of a premature termination codon typically engages the NMD pathway to degrade aberrant transcripts and prevent the synthesis of truncated proteins (Kurosaki et al., 2019) (Lee et al., 2021). However, in *Grin2b*⁺^/Δ^ mice, the relative proportions of wild-type and the premature stop codon-containing *Grin2b* mRNAs were equal (Fig. 2C, D), challenging the involvement of canonical NMD mechanisms (Howe and Patani, 2023). Generally, NMD does not target transcripts with a premature stop codon located within the penultimate exons if they occur less than 50–55 nucleotides upstream of the final exon-exon junction, or with a premature stop codon located anywhere within the last exon, unless an unusually long 3’ UTR is present (Kurosaki et al., 2019) (Lee et al., 2021). The stop codon in the *Grin2b*^Δ^ allele lies approximately 85 nucleotides upstream of the final exon-exon junction, thus just out of range for the transcript to escape NMD. This suggests that the aberrant *Grin2b*^Δ^ transcripts may instead be spared due to suppressed NMD activity in neurons. Neurons exhibit reduced NMD efficiency compared to other cell types, potentially increasing their susceptibility to truncated protein expression from premature stop codon-bearing transcripts (Lee et al., 2021), in addition to a broader tolerance for some frameshift variants (Wang et al., 2022). The parallel modest downregulation of *Grin1* and *Grin2b* (wild-type and PTV; Fig. 2C, D) mRNA (Fig. 2A, B) suggests a shared regulatory mechanism affecting mRNA processing.

### Consequences of subunit availability for NMDAR subunit composition in *Grin2b*⁺^/Δ^ mice

Despite only a modest reduction in *Grin2b* mRNA levels in hippocampal tissue, proteomic analysis revealed that the wild-type GluN2B subunit levels were substantially reduced and the GluN2B^Δ^ subunit was undetectable by mass spectrometry in the hippocampal proteome of male *Grin2b*⁺^/Δ^ mice. This suggests that GluN2B^Δ^ is efficiently eliminated by protein quality control mechanisms such as the unfolded protein response, ER-associated degradation, or ER-phagy (Benske et al., 2022). Interestingly, GluN2A protein levels were slightly elevated in the *Grin2b*⁺^/Δ^ hippocampus despite unchanged *Grin2a* mRNA levels, potentially reflecting reduced internalization or degradation of GluN2A-containing NMDARs.

Changes of relative copy numbers of NMDAR subunits in the presence of disease-associated variants may provide insights into disease pathophysiology. Proteomic analysis of *Grin2b*^+/+^ hippocampal tissue revealed that the relative CN values of GluN1, GluN2A, and GluN2B subunits are approximately equivalent. Given that NMDARs are tetramers composed of two GluN1 and two GluN2 subunits (Laube et al., 1998), this suggests that the availability of GluN1 may be the limiting factor in determining total NMDAR numbers in hippocampal neurons. In the *Grin2b*⁺^/Δ^ hippocampus, the relative CN values of GluN1, GluN2A, and GluN2B were altered by –9%, +16%, and –38%, respectively, compared to *Grin2b*^+/+^ controls. If GluN1 availability limits the total number of NMDARs, the 9% reduction in GluN1 would lead to a small decrease in overall receptor abundance. However, the total surplus of GluN2 subunits above GluN1 would buffer the loss of GluN2B, maintaining high receptor numbers but shifting receptor subunit composition. Our calculations predict a 2.2-fold reduction in GluN1/GluN2B diheteromers, a 1.7-fold increase in GluN1/GluN2A diheteromers, and a modest increase in triheteromeric GluN1/GluN2A/GluN2B receptors. It is important to note that in the *Grin2b*^+/Δ^ mice, all the assembled NMDARs will retain their normal function. In contrast, the *Grin2b*^+/L825V^ mice showed no change in the relative CN for any of the NMDAR subunits. In the case of the missense variant, total receptor numbers and receptor subunit compositions remain unchanged, but receptors containing the GluN2B(L825V) variant are hypofunctional due to a reduced channel open probability (Candelas Serra et al., 2024).

Some evidence, however, argues against the limiting role of GluN1 availability in NMDAR assembly. Perhaps most intriguingly, *GRIN1* haploinsufficiency in humans does not appear to be pathogenic (Santos-Gomez et al., 2021). In addition, NMDA-induced whole-cell currents in hippocampal and cerebellar neurons from mice heterozygous for a conventional *Grin1* deletion are not reduced compared to wild-type (Forrest et al., 1994), and in cerebellar granule cell cultures, overexpression of GluN2 subunits but not of the GluN1 subunit increases NMDAR current amplitudes (Prybylowski et al., 2002). Further research will be needed to better understand how subunit availability may influence NMDAR assembly.

### Comparing proteomic and functional profiles of NMDARs in *Grin2b*⁺^/Δ^ mice

The interpretation of the observed relative subunit abundance is constrained by the nature of mass spectrometry-based proteomics, which measures both intracellular and surface-expressed NMDAR subunits. In contrast, electrophysiological recordings reflect the functional surface (extrasynaptic and/or synaptic) receptor populations. Previous studies have shown that NMDAR subunit composition regulates ER export and synaptic targeting (Barria and Malinow, 2002) (Groc et al., 2006) (Horak et al., 2008). As receptor trafficking is subunit dependent, the intracellular and surface receptor pools may differ in subunit composition.

Nevertheless, our electrophysiological results generally support the reduced relative GluN2B content suggested by the proteomic analysis. *Grin2b^+^*^/Δ^ neurons exhibited significantly decreased whole-cell NMDAR current densities with reduced sensitivity to ifenprodil, a selective GluN2B antagonist (Williams, 1993). Peak NMDAR-eEPSC amplitudes were preserved but their deactivation kinetics were faster in *Grin2b^+^*^/Δ^ than in *Grin2b^+^*^/*+*^ neurons, suggesting a reduced GluN2B subunit contribution, although NMDAR-eEPSC sensitivity to ifenprodil was not reduced. Different impact of the *Grin2b*^Δ^ variant on whole-cell *vs.* synaptic responses may reflect distinct subunit compositions at synaptic *vs.* extrasynaptic sites, consistent with previous studies (Groc et al., 2006) (Petralia et al., 2010).

For further insight into NMDAR subunit compositions in *Grin2b^+^*^/*+*^ and *Grin2b^+^*^/Δ^ neurons we compared our data with published results from studies where GluN2A and GluN2B content was experimentally manipulated. For whole-cell NMDA-induced currents (Fig.6) we considered the data from (Hansen et al., 2014), where recombinant diheteromeric GluN1/GluN2A or GluN1/GluN2B, or triheteromeric GluN1/GluN2A/GluN2B receptors were studied in a heterologous expression system. The degree of ifenprodil (3 μM) inhibition of NMDA-induced currents in *Grin2b^+^*^/Δ^ neurons across all DIV age groups is lower than the expected inhibition of triheteromeric GluN1/GluN2A/GluN2B receptors, consistent with the presence of a high proportion of diheteromeric GluN1/GluN2A receptors. In the older *Grin2b^+^*^/Δ^ neurons (18–21 DIV) ifenprodil inhibition reaches the low level expected for a pure population of diheteromeric GluN1/GluN2A receptors.

Comparing NMDAR-eEPSCs in *Grin2b^+^*^/*+*^ and *Grin2b^+^*^/Δ^ neurons with NMDAR-eEPSCs in neurons with a conditional deletion of the GluN2A (ΔGluN2A) or the GluN2B (ΔGluN2B) subunit (Gray et al., 2011) (Fig. 7) reveals that the deactivation time course and ifenprodil inhibition of NMDAR-eEPSCs in *Grin2b^+^*^/Δ^ neurons approaches the values observed in neurons completely lacking GluN2B (ΔGluN2B) but does not quite reach them even in the oldest age group (Fig. 7C, D). This indicates that the GluNB content in synaptic NMDARs is at least partially preserved, perhaps more so than in the extrasynaptic receptor populations (Fig. 6). NMDAR-eEPSC deactivation time course is accelerated by ifenprodil in both genotypes (Fig. 7E) and in *Grin2b^+^*^/Δ^ neurons the remaining NMDAR-eEPSCs resemble diheteromeric GluN1/GluN2A eEPSCs recorded in ΔGluN2B neurons (Gray et al., 2011), consistent with fast-deactivating GluN1/GluN2A-mediated responses (Erreger et al., 2007) (Wyllie et al., 2013). Ifenprodil-treated *Grin2b^+^*^/*+*^ neurons showed slower and more variable NMDAR-eEPSC deactivation kinetics, consistent with some remaining contribution of GluN2B-containing NMDARs to the synaptic response.

### Comparison between human *GRIN2B* variant phenotypes and mouse model outcomes

Both mouse strains, *Grin2^+^*^/Δ^ (this study) and *Grin2b*^+/L825V^ (Candelas Serra et al., 2024), enabled system-level assessment of the effects of a PTV and a missense loss-of-function variant, respectively, each affecting the GluN2B subunit at a similar site. The *Grin2b*⁺^/Δ^ mice exhibited a milder phenotype: males showed hypoactivity and increased anxiety, and both sexes had impaired sensorimotor gating, while learning, memory, and social behavior were preserved. In contrast, *Grin2b*^+/L825V^ mice were more severely affected—males displayed hypoactivity, heightened anxiety, and sensorimotor gating deficits, while females exhibited cognitive inflexibility and working memory impairments without anxiety-related changes (Candelas Serra et al., 2024). These results are consistent with the broader observation that PTVs, which account for approximately 23% of disease-associated *GRIN* gene variants, are generally linked to less severe DD/ID phenotypes than missense variants (Platzer et al., 2017) (Strehlow et al., 2019) (Santos-Gomez et al., 2021). The comparison of the two mouse strains also generally aligns with the specific case reports of the severely affected patient carrying the missense GluNB-L825V variant (Awadalla et al., 2010) (Tarabeux et al., 2011) and the relatively more moderate case of the patient carrying the GluN2B-E839* variant (Santos-Gomez et al., 2021), which is similar to the PTV studied here (GluN2B-L825Ffs*15).

The behavioral phenotypes observed in *Grin2b*⁺^/Δ^ and *Grin2b*^+/L825V^ mice, however, were generally milder than the more substantial clinical symptoms associated with human *GRIN2B* variants. Several potential explanations can be considered. First, while the mouse models isolate the *Grin2b* mutations on an unaltered wild-type genetic background, affected patients may carry additional genetic variants that contribute to disease severity (Hassan et al., 2018). Second, if, in the case of GluN2B PTVs, disease pathology involves the altered distribution of functionally intact GluN2A- and/or GluN2B-containing NMDARs, and since GluN2B expression declines postnatally (Bar-Shira et al., 2015), one might expect symptom severity to lessen with age. Further study of the developmental trajectories of GRIN variant-associated phenotypes is necessary. Finally, it may be challenging to capture the complexity of human behavior in mouse models. Another mouse model of an ASD-associated *GRIN2B* loss-of-function variant (GluN2B-C456Y; (Shin et al., 2020) also showed a relatively mild behavioral phenotype including hypoactivity in the open field and age-dependent changes in anxiety-like behaviors, but normal social interactions. Despite some limitations, the existing mouse models of human *GRIN2B* variant-associated neurodevelopmental disorders are consistent with the general observations from clinical data, give insights into possible disease mechanisms, and provide the platform for the screening of pharmacological or genetic therapies.

In conclusion, the frameshift *Grin2b* variant studied here reveals a spectrum of molecular and phenotypic alterations, highlighting the complex consequences of *Grin2b* PTV. Neuron-specific regulatory mechanisms underscore the remarkable plasticity of the NMDAR system and the need to better understand the processes that shape functional outcomes. The comparison of the *Grin2b* PTV and the previously characterized *Grin2b* missense variants suggests that *Grin2b* haploinsufficiency may be better tolerated than the presence of functionally altered GluN2B subunits, with implications for the possibilities of gene therapy.

## Abbreviations

ABD: agonist-binding domain
AMPA: α-Amino-3-hydroxy-5-methyl-4-isoxazolepropionic acid
AMPAR: α-Amino-3-hydroxy-5-methyl-4-isoxazolepropionic acid receptor
ANOVA: analysis of variance
AP-5: D-(-)-2-amino-5-phosphonopentanoic acid
ASD: autism spectrum disorder
ATD: amino-terminal domain
β2M: beta-2-mikroglobulin
CN: copy number
COS-7 cell: African green monkey kidney fibroblast-like cell
CTD: C-terminal domain
DAPI: 4′,6-diamidino-2-phenylindole
DIV: days *in vitro*
DMEM: Dulbecco’s Modified Eagle Medium
EDTA: ethylenediaminetetraacetic acid
eEPSC: evoked excitatory postsynaptic current
EPM: elevated plus maze
ER: Endoplasmic reticulum
FBS: fetal bovine serum
FDR: false discovery rate
GABAAR: γ-aminobutyric acid type A receptor
GFAP: glial fibrillary acidic protein
GFP: enhanced green fluorescent protein
HEK293T: human embryonic kidney 293T
HBSS: Hank’s Balanced Salt Solution
HRP: horseradish peroxidase
IC_50_: half maximal inhibitory concentration
ID: intellectual disability
IVA: in-vivo assembly
LSD: least significant difference
MD: molecular dynamics
mEPSCs: miniature excitatory postsynaptic currents
M1–M4: membrane helices 1–4
NBQX: 2,3-dioxo-6-nitro-1,2,3,4-tetrahydrobenzo[f]quinoxaline-7-sulfonamide
NMD: nonsense-mediated decay
NMDA: N-methyl-D-aspartate
NMDAR: N-methyl-D-aspartate receptor
NOR: novel object recognition
PBS: phosphate-buffered saline
PFA: paraformaldehyde
PME: Particle Mesh Ewald
PPI: prepulse inhibition
PPR: paired-pulse ratio
PSD-95: postsynaptic density protein 95
PTV: protein-truncating variant
ROI: region of interest
SDC: sodium deoxycholate
SEM: standard error of the mean
TCEP: Tris(2-carboxyethyl) phosphine
TEAB: triethylammonium bicarbonate
TMD: transmembrane domain
TMTpro: tandem mass tags

## Acknowledgements

This work was supported by the Czech Science Foundation (GACR): 23-04922S, Charles University Grant Agency (GAUK): 318323 and Research Project of the CAS RVO: 67985823. We thank Mgr. Romana Markova (Institute of Physiology CAS, Prague) for excellent technical assistance.

## Author contributions

L.V. designed research; K.F., V.K., E.T., M.C.S., A.K.Z., K.H., B.H.K., V.A., M.D., F.E.S.A.R., J.P., A.B., and T.S. performed research; K.F., P.B., J.C., M.K., and L.V. analyzed data; T.S., and L.V. wrote the paper.

## Supplementary information

### Supplementary Tables

**Table S1.**
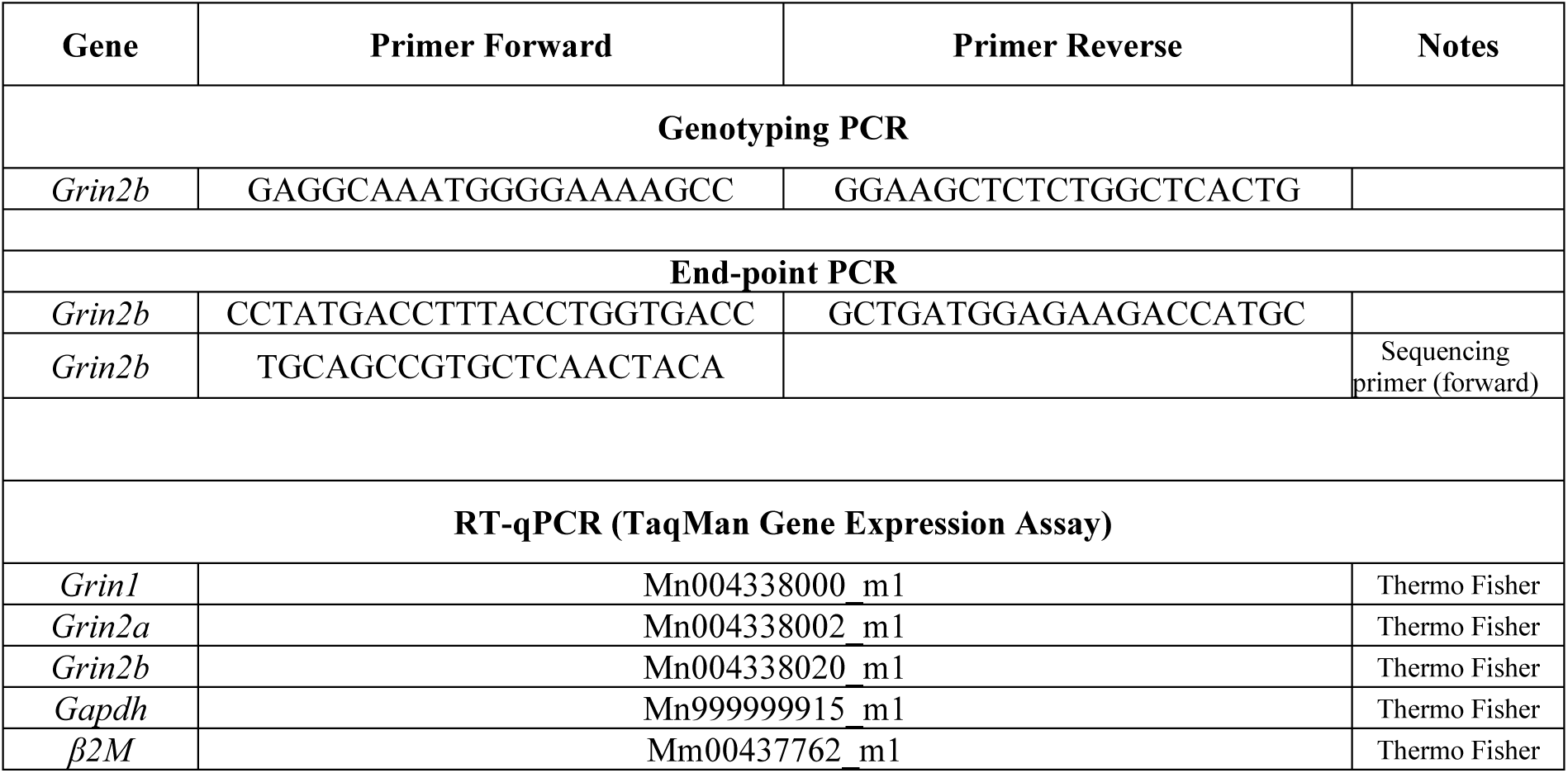
Sequences of primers used for PCR analyses and inventory numbers of TaqMan Gene Expresssion Assays used for qPCR.

**Table S2.**
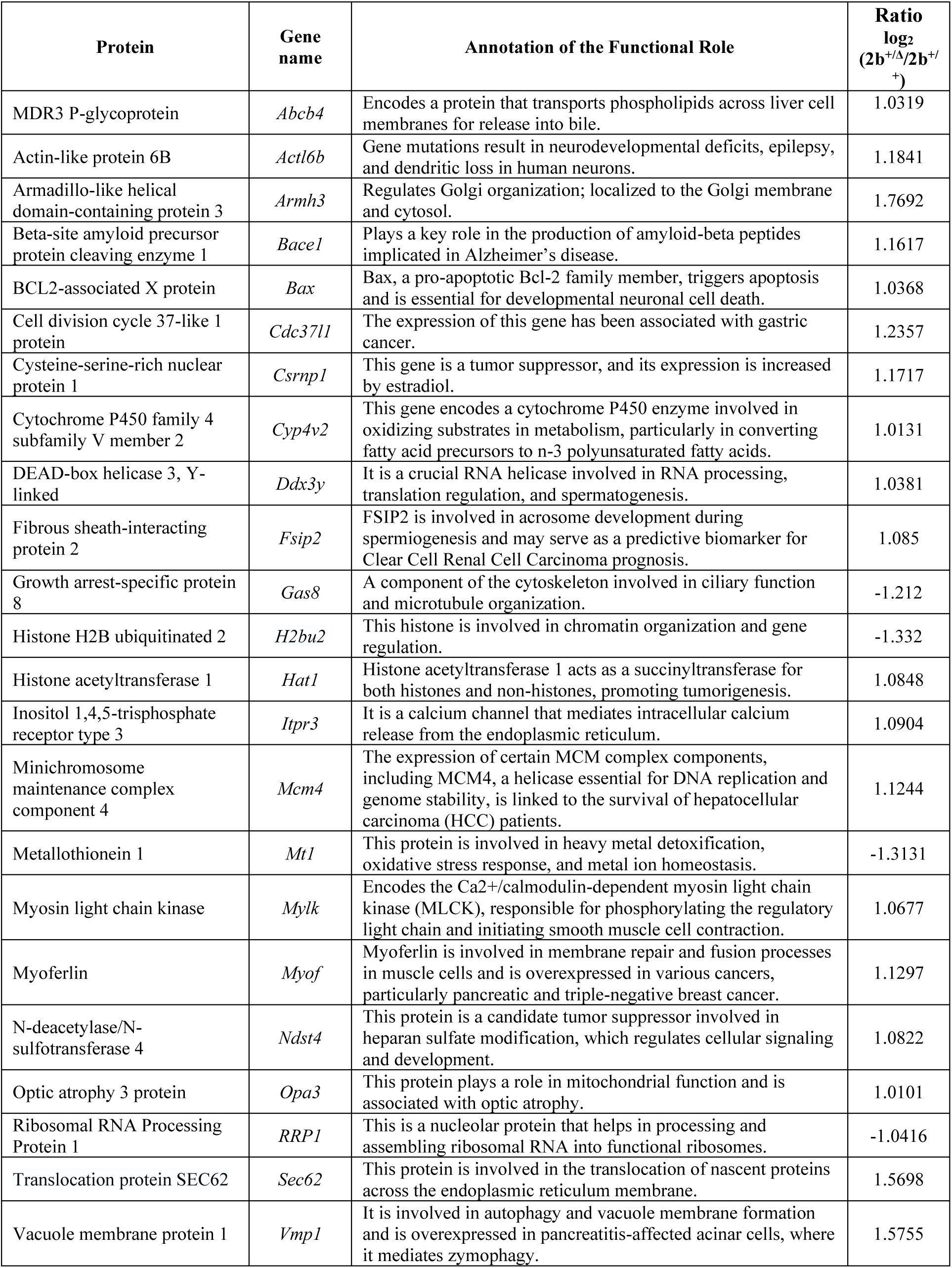
The list of significantly altered proteins and the annotation of their functional role.

**Table S3.**
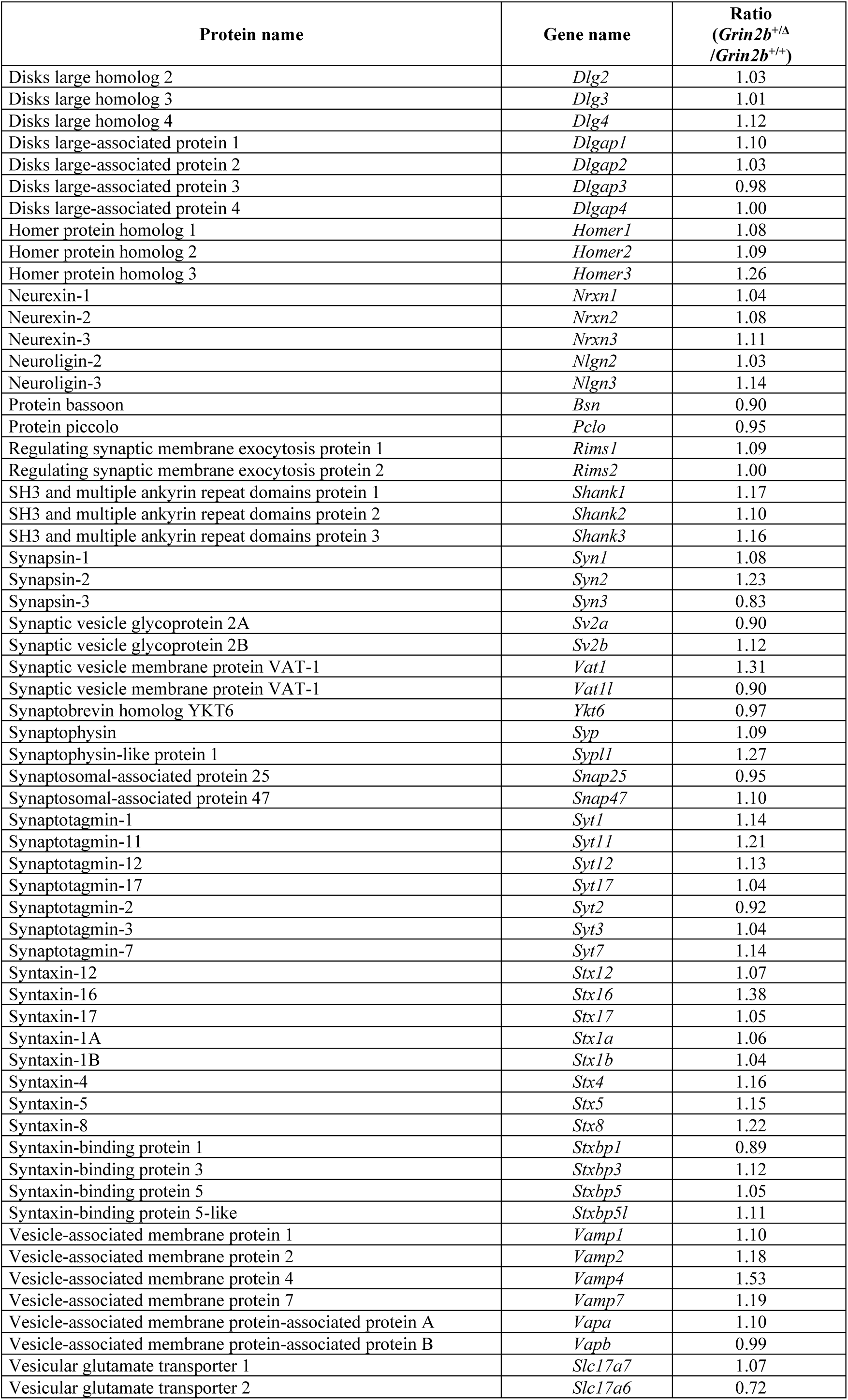
The list of selected synaptic proteins and their CN ratio in *Grin2b*^+/Δ^ relative to *Grin2b*^+/+^ hippocampal tissue.

**Table S4.**
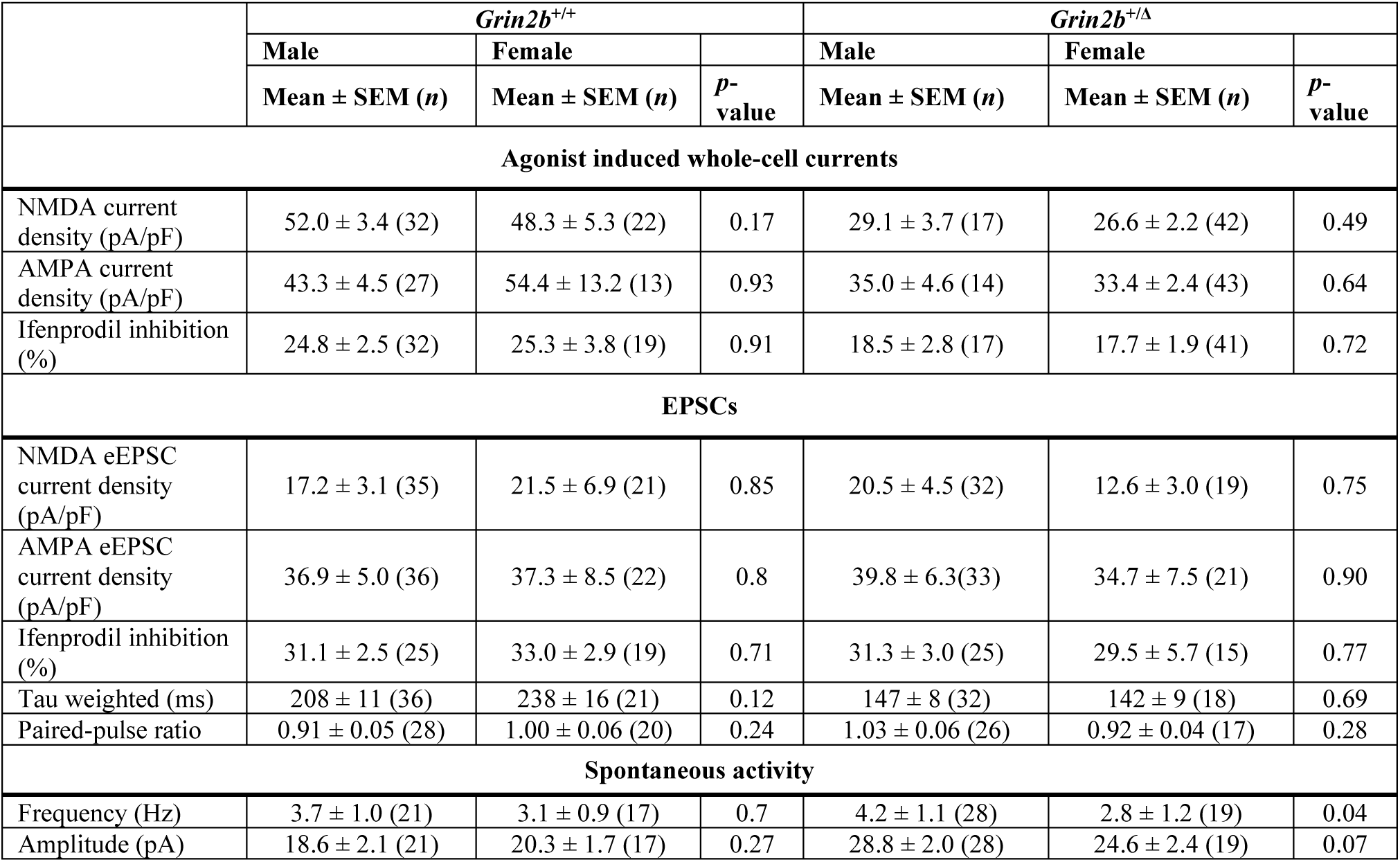
Data reported in Fig. 6-8, grouped by sex. Data from 13–43 neurons per group (*Grin2b*^+/+^ and *Grin2b*^+/Δ^) are presented as mean ± SEM. Genotype comparisons within each sex (male and female) were analyzed using Student’s t-test. Reported *p*-values indicate statistical differences between genotypes within each sex.

**Table S5.**
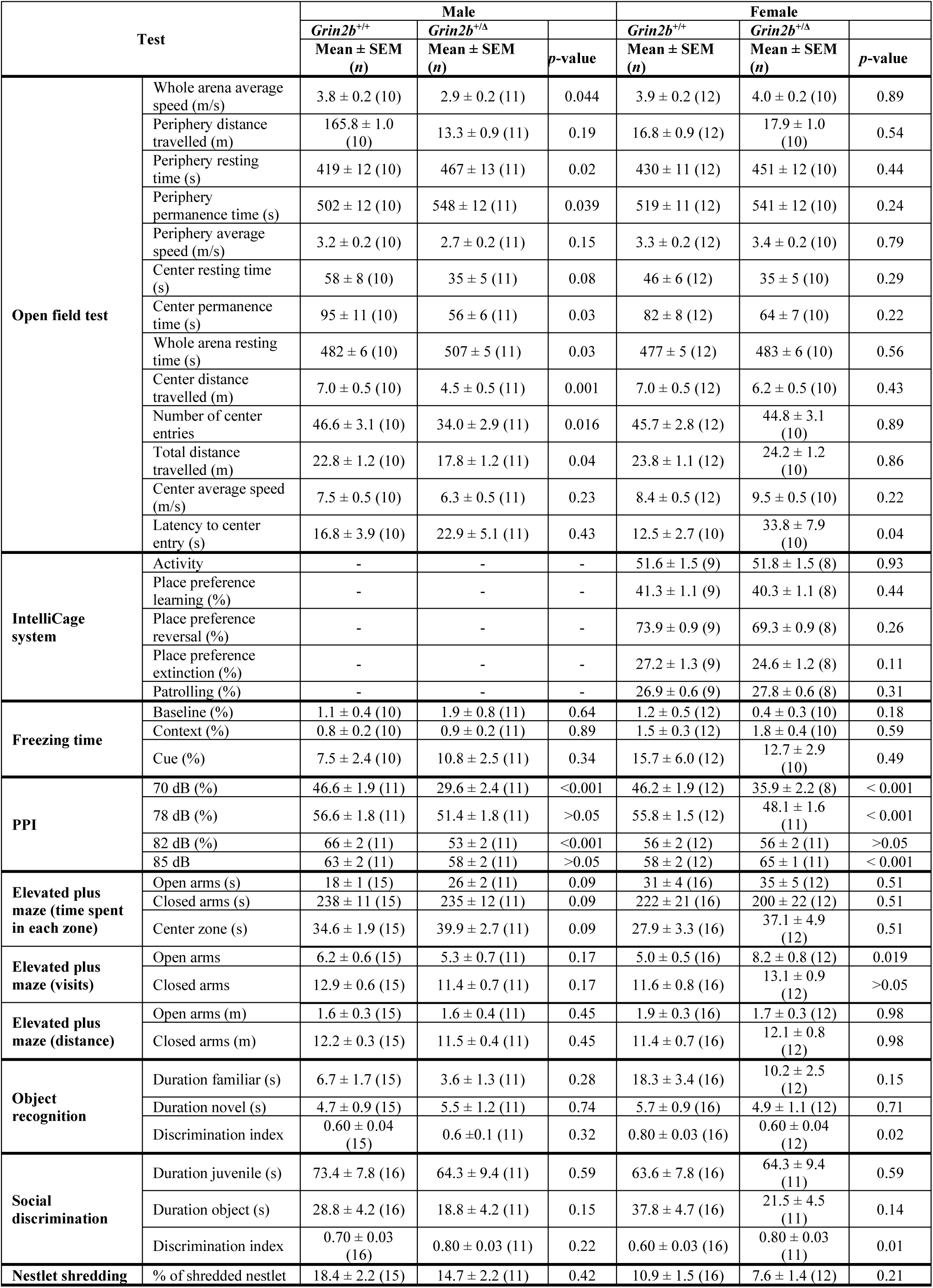
Numerical data underlying Fig. 9–10 and Fig. S3–S6. Data from 8–16 animals per group (*Grin2b*^+/+^ and *Grin2b*^+/Δ^ mice) are presented as mean ± SEM. Genotype comparisons within each sex (male and female) were made using ANOVA. Reported *p*-values indicate statistical differences between genotypes within each sex.

| Test |  | Male |  |  | Female |  |  |
| --- | --- | --- | --- | --- | --- | --- | --- |
|  |  | *Grin2b*<sup>+/+</sup> | *Grin2b*<sup>+/-</sup> |  | *Grin2b*<sup>+/+</sup> | *Grin2b*<sup>+/-</sup> |  |
| | | Mean ± SEM<br>(n) | Mean ± SEM<br>(n) | p-value | Mean ± SEM<br>(n) | Mean ± SEM<br>(n) | p-value |
| Open field test | Whole arena average speed (m/s) | 3.8 ± 0.2 (10) | 2.9 ± 0.2 (11) | 0.044 | 3.9 ± 0.2 (12) | 4.0 ± 0.2 (10) | 0.89 |
| | Periphery distance travelled (m) | 165.8 ± 1.0 (10) | 13.3 ± 0.9 (11) | 0.19 | 16.8 ± 0.9 (12) | 17.9 ± 1.0 (10) | 0.54 |
| | Periphery resting time (s) | 419 ± 12 (10) | 467 ± 13 (11) | 0.02 | 430 ± 11 (12) | 451 ± 12 (10) | 0.44 |
| | Periphery permanence time (s) | 502 ± 12 (10) | 548 ± 12 (11) | 0.039 | 519 ± 11 (12) | 541 ± 12 (10) | 0.24 |
| | Periphery average speed (m/s) | 3.2 ± 0.2 (10) | 2.7 ± 0.2 (11) | 0.15 | 3.3 ± 0.2 (12) | 3.4 ± 0.2 (10) | 0.79 |
| | Center resting time (s) | 58 ± 8 (10) | 35 ± 5 (11) | 0.08 | 46 ± 6 (12) | 35 ± 5 (10) | 0.29 |
| | Center permanence time (s) | 95 ± 11 (10) | 56 ± 6 (11) | 0.03 | 82 ± 8 (12) | 64 ± 7 (10) | 0.22 |
| | Whole arena resting time (s) | 482 ± 6 (10) | 507 ± 5 (11) | 0.03 | 477 ± 5 (12) | 483 ± 6 (10) | 0.56 |
| | Center distance travelled (m) | 7.0 ± 0.5 (10) | 4.5 ± 0.5 (11) | 0.001 | 7.0 ± 0.5 (12) | 6.2 ± 0.5 (10) | 0.43 |
| | Number of center entries | 46.6 ± 3.1 (10) | 34.0 ± 2.9 (11) | 0.016 | 45.7 ± 2.8 (12) | 44.8 ± 3.1 (10) | 0.89 |
| | Total distance travelled (m) | 22.8 ± 1.2 (10) | 17.8 ± 1.2 (11) | 0.04 | 23.8 ± 1.1 (12) | 24.2 ± 1.2 (10) | 0.86 |
| | Center average speed (m/s) | 7.5 ± 0.5 (10) | 6.3 ± 0.5 (11) | 0.23 | 8.4 ± 0.5 (12) | 9.5 ± 0.5 (10) | 0.22 |
| | Latency to center entry (s) | 16.8 ± 3.9 (10) | 22.9 ± 5.1 (11) | 0.43 | 12.5 ± 2.7 (10) | 33.8 ± 7.9 (10) | 0.04 |
| IntelliCage system | Activity | - | - | - | 51.6 ± 1.5 (9) | 51.8 ± 1.5 (8) | 0.93 |
| | Place preference learning (%) | - | - | - | 41.3 ± 1.1 (9) | 40.3 ± 1.1 (8) | 0.44 |
| | Place preference reversal (%) | - | - | - | 73.9 ± 0.9 (9) | 69.3 ± 0.9 (8) | 0.26 |
| | Place preference extinction (%) | - | - | - | 27.2 ± 1.3 (9) | 24.6 ± 1.2 (8) | 0.11 |
| | Patrolling (%) | - | - | - | 26.9 ± 0.6 (9) | 27.8 ± 0.6 (8) | 0.31 |
| Freezing time | Baseline (%) | 1.1 ± 0.4 (10) | 1.9 ± 0.8 (11) | 0.64 | 1.2 ± 0.5 (12) | 0.4 ± 0.3 (10) | 0.18 |
| | Context (%) | 0.8 ± 0.2 (10) | 0.9 ± 0.2 (11) | 0.89 | 1.5 ± 0.3 (12) | 1.8 ± 0.4 (10) | 0.59 |
| | Cue (%) | 7.5 ± 2.4 (10) | 10.8 ± 2.5 (11) | 0.34 | 15.7 ± 6.0 (12) | 12.7 ± 2.9 (10) | 0.49 |
| PPI | 70 dB (%) | 46.6 ± 1.9 (11) | 29.6 ± 2.4 (11) | <0.001 | 46.2 ± 1.9 (12) | 35.9 ± 2.2 (8) | < 0.001 |
| | 78 dB (%) | 56.6 ± 1.8 (11) | 51.4 ± 1.8 (11) | >0.05 | 55.8 ± 1.5 (12) | 48.1 ± 1.6 (11) | < 0.001 |
| | 82 dB (%) | 66 ± 2 (11) | 53 ± 2 (11) | <0.001 | 56 ± 2 (12) | 56 ± 2 (11) | >0.05 |
| | 85 dB | 63 ± 2 (11) | 58 ± 2 (11) | >0.05 | 58 ± 2 (12) | 65 ± 1 (11) | < 0.001 |
| Elevated plus maze (time spent in each zone) | Open arms (s) | 18 ± 1 (15) | 26 ± 2 (11) | 0.09 | 31 ± 4 (16) | 35 ± 5 (12) | 0.51 |
| | Closed arms (s) | 238 ± 11 (15) | 235 ± 12 (11) | 0.09 | 222 ± 21 (16) | 200 ± 22 (12) | 0.51 |
| | Center zone (s) | 34.6 ± 1.9 (15) | 39.9 ± 2.7 (11) | 0.09 | 27.9 ± 3.3 (16) | 37.1 ± 4.9 (12) | 0.51 |
| Elevated plus maze (visits) | Open arms | 6.2 ± 0.6 (15) | 5.3 ± 0.7 (11) | 0.17 | 5.0 ± 0.5 (16) | 8.2 ± 0.8 (12) | 0.019 |
| | Closed arms | 12.9 ± 0.6 (15) | 11.4 ± 0.7 (11) | 0.17 | 11.6 ± 0.8 (16) | 13.1 ± 0.9 (12) | >0.05 |
| Elevated plus maze (distance) | Open arms (m) | 1.6 ± 0.3 (15) | 1.6 ± 0.4 (11) | 0.45 | 1.9 ± 0.3 (16) | 1.7 ± 0.3 (12) | 0.98 |
| | Closed arms (m) | 12.2 ± 0.3 (15) | 11.5 ± 0.4 (11) | 0.45 | 11.4 ± 0.7 (16) | 12.1 ± 0.8 (12) | 0.98 |
| Object recognition | Duration familiar (s) | 6.7 ± 1.7 (15) | 3.6 ± 1.3 (11) | 0.28 | 18.3 ± 3.4 (16) | 10.2 ± 2.5 (12) | 0.15 |
| | Duration novel (s) | 4.7 ± 0.9 (15) | 5.5 ± 1.2 (11) | 0.74 | 5.7 ± 0.9 (16) | 4.9 ± 1.1 (12) | 0.71 |
| | Discrimination index | 0.60 ± 0.04 (15) | 0.6 ± 0.1 (11) | 0.32 | 0.80 ± 0.03 (16) | 0.60 ± 0.04 (12) | 0.02 |
| Social discrimination | Duration juvenile (s) | 73.4 ± 7.8 (16) | 64.3 ± 9.4 (11) | 0.59 | 63.6 ± 7.8 (16) | 64.3 ± 9.4 (11) | 0.59 |
| | Duration object (s) | 28.8 ± 4.2 (16) | 18.8 ± 4.2 (11) | 0.15 | 37.8 ± 4.7 (16) | 21.5 ± 4.5 (11) | 0.14 |
| | Discrimination index | 0.70 ± 0.03 (16) | 0.80 ± 0.03 (11) | 0.22 | 0.60 ± 0.03 (16) | 0.80 ± 0.03 (11) | 0.01 |
| Nestlet shredding | % of shredded nestlet | 18.4 ± 2.2 (15) | 14.7 ± 2.2 (11) | 0.42 | 10.9 ± 1.5 (16) | 7.6 ± 1.4 (12) | 0.21 |

### Supplementary Figures

**Figure S1.**
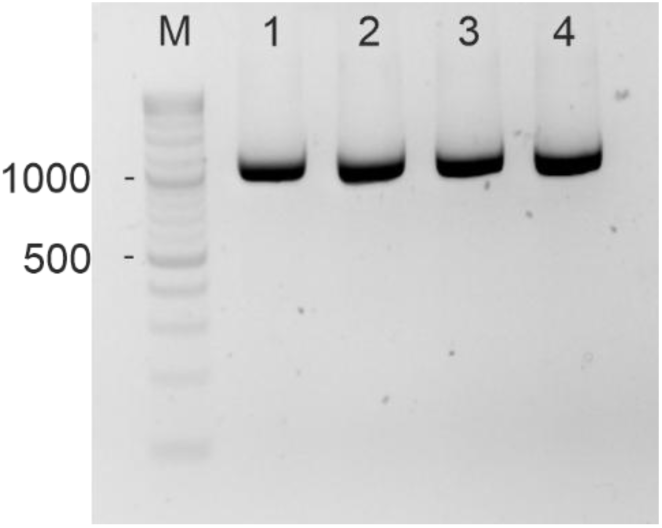
A representative agarose gel shows the amplification of a 1200 nt fragment of *Grin2b* mRNA. The mRNA was obtained from hippocampal tissue of *Grin2b*^+/Δ^ female mice at 4 months of age. The forward primer was located in exon 7 and the reverse primer in exon 13 (see Table S1 for sequences). For direct Sanger sequencing of the PCR amplicons, an additional forward primer located in exon 12 was used (see Table S1). M denotes the molecular weight marker; lanes 1–4 represent samples from four individual mice.

**Figure S2.**
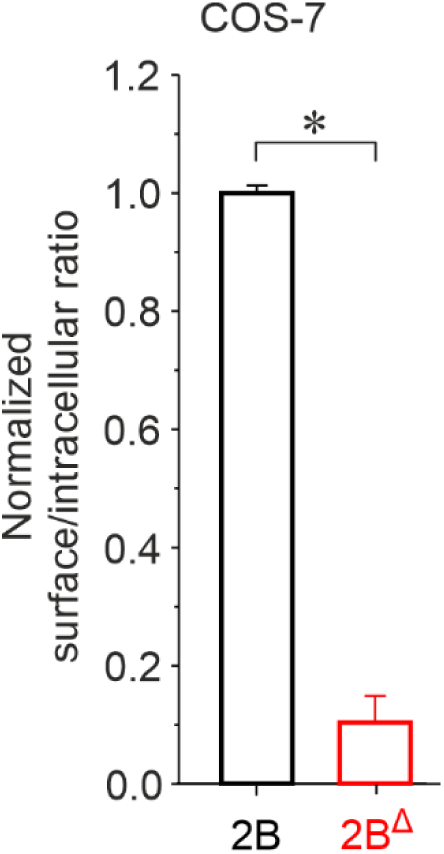
GluN2B^Δ^ subunit disrupts NMDAR surface expression in COS-7 cells. COS_-_7 cells were co-transfected with GluN1 and wild-type GluN2B-GFP (2B) or mutant GluN2B^Δ^-GFP (2B^Δ^) subunits. The graph shows the relative surface expression of GluN1/GFP-GluN2B and GluN1/GluN2B^Δ^-GFP receptors, quantified using a colorimetric assay. Data are presented as mean ± SEM. Asterisk (*) indicates a statistically significant difference (Student’s t-test; *n* = 4 independent experiments, with the relative surface expression measured in quadruplicate per experiment).

**Figure S3.**
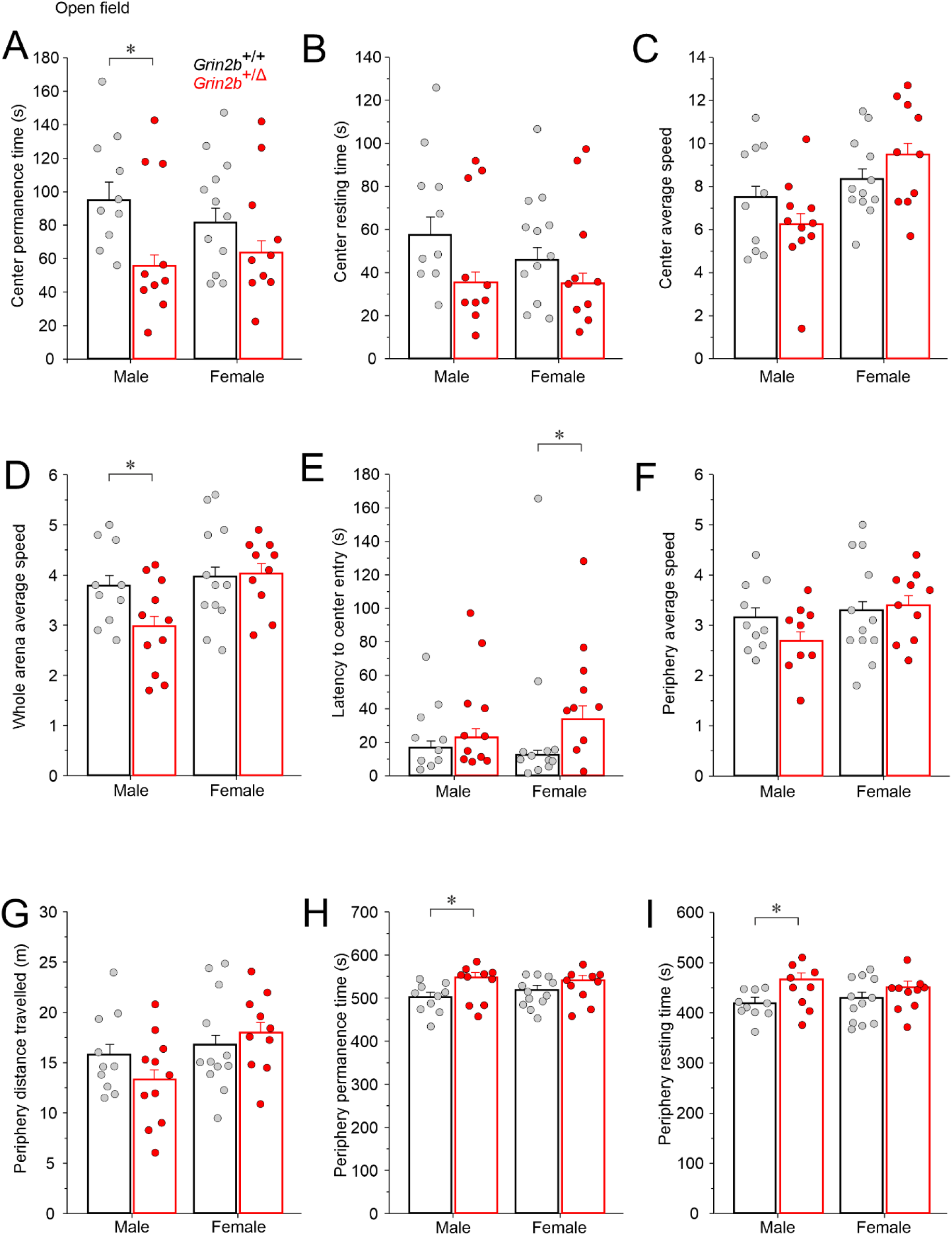
*Grin2b*^+/Δ^ mice display hypoactivity and increased anxiety. Supplementary data from the open field test. (A–I) Center permanence time, center resting time, center average speed, whole arena average speed, latency to first center entry, periphery average speed, periphery distance traveled, periphery permanence time, and periphery resting time were measured in male and female *Grin2b*^+/Δ^ and *Grin2b*^+/+^ mice. Data are presented as mean ± SEM. Number of animals *Grin2b*^+/+^/*Grin2b*^+/Δ^: *n* = 10/11 males and *n* = 12/10 females. Asterisks (*) indicate significant genotype differences (Student’s t-test). The numerical data underlying this figure are shown in Table S5.

**Figure S4.**
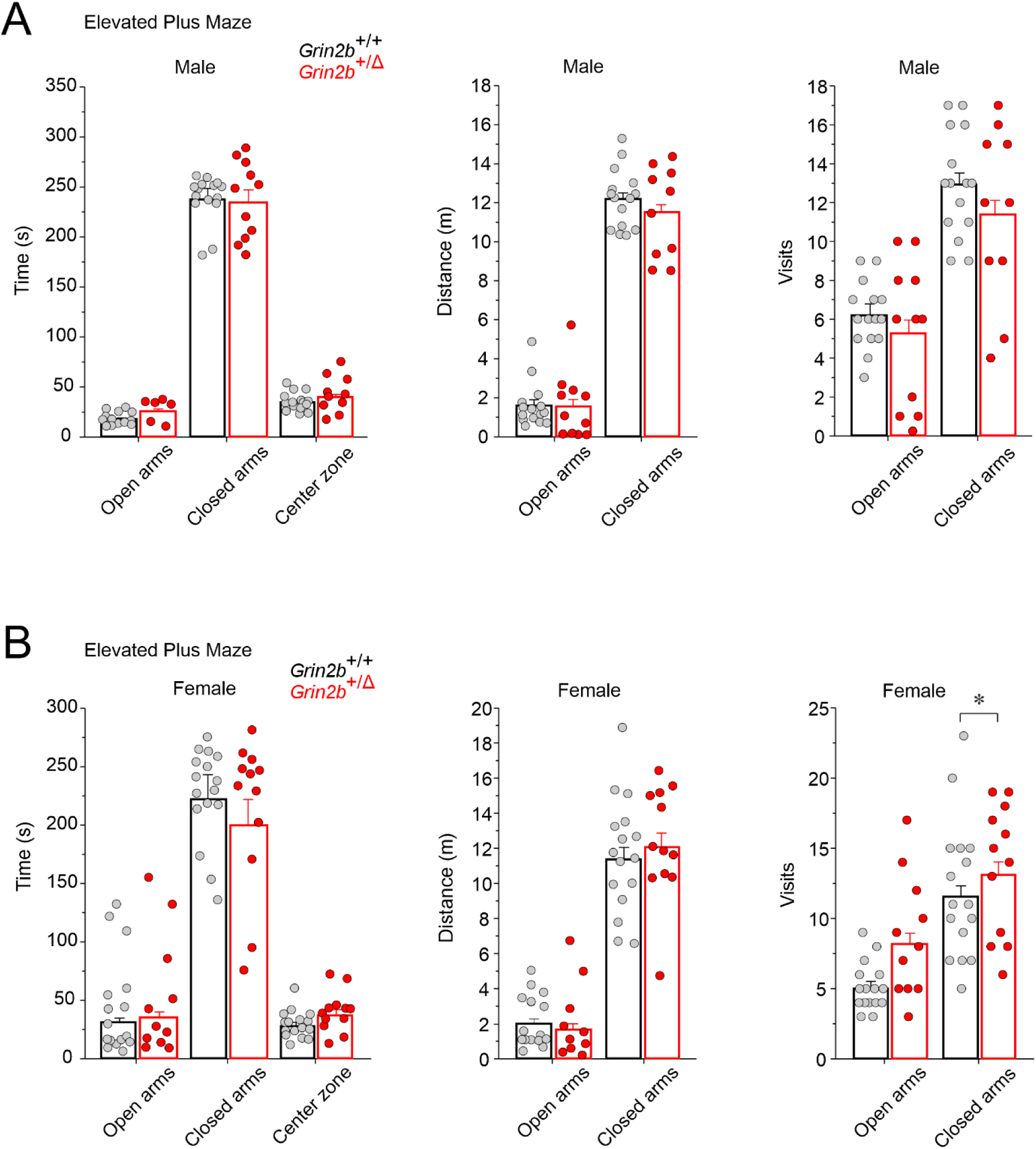
Elevated plus maze analysis of anxiety-like behavior in *Grin2b*^+/Δ^ mice. Bar graphs show measures of anxiety for male (A) and female (B) *Grin2b*^+/Δ^ mice, including time spent, distance traveled, and number of entries into the closed and open arms. Data are presented as mean ± SEM and were analyzed by ANOVA followed by post hoc LSD test. Number of animals *Grin2b*^+/+^/*Grin2b*^+/Δ^: *n* = 15/11 males and *n* = 16/12 females. Asterisk (*) indicates a significant genotype difference. The numerical data underlying this figure are shown in Table S5.

**Figure S5.**
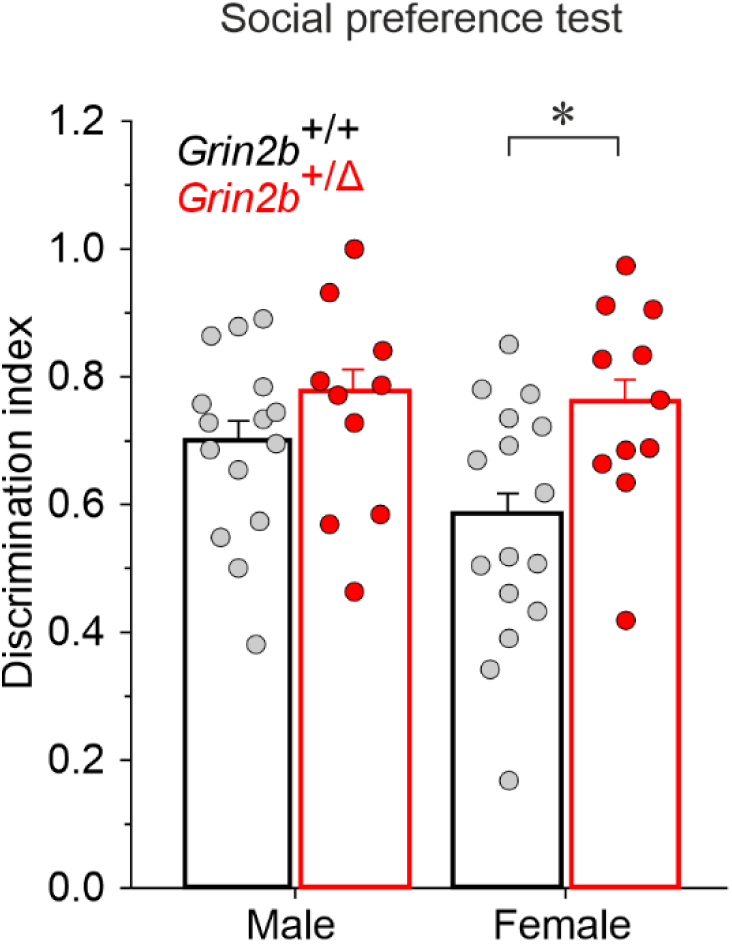
Social behavior in *Grin2b*^+/Δ^ mice. Social preference was assessed, with the discrimination index representing the time spent exploring a juvenile target relative to an object. Data are presented as mean ± SEM and were analyzed by Student’s t-test. Number of animals *Grin2b*^+/+^/*Grin2b*^+/Δ^: 16/11 males, 16/11 females. Asterisk (*) indicates a significant genotype difference. The numerical data underlying this figure are shown in Table S5.

**Figure S6.**
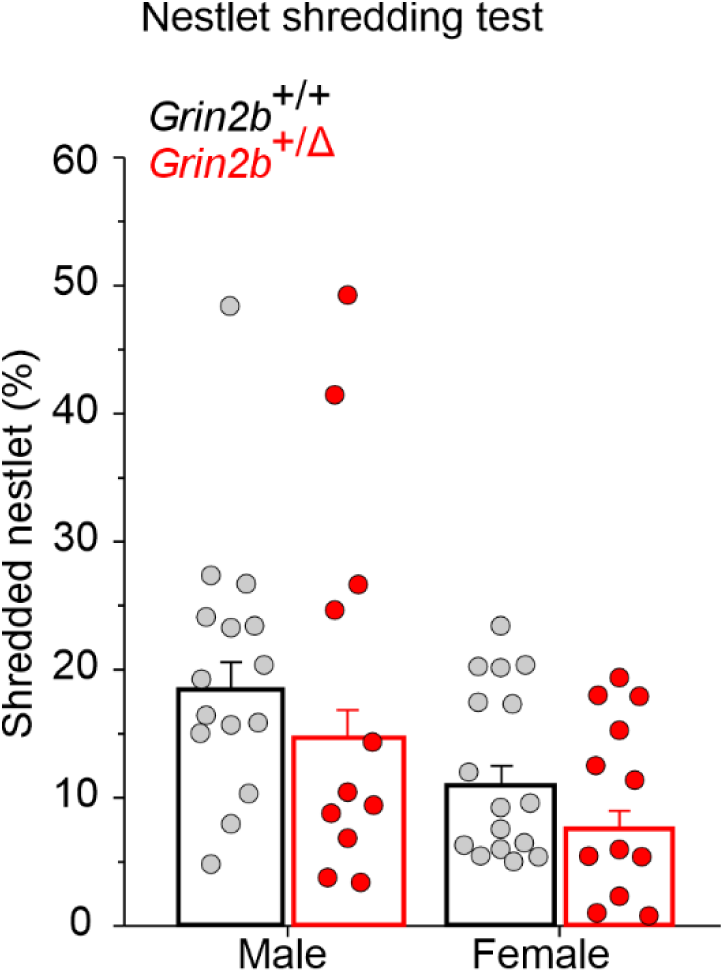
Repetitive behavior in *Grin2b*^+/Δ^ mice. Performance in the nestlet shredding test, measured as the % weight of unused nesting material (initial weight = 10 g). Data are presented as mean ± SEM and were analyzed by Student’s *t-test*. Number of animals *Grin2b*^+/+^/*Grin2b*^+/Δ^: *n* = 15/11 males, *n* = 16/12 females. The numerical data underlying this figure are provided in Table S5.

## References

Abraham MJ, Murtola T, Schulz R, Pall S, Smith JC, Hess B, Lindahl E (2015) GROMACS: High performance molecular simulations through multi-level parallelism from laptops to supercomputers. SoftwareX 1–2:19-25.

Al-Hallaq RA, Conrads TP, Veenstra TD, Wenthold RJ (2007) NMDA di-heteromeric receptor populations and associated proteins in rat hippocampus. J Neurosci 27:8334–8343.

Alvarez VA, Ridenour DA, Sabatini BL (2007) Distinct structural and ionotropic roles of NMDA receptors in controlling spine and synapse stability. J Neurosci 27:7365–7376.

Amin JB, Moody GR, Wollmuth LP (2021) From bedside-to-bench: What disease-associated variants are teaching us about the NMDA receptor. J Physiol 599:397–416.

Awadalla P et al. (2010) Direct measure of the de novo mutation rate in autism and schizophrenia cohorts. Am J Hum Genet 87:316–324.

Bar-Shira O, Maor R, Chechik G (2015) Gene Expression Switching of Receptor Subunits in Human Brain Development. Plos Comput Biol 11:e1004559.

Barria A, Malinow R (2002) Subunit-Specific NMDA Receptor Trafficking to Synapses. Neuron 35:345–353.

Benke TA, Park K, Krey I, Camp CR, Song R, Ramsey AJ, Yuan H, Traynelis SF, Lemke J (2021) Clinical and therapeutic significance of genetic variation in the GRIN gene family encoding NMDARs. Neuropharmacology 199:108805.

Benske TM, Mu TW, Wang YJ (2022) Protein quality control of N-methyl-D-aspartate receptors. Front Cell Neurosci 16:907560.

Borovska J, Vyklicky V, Stastna E, Kapras V, Slavikova B, Horak M, Chodounska H, Vyklicky L, Jr. (2012) Access of inhibitory neurosteroids to the NMDA receptor. Br J Pharmacol 166:1069–1083.

Candelas Serra M, Kuchtiak V, Kubik-Zahorodna A, Kysilov B, Fili K, Hrcka Krausova B, Abramova V, Dobrovolski M, Harant K, Bozikova P, Cerny J, Prochazka J, Kasparek P, Sedlacek R, Balik A, Smejkalova T, Vyklicky L (2024) Characterization of Mice Carrying a Neurodevelopmental Disease-Associated GluN2B(L825V) Variant. J Neurosci 44.

Cerny J, Bozikova P, Balik A, Marques SM, Vyklicky L (2019) NMDA Receptor Opening and Closing-Transitions of a Molecular Machine Revealed by Molecular Dynamics. Biomolecules 9.

de Visser JA, Hermisson J, Wagner GP, Ancel Meyers L, Bagheri-Chaichian H, Blanchard JL, Chao L, Cheverud JM, Elena SF, Fontana W, Gibson G, Hansen TF, Krakauer D, Lewontin RC, Ofria C, Rice SH, von Dassow G, Wagner A, Whitlock MC (2003) Perspective: Evolution and detection of genetic robustness. Evolution 57:1959–1972.

Downes JH, Hammond MW, Xydas D, Spencer MC, Becerra VM, Warwick K, Whalley BJ, Nasuto SJ (2012) Emergence of a small-world functional network in cultured neurons. Plos Comput Biol 8:e1002522.

Endele S et al. (2010) Mutations in GRIN2A and GRIN2B encoding regulatory subunits of NMDA receptors cause variable neurodevelopmental phenotypes. Nat Genet 42:1021–1026.

Erreger K, Geballe MT, Kristensen A, Chen PE, Hansen KB, Lee CJ, Yuan H, Le P, Lyuboslavsky PN, Micale N, Jorgensen L, Clausen RP, Wyllie DJ, Snyder JP, Traynelis SF (2007) Subunit-specific agonist activity at NR2A-, NR2B-, NR2C-, and NR2D-containing N-methyl-D-aspartate glutamate receptors. Mol Pharmacol 72:907–920.

Essmann U, Perera L, Berkowitz ML, Darden T, Lee H, Pedersen LG (1995) A Smooth Particle Mesh Ewald Method. J Chem Phys 103:8577–8593.

Flint AC, Maisch US, Weishaupt JH, Kriegstein AR, Monyer H (1997) NR2A Subunit Expression Shortens NMDA Receptor Synaptic Currents in Developing Neocortex. J Neurosci 17:2469–2476.

Forrest D, Yuzaki M, Soares HD, Ng L, Luk DC, Sheng M, Stewart CL, Morgan JI, Connor JA, Curran T (1994) Targeted disruption of NMDA receptor 1 gene abolishes NMDA response and results in neonatal death. Neuron 13:325–338.

Garcia-Nafria J, Watson JF, Greger IH (2016) IVA cloning: A single-tube universal cloning system exploiting bacterial In Vivo Assembly. Sci Rep 6:27459.

Gray JA, Shi Y, Usui H, During MJ, Sakimura K, Nicoll RA (2011) Distinct modes of AMPA receptor suppression at developing synapses by GluN2A and GluN2B: single-cell NMDA receptor subunit deletion in vivo. Neuron 71:1085–1101.

Groc L, Heine M, Cousins SL, Stephenson FA, Lounis B, Cognet L, Choquet D (2006) NMDA receptor surface mobility depends on NR2A-2B subunits. Proc Natl Acad Sci U S A 103:18769–18774.

Hall BJ, Ripley B, Ghosh A (2007) NR2B signaling regulates the development of synaptic AMPA receptor current. J Neurosci 27:13446–13456.

Hansen KB, Ogden KK, Yuan H, Traynelis SF (2014) Distinct functional and pharmacological properties of Triheteromeric GluN1/GluN2A/GluN2B NMDA receptors. Neuron 81:1084–1096.

Hansen KB, Wollmuth LP, Bowie D, Furukawa H, Menniti FS, Sobolevsky AI, Swanson GT, Swanger SA, Greger IH, Nakagawa T, McBain CJ, Jayaraman V, Low CM, Dell’Acqua ML, Diamond JS, Camp CR, Perszyk RE, Yuan H, Traynelis SF (2021) Structure, Function, and Pharmacology of Glutamate Receptor Ion Channels. Pharmacol Rev 73:298–487.

Hanson JE, Yuan H, Perszyk RE, Banke TG, Xing H, Tsai MC, Menniti FS, Traynelis SF (2024) Therapeutic potential of N-methyl-D-aspartate receptor modulators in psychiatry. Neuropsychopharmacology 49:51–66.

Hassan S, Lopez G, Stubblefield BK, Tayebi N, Sidransky E (2018) Alleles with more than one mutation can complicate genotype/phenotype studies in Mendelian disorders: Lessons from Gaucher disease. Mol Genet Metab 125:1–3.

Hawkins LM, Prybylowski K, Chang K, Moussan C, Stephenson FA, Wenthold RJ (2004) Export from the endoplasmic reticulum of assembled N-methyl-D-aspartic acid receptors is controlled by a motif in the C terminus of the NR2 subunit. Journal of Biological Chemistry 279:28903–28910.

Hestrin S (1992) Developmental regulation of NMDA receptor-mediated synaptic currents at a central synapse. Nature 357:686–689.

Horak M, Chang K, Wenthold RJ (2008) Masking of the endoplasmic reticulum retention signals during assembly of the NMDA receptor. J Neurosci 28:3500–3509.

Howe MP, Patani R (2023) Nonsense-mediated mRNA decay in neuronal physiology and neurodegeneration. Trends in Neurosciences 46:879–892.

Hu C, Chen W, Myers SJ, Yuan H, Traynelis SF (2016) Human GRIN2B variants in neurodevelopmental disorders. J Pharmacol Sci 132:115–121.

Jo S, Kim T, Iyer VG, Im W (2008) CHARMM-GUI: a web-based graphical user interface for CHARMM. J Comput Chem 29:1859–1865.

Karakas E, Furukawa H (2014) Crystal structure of a heterotetrameric NMDA receptor ion channel. Science 344:992–997.

Kaur M, Esau L (2015) Two-step protocol for preparing adherent cells for high-throughput flow cytometry. Biotechniques 59:119–126.

Kelsch W, Li ZJ, Wieland S, Senkov O, Herb A, Göngrich C, Monyer H (2014) GluN2B-Containing NMDA Receptors Promote Glutamate Synapse Development in Hippocampal Interneurons. Journal of Neuroscience 34:16022–16030.

Klauda JB, Venable RM, Freites JA, O’Connor JW, Tobias DJ, Mondragon-Ramirez C, Vorobyov I, MacKerell AD, Jr., Pastor RW (2010) Update of the CHARMM all-atom additive force field for lipids: validation on six lipid types. J Phys Chem B 114:7830–7843.

Korinek M, Serra MC, Abdel Rahman F, Dobrovolski M, Kuchtiak V, Abramova V, Fili K, Tomovic E, Hrcka Krausova B, Krusek J, Cerny J, Vyklicky L, Balik A, Smejkalova TT (2024) Disease-Associated Variants in GRIN1, GRIN2A and GRIN2B genes: Insights into NMDA Receptor Structure, Function, and Pathophysiology. Physiol Res.

Kuchtiak V, Smejkalova T, Horak M, Vyklicky L, Balik A (2024) Analysis of Surface Expression of NMDAR Subunits in Primary Hippocampal Neurons. Methods Mol Biol 2799:29–46.

Kurosaki T, Popp MW, Maquat LE (2019) Quality and quantity control of gene expression by nonsense-mediated mRNA decay. Nat Rev Mol Cell Bio 20:406–420.

Kutsuwada T, Sakimura K, Manabe T, Takayama C, Katakura N, Kushiya E, Natsume R, Watanabe M, Inoue Y, Yagi T, Aizawa S, Arakawa M, Takahashi T, Nakamura Y, Mori H, Mishina M (1996) Impairment of suckling response, trigeminal neuronal pattern formation, and hippocampal LTD in NMDA receptor epsilon 2 subunit mutant mice. Neuron 16:333–344.

Kysilov B, Kuchtiak V, Krausova BH, Balik A, Korinek M, Fili K, Dobrovolski M, Abramova V, Chodounska H, Kudova E, Bozikova P, Cerny J, Smejkalova T, Vyklicky L (2024) Disease-associated nonsense and frame-shift variants resulting in the truncation of the GluN2A or GluN2B C-terminal domain decrease NMDAR surface expression and reduce potentiating effects of neurosteroids. Cell Mol Life Sci 81.

Laube B, Kuhse J, Betz H (1998) Evidence for a tetrameric structure of recombinant NMDA receptors. J Neurosci 18:2954–2961.

Lee G, Zhou Y (2019) NMDAR Hypofunction Animal Models of Schizophrenia. Front Mol Neurosci 12.

Lee CH, Lu W, Michel JC, Goehring A, Du J, Song X, Gouaux E (2014) NMDA receptor structures reveal subunit arrangement and pore architecture. Nature 511:191–197.

Lee J, Cheng X, Swails JM, Yeom MS, Eastman PK, Lemkul JA, Wei S, Buckner J, Jeong JC, Qi Y, Jo S, Pande VS, Case DA, Brooks CL, 3rd, MacKerell AD, Jr., Klauda JB, Im W (2016) CHARMM-GUI Input Generator for NAMD, GROMACS, AMBER, OpenMM, and CHARMM/OpenMM Simulations Using the CHARMM36 Additive Force Field. J Chem Theory Comput 12:405–413.

Lee PJ, Yang S, Sun Y, Guo JU (2021) Regulation of nonsense-mediated mRNA decay in neural development and disease. J Mol Cell Biol 13:269–281.

Livak KJ, Schmittgen TD (2001) Analysis of relative gene expression data using real-time quantitative PCR and the 2(-Delta Delta C(T)) method. Methods 25:402–408.

López-Rivera JA, Pérez-Palma E, Symonds J, Lindy AS, McKnight DA, Leu C, Zuberi S, Brunklaus A, Moller RS, Lal D (2020) A catalogue of new incidence estimates of monogenic neurodevelopmental disorders caused by variants. Brain 143:1099–1105.

Luo J, Wang Y, Yasuda RP, Dunah AW, Wolfe BB (1997) The majority of N-methyl-D-aspartate receptor complexes in adult rat cerebral cortex contain at least three different subunits (NR1/NR2A/NR2B). Mol Pharmacol 51:79–86.

Luo JH, Fu ZY, Losi G, Kim BG, Prybylowski K, Vissel B, Vicini S (2002) Functional expression of distinct NMDA channel subunits tagged with green fluorescent protein in hippocampal neurons in culture. Neuropharmacology 42:306–318.

McKay S, Ryan TJ, McQueen J, Indersmitten T, Marwick KFM, Hasel P, Kopanitsa MV, Baxter PS, Martel MA, Kind PC, Wyllie DJA, O’Dell TJ, Grant SGN, Hardingham GE, Komiyama NH (2018) The Developmental Shift of NMDA Receptor Composition Proceeds Independently of GluN2 Subunit-Specific GluN2 C-Terminal Sequences. Cell Rep 25:841–851 e844.

Monyer H, Burnashev N, Laurie DJ, Sakmann B, Seeburg PH (1994) Developmental and regional expression in the rat brain and functional properties of four NMDA receptors. Neuron 12:529–540.

Monyer H, Sprengel R, Schoepfer R, Herb A, Higuchi M, Lomeli H, Burnashev N, Sakmann B, Seeburg PH (1992) Heteromeric NMDA receptors: molecular and functional distinction of subtypes. Science 256:1217–1221.

Perry W, Minassian A, Lopez B, Maron L, Lincoln A (2007) Sensorimotor gating deficits in adults with autism. Biol Psychiatry 61:482–486.

Perszyk RE, Kristensen AS, Lyuboslavsky P, Traynelis SF (2021) Three-dimensional missense tolerance ratio analysis. Genome Res 31:1447–1461.

Petralia RS, Wang YX, Hua F, Yi Z, Zhou A, Ge L, Stephenson FA, Wenthold RJ (2010) Organization of NMDA receptors at extrasynaptic locations. Neuroscience 167:68–87.

Platzer K, Lemke JR (1993) GRIN2B-Related Neurodevelopmental Disorder. In: GeneReviews((R)) (Adam MP, Mirzaa GM, Pagon RA, Wallace SE, Bean LJH, Gripp KW, Amemiya A, eds). Seattle (WA).

Platzer K et al. (2017) GRIN2B encephalopathy: novel findings on phenotype, variant clustering, functional consequences and treatment aspects. Journal of Medical Genetics 54:460–470.

Prybylowski K, Fu ZY, Losi G, Hawkins LM, Luo JH, Chang K, Wenthold RJ, Vicini S (2002) Relationship between availability of NMDA receptor subunits and their expression at the synapse. Journal of Neuroscience 22:8902–8910.

Quatraccioni A, Cases-Cunillera S, Balagura G, Coleman M, Rossini L, Mills JD, Casillas-Espinosa PM, Moshe SL, Sankar R, Baulac S, Noebels JL, Auvin S, O’Brien TJ, Henshall DC, Akman O, Galanopoulou AS (2024) WONOEP appraisal: Genetic insights into early onset epilepsies. Epilepsia 65:3138–3154.

Rampado R, Giordano F, Moracci L, Crotti S, Caliceti P, Agostini M, Taraballi F (2022) Optimization of a detergent-based protocol for membrane proteins purification from mammalian cells. J Pharm Biomed Anal 219:114926.

Rauner C, Kohr G (2011) Triheteromeric NR1/NR2A/NR2B receptors constitute the major N-methyl-D-aspartate receptor population in adult hippocampal synapses. J Biol Chem 286:7558–7566.

Sabo SL, Lahr JM, Offer M, Weekes A, Sceniak MP (2022) GRIN2B-related neurodevelopmental disorder: current understanding of pathophysiological mechanisms. Front Synaptic Neurosci 14:1090865.

Sali A, Blundell TL (1993) Comparative protein modelling by satisfaction of spatial restraints. J Mol Biol 234:779–815.

Santos-Gomez A, Miguez-Cabello F, Garcia-Recio A, Locubiche-Serra S, Garcia-Diaz R, Soto-Insuga V, Guerrero-Lopez R, Julia-Palacios N, Ciruela F, Garcia-Cazorla A, Soto D, Olivella M, Altafaj X (2021) Disease-associated GRIN protein truncating variants trigger NMDA receptor loss-of-function. Hum Mol Genet 29:3859–3871.

Sheng M, Cummings J, Roldan LA, Jan YN, Jan LY (1994) Changing subunit composition of heteromeric NMDA receptors during development of rat cortex. Nature 368:144–147.

Shin W, Kim K, Serraz B, Cho YS, Kim D, Kang M, Lee EJ, Lee H, Bae YC, Paoletti P, Kim E (2020) Early correction of synaptic long-term depression improves abnormal anxiety-like behavior in adult GluN2B-C456Y-mutant mice. PLoS Biol 18:e3000717.

Schauder DM, Kuybeda O, Zhang J, Klymko K, Bartesaghi A, Borgnia MJ, Mayer ML, Subramaniam S (2013) Glutamate receptor desensitization is mediated by changes in quaternary structure of the ligand binding domain. Proc Natl Acad Sci U S A 110:5921–5926.

Silverman JL, Yang M, Lord C, Crawley JN (2010) Behavioural phenotyping assays for mouse models of autism. Nat Rev Neurosci 11:490–502.

Smejkalova T, Korinek M, Krusek J, Hrcka Krausova B, Candelas Serra M, Hajdukovic D, Kudova E, Chodounska H, Vyklicky L (2021) Endogenous neurosteroids pregnanolone and pregnanolone sulfate potentiate presynaptic glutamate release through distinct mechanisms. Br J Pharmacol 178:3888–3904.

Sonzogni M, Zhai P, Mientjes EJ, van Woerden GM, Elgersma Y (2020) Assessing the requirements of prenatal UBE3A expression for rescue of behavioral phenotypes in a mouse model for Angelman syndrome. Mol Autism 11:70.

Strehlow V et al. (2019) GRIN2A-related disorders: genotype and functional consequence predict phenotype. Brain 142:80–92.

Swerdlow NR, Benbow CH, Zisook S, Geyer MA, Braff DL (1993) A preliminary assessment of sensorimotor gating in patients with obsessive compulsive disorder. Biol Psychiatry 33:298–301.

Takasaki Y et al. (2016) Mutation screening of GRIN2B in schizophrenia and autism spectrum disorder in a Japanese population. Sci Rep 6:33311.

Takeuchi T, Kiyama Y, Nakamura K, Tsujita M, Matsuda I, Mori H, Munemoto Y, Kuriyama H, Natsume R, Sakimura K, Mishina M (2001) Roles of the glutamate receptor epsilon2 and delta2 subunits in the potentiation and prepulse inhibition of the acoustic startle reflex. Eur J Neurosci 14:153–160.

Tarabeux J et al. (2011) Rare mutations in N-methyl-D-aspartate glutamate receptors in autism spectrum disorders and schizophrenia. Translational Psychiatry 1:e55.

Taylor SC, Nadeau K, Abbasi M, Lachance C, Nguyen M, Fenrich J (2019) The Ultimate qPCR Experiment: Producing Publication Quality, Reproducible Data the First Time. Trends Biotechnol 37:761–774.

Tovar KR, McGinley MJ, Westbrook GL (2013) Triheteromeric NMDA receptors at hippocampal synapses. J Neurosci 33:9150–9160.

Tyanova S, Cox J (2018) Perseus: A Bioinformatics Platform for Integrative Analysis of Proteomics Data in Cancer Research. Methods Mol Biol 1711:133–148.

UniProt C (2021) UniProt: the universal protein knowledgebase in 2021. Nucleic Acids Res 49:D480–D489.

Velasco CD, Llobet A (2020) Synapse elimination activates a coordinated homeostatic presynaptic response in an autaptic circuit. Commun Biol 3:260.

Vicini S, Wang JF, Li JH, Zhu WJ, Wang YH, Luo JH, Wolfe BB, Grayson DR (1998) Functional and pharmacological differences between recombinant N-methyl-D-aspartate receptors. J Neurophysiol 79:555–566.

von Engelhardt J, Doganci B, Jensen V, Hvalby O, Gongrich C, Taylor A, Barkus C, Sanderson DJ, Rawlins JN, Seeburg PH, Bannerman DM, Monyer H (2008) Contribution of hippocampal and extra-hippocampal NR2B-containing NMDA receptors to performance on spatial learning tasks. Neuron 60:846–860.

Wang X, Dong Q, Chen G, Zhang J, Liu Y, Cai Y (2022) Frameshift and wild-type proteins are often highly similar because the genetic code and genomes were optimized for frameshift tolerance. BMC Genomics 23:416.

Webb B, Sali A (2014) Comparative Protein Structure Modeling Using MODELLER. Curr Protoc Bioinformatics 47:5 6 1–32.

Wenthold RJ, Prybylowski K, Standley S, Sans N, Petralia RS (2003) Trafficking of NMDA receptors. Annu Rev Pharmacol Toxicol 43:335–358.

Williams K (1993) Ifenprodil discriminates subtypes of the N-methyl-D-aspartate receptor: selectivity and mechanisms at recombinant heteromeric receptors. Mol Pharmacol 44:851–859.

Wisniewski JR, Hein MY, Cox J, Mann M (2014) A “Proteomic Ruler” for Protein Copy Number and Concentration Estimation without Spike-in Standards. Molecular & Cellular Proteomics 13:3497–3506.

Wyllie DJ, Livesey MR, Hardingham GE (2013) Influence of GluN2 subunit identity on NMDA receptor function. Neuropharmacology 74:4–17.

