## Supplementary Information for "Phenotype of mice carrying an NMDA receptor GluN2B protein-truncating variant associated with intellectual disability"

**Title:**

**Abbreviated title**: Phenotype of *Grin2b*^+/Δ^ mice

**Author listing:**

Klevinda Fili^1,4^, Viktor Kuchtiak^1,5^, Eni Tomovic^1,5^, Miriam Candelas Serra^1^, Agnieszka Kubik-Zahorodna^2^, Karel Harant^3^, Paulina Bozikova^1^, Jiri Cerny^1^, Miloslav Korinek^1^, Barbora Hrcka Krausova^1^, Vera Abramova^1,4^, Mark Dobrovolski^1,4^, Fatma Elzahraa S. Abdel Rahman^1,5^, Jan Prochazka^2^, Ales Balik^1^, Tereza Smejkalova^1*^, Ladislav Vyklicky^1*^

^1^Institute of Physiology CAS, 14220, Prague, Czech Republic

^2^Czech Centre for Phenogenomics, Institute of Molecular Genetics of the Czech Academy of Sciences, 25050 Vestec, Czech Republic

^3^Proteomics Core Facility, Faculty of Science, Charles University, Biocev, 25050 Vestec, Czech Republic

^4^Third Faculty of Medicine, Charles University, 10000 Prague, Czech Republic

^5^Faculty of Science, Charles University, Albertov 2038, 128 00 Prague 2, Czech Republic

^*^Co-corresponding authors: Prof. Ladislav Vyklicky, MD, PhD

Institute of Physiology, Czech Academy of Sciences

Videnska 1083, 142 00 Prague 4, Czech Republic

Phone: (+420) 241 062 450


Tereza Smejkalova, PhD

Institute of Physiology, Czech Academy of Sciences

Videnska 1083, 142 00 Prague 4, Czech Republic

Phone: (+420) 241 062 543

**Supplementary Tables:**

**Table S1 |** Sequences of primers used for PCR analysis and inventory numbers of TaqMan Gene Expresssion Assays used for qPCR.

**Table S2 |** The list of significantly altered proteins and annotation of their functional role.

**Table S3 |** The list of selected synaptic proteins and their CN ratio assessed in *Grin2b*^+/Δ^ relative to *Grin2b*^+/+^ hippocampal tissue.

**Table S4 |** Data reported in Fig. 6–8, grouped by sex.

**Table S5 |** Numerical data underlying Fig. 9–10 and Fig. S3–S6.

**Supplementary Figures:**

**Figure S1 |** A representative agarose gel shows amplification of a 1200 nt fragment of *Grin2b* mRNA.

**Figure S2 |** The GluN2B^Δ^ subunit disrupts NMDAR surface expression in COS-7 cells.

**Figure S3 |** *Grin2b*^+/Δ^ **mice display hypoactivity and increased anxiety. Supplementary data from the open field test.**

**Figure S4 | Elevated plus maze analysis of anxiety-like behavior in** *Grin2b*^+/Δ^ **mice.**

**Figure S5 | Social behavior in** *Grin2b*^+/Δ^ **mice.**

**Figure S6 | Repetitive behavior in** *Grin2b*^+/Δ^ **mice.**

### Supplementary Tables

| **Gene** | **Primer Forward** | **Primer Reverse** | **Notes** |
| --- | --- | --- | --- |
| **Genotyping PCR** | | | |
| *Grin2b* | GAGGCAAATGGGGAAAAGCC | GGAAGCTCTCTGGCTCACTG |  |
| **End-point PCR** | | | |
| *Grin2b* | CCTATGACCTTTACCTGGTGACC | GCTGATGGAGAAGACCATGC |  |
| *Grin2b* | TGCAGCCGTGCTCAACTACA |  | Sequencing primer (forward) |
| **RT-qPCR (TaqMan Gene Expression Assay)** | | | |
| *Grin1* | Mn004338000_m1 | | Thermo Fisher |
| *Grin2a* | Mn004338002_m1 | | Thermo Fisher |
| *Grin2b* | Mn004338020_m1 | | Thermo Fisher |
| *Gapdh* | Mn999999915_m1 | | Thermo Fisher |
| *β2M* | Mm00437762_m1 | | Thermo Fisher |

**Table S1 | Sequences of primers used for PCR analyses and inventory numbers of TaqMan Gene Expresssion Assays used for qPCR.**

| **Protein** | **Gene name** | **Annotation of the Functional Role** | **Ratio**  **log_2_ (2b^+/Δ^/2b^+/+^)** |
| --- | --- | --- | --- |
| MDR3 P-glycoprotein | *Abcb4* | Encodes a protein that transports phospholipids across liver cell membranes for release into bile. | 1.0319 |
| Actin-like protein 6B | *Actl6b* | Gene mutations result in neurodevelopmental deficits, epilepsy, and dendritic loss in human neurons. | 1.1841 |
| Armadillo-like helical domain-containing protein 3 | *Armh3* | Regulates Golgi organization; localized to the Golgi membrane and cytosol. | 1.7692 |
| Beta-site amyloid precursor protein cleaving enzyme 1 | *Bace1* | Plays a key role in the production of amyloid-beta peptides implicated in Alzheimer’s disease. | 1.1617 |
| BCL2-associated X protein | *Bax* | Bax, a pro-apoptotic Bcl-2 family member, triggers apoptosis and is essential for developmental neuronal cell death. | 1.0368 |
| Cell division cycle 37-like 1 protein | *Cdc37l1* | The expression of this gene has been associated with gastric cancer. | 1.2357 |
| Cysteine-serine-rich nuclear protein 1 | *Csrnp1* | This gene is a tumor suppressor, and its expression is increased by estradiol. | 1.1717 |
| Cytochrome P450 family 4 subfamily V member 2 | *Cyp4v2* | This gene encodes a cytochrome P450 enzyme involved in oxidizing substrates in metabolism, particularly in converting fatty acid precursors to n-3 polyunsaturated fatty acids. | 1.0131 |
| DEAD-box helicase 3, Y-linked | *Ddx3y* | It is a crucial RNA helicase involved in RNA processing, translation regulation, and spermatogenesis. | 1.0381 |
| Fibrous sheath-interacting protein 2 | *Fsip2* | FSIP2 is involved in acrosome development during spermiogenesis and may serve as a predictive biomarker for Clear Cell Renal Cell Carcinoma prognosis. | 1.085 |
| Growth arrest-specific protein 8 | *Gas8* | A component of the cytoskeleton involved in ciliary function and microtubule organization. | -1.212 |
| Histone H2B ubiquitinated 2 | *H2bu2* | This histone is involved in chromatin organization and gene regulation. | -1.332 |
| Histone acetyltransferase 1 | *Hat1* | Histone acetyltransferase 1 acts as a succinyltransferase for both histones and non-histones, promoting tumorigenesis. | 1.0848 |
| Inositol 1,4,5-trisphosphate receptor type 3 | *Itpr3* | It is a calcium channel that mediates intracellular calcium release from the endoplasmic reticulum. | 1.0904 |
| Minichromosome maintenance complex component 4 | *Mcm4* | The expression of certain MCM complex components, including MCM4, a helicase essential for DNA replication and genome stability, is linked to the survival of hepatocellular carcinoma (HCC) patients. | 1.1244 |
| Metallothionein 1 | *Mt1* | This protein is involved in heavy metal detoxification, oxidative stress response, and metal ion homeostasis. | -1.3131 |
| Myosin light chain kinase | *Mylk* | Encodes the Ca2+/calmodulin-dependent myosin light chain kinase (MLCK), responsible for phosphorylating the regulatory light chain and initiating smooth muscle cell contraction. | 1.0677 |
| Myoferlin | *Myof* | Myoferlin is involved in membrane repair and fusion processes in muscle cells and is overexpressed in various cancers, particularly pancreatic and triple-negative breast cancer. | 1.1297 |
| N-deacetylase/N-sulfotransferase 4 | *Ndst4* | This protein is a candidate tumor suppressor involved in heparan sulfate modification, which regulates cellular signaling and development. | 1.0822 |
| Optic atrophy 3 protein | *Opa3* | This protein plays a role in mitochondrial function and is associated with optic atrophy. | 1.0101 |
| Ribosomal RNA Processing Protein 1 | *RRP1* | This is a nucleolar protein that helps in processing and assembling ribosomal RNA into functional ribosomes. | -1.0416 |
| Translocation protein SEC62 | *Sec62* | This protein is involved in the translocation of nascent proteins across the endoplasmic reticulum membrane. | 1.5698 |
| Vacuole membrane protein 1 | *Vmp1* | It is involved in autophagy and vacuole membrane formation and is overexpressed in pancreatitis-affected acinar cells, where it mediates zymophagy. | 1.5755 |

**Table S2 | The list of significantly altered proteins and the annotation of their functional role**

| **Protein name** | **Gene name** | **Ratio**  **(*Grin2b*^+/Δ^ /*Grin2b*^+/+^)** |
| --- | --- | --- |
| Disks large homolog 2 | *Dlg2* | 1.03 |
| Disks large homolog 3 | *Dlg3* | 1.01 |
| Disks large homolog 4 | *Dlg4* | 1.12 |
| Disks large-associated protein 1 | *Dlgap1* | 1.10 |
| Disks large-associated protein 2 | *Dlgap2* | 1.03 |
| Disks large-associated protein 3 | *Dlgap3* | 0.98 |
| Disks large-associated protein 4 | *Dlgap4* | 1.00 |
| Homer protein homolog 1 | *Homer1* | 1.08 |
| Homer protein homolog 2 | *Homer2* | 1.09 |
| Homer protein homolog 3 | *Homer3* | 1.26 |
| Neurexin-1 | *Nrxn1* | 1.04 |
| Neurexin-2 | *Nrxn2* | 1.08 |
| Neurexin-3 | *Nrxn3* | 1.11 |
| Neuroligin-2 | *Nlgn2* | 1.03 |
| Neuroligin-3 | *Nlgn3* | 1.14 |
| Protein bassoon | *Bsn* | 0.90 |
| Protein piccolo | *Pclo* | 0.95 |
| Regulating synaptic membrane exocytosis protein 1 | *Rims1* | 1.09 |
| Regulating synaptic membrane exocytosis protein 2 | *Rims2* | 1.00 |
| SH3 and multiple ankyrin repeat domains protein 1 | *Shank1* | 1.17 |
| SH3 and multiple ankyrin repeat domains protein 2 | *Shank2* | 1.10 |
| SH3 and multiple ankyrin repeat domains protein 3 | *Shank3* | 1.16 |
| Synapsin-1 | *Syn1* | 1.08 |
| Synapsin-2 | *Syn2* | 1.23 |
| Synapsin-3 | *Syn3* | 0.83 |
| Synaptic vesicle glycoprotein 2A | *Sv2a* | 0.90 |
| Synaptic vesicle glycoprotein 2B | *Sv2b* | 1.12 |
| Synaptic vesicle membrane protein VAT-1 | *Vat1* | 1.31 |
| Synaptic vesicle membrane protein VAT-1 | *Vat1l* | 0.90 |
| Synaptobrevin homolog YKT6 | *Ykt6* | 0.97 |
| Synaptophysin | *Syp* | 1.09 |
| Synaptophysin-like protein 1 | *Sypl1* | 1.27 |
| Synaptosomal-associated protein 25 | *Snap25* | 0.95 |
| Synaptosomal-associated protein 47 | *Snap47* | 1.10 |
| Synaptotagmin-1 | *Syt1* | 1.14 |
| Synaptotagmin-11 | *Syt11* | 1.21 |
| Synaptotagmin-12 | *Syt12* | 1.13 |
| Synaptotagmin-17 | *Syt17* | 1.04 |
| Synaptotagmin-2 | *Syt2* | 0.92 |
| Synaptotagmin-3 | *Syt3* | 1.04 |
| Synaptotagmin-7 | *Syt7* | 1.14 |
| Syntaxin-12 | *Stx12* | 1.07 |
| Syntaxin-16 | *Stx16* | 1.38 |
| Syntaxin-17 | *Stx17* | 1.05 |
| Syntaxin-1A | *Stx1a* | 1.06 |
| Syntaxin-1B | *Stx1b* | 1.04 |
| Syntaxin-4 | *Stx4* | 1.16 |
| Syntaxin-5 | *Stx5* | 1.15 |
| Syntaxin-8 | *Stx8* | 1.22 |
| Syntaxin-binding protein 1 | *Stxbp1* | 0.89 |
| Syntaxin-binding protein 3 | *Stxbp3* | 1.12 |
| Syntaxin-binding protein 5 | *Stxbp5* | 1.05 |
| Syntaxin-binding protein 5-like | *Stxbp5l* | 1.11 |
| Vesicle-associated membrane protein 1 | *Vamp1* | 1.10 |
| Vesicle-associated membrane protein 2 | *Vamp2* | 1.18 |
| Vesicle-associated membrane protein 4 | *Vamp4* | 1.53 |
| Vesicle-associated membrane protein 7 | *Vamp7* | 1.19 |
| Vesicle-associated membrane protein-associated protein A | *Vapa* | 1.10 |
| Vesicle-associated membrane protein-associated protein B | *Vapb* | 0.99 |
| Vesicular glutamate transporter 1 | *Slc17a7* | 1.07 |
| Vesicular glutamate transporter 2 | *Slc17a6* | 0.72 |

**Table S3 | The list of selected synaptic proteins and their CN ratio in *Grin2b*^+/Δ^ relative to *Grin2b*^+/+^ hippocampal tissue.**

|  | ***Grin2b*^+/+^** | | | ***Grin2b*^+/Δ^** | | |
| --- | --- | --- | --- | --- | --- | --- |
|  | **Male** | **Female** |  | **Male** | **Female** |  |
|  | **Mean ± SEM (*n*)** | **Mean ± SEM (*n*)** | ***p*-value** | **Mean ± SEM (*n*)** | **Mean ± SEM (*n*)** | ***p*-value** |
| **Agonist induced whole-cell currents** | | | | | | |
| NMDA current density (pA/pF) | 52.0 ± 3.4 (32) | 48.3 ± 5.3 (22) | 0.17 | 29.1 ± 3.7 (17) | 26.6 ± 2.2 (42) | 0.49 |
| AMPA current density (pA/pF) | 43.3 ± 4.5 (27) | 54.4 ± 13.2 (13) | 0.93 | 35.0 ± 4.6 (14) | 33.4 ± 2.4 (43) | 0.64 |
| Ifenprodil inhibition (%) | 24.8 ± 2.5 (32) | 25.3 ± 3.8 (19) | 0.91 | 18.5 ± 2.8 (17) | 17.7 ± 1.9 (41) | 0.72 |
| **EPSCs** | | | | | | |
| NMDA eEPSC current density (pA/pF) | 17.2 ± 3.1 (35) | 21.5 ± 6.9 (21) | 0.85 | 20.5 ± 4.5 (32) | 12.6 ± 3.0 (19) | 0.75 |
| AMPA eEPSC current density (pA/pF) | 36.9 ± 5.0 (36) | 37.3 ± 8.5 (22) | 0.8 | 39.8 ± 6.3(33) | 34.7 ± 7.5 (21) | 0.90 |
| Ifenprodil inhibition (%) | 31.1 ± 2.5 (25) | 33.0 ± 2.9 (19) | 0.71 | 31.3 ± 3.0 (25) | 29.5 ± 5.7 (15) | 0.77 |
| Tau weighted (ms) | 208 ± 11 (36) | 238 ± 16 (21) | 0.12 | 147 ± 8 (32) | 142 ± 9 (18) | 0.69 |
| Paired-pulse ratio | 0.91 ± 0.05 (28) | 1.00 ± 0.06 (20) | 0.24 | 1.03 ± 0.06 (26) | 0.92 ± 0.04 (17) | 0.28 |
| **Spontaneous activity** | | | | | | |
| Frequency (Hz) | 3.7 ± 1.0 (21) | 3.1 ± 0.9 (17) | 0.7 | 4.2 ± 1.1 (28) | 2.8 ± 1.2 (19) | 0.04 |
| Amplitude (pA) | 18.6 ± 2.1 (21) | 20.3 ± 1.7 (17) | 0.27 | 28.8 ± 2.0 (28) | 24.6 ± 2.4 (19) | 0.07 |

**Table S4 | Data reported in Fig. 6-8, grouped by sex.** Data from 13–43 neurons per group (*Grin2b*^+/+^ and *Grin2b*^+/Δ^) are presented as mean ± SEM. Genotype comparisons within each sex (male and female) were analyzed using Student’s t-test. Reported *p*-values indicate statistical differences between genotypes within each sex.

| **Test** | | **Male** | | | **Female** | | |
| --- | --- | --- | --- | --- | --- | --- | --- |
|  |  | ***Grin2b*^+/+^** | ***Grin2b*^+/Δ^** |  | ***Grin2b*^+/+^** | ***Grin2b*^+/Δ^** |  |
|  |  | **Mean ± SEM (*n*)** | **Mean ± SEM (*n*)** | ***p*-value** | **Mean ± SEM (*n*)** | **Mean ± SEM (*n*)** | ***p*-value** |
| **Open field test** | Whole arena average speed (m/s) | 3.8 ± 0.2 (10) | 2.9 ± 0.2 (11) | 0.044 | 3.9 ± 0.2 (12) | 4.0 ± 0.2 (10) | 0.89 |
|  | Periphery distance travelled (m) | 165.8 ± 1.0 (10) | 13.3 ± 0.9 (11) | 0.19 | 16.8 ± 0.9 (12) | 17.9 ± 1.0 (10) | 0.54 |
|  | Periphery resting time (s) | 419 ± 12 (10) | 467 ± 13 (11) | 0.02 | 430 ± 11 (12) | 451 ± 12 (10) | 0.44 |
|  | Periphery permanence time (s) | 502 ± 12 (10) | 548 ± 12 (11) | 0.039 | 519 ± 11 (12) | 541 ± 12 (10) | 0.24 |
|  | Periphery average speed (m/s) | 3.2 ± 0.2 (10) | 2.7 ± 0.2 (11) | 0.15 | 3.3 ± 0.2 (12) | 3.4 ± 0.2 (10) | 0.79 |
|  | Center resting time (s) | 58 ± 8 (10) | 35 ± 5 (11) | 0.08 | 46 ± 6 (12) | 35 ± 5 (10) | 0.29 |
|  | Center permanence time (s) | 95 ± 11 (10) | 56 ± 6 (11) | 0.03 | 82 ± 8 (12) | 64 ± 7 (10) | 0.22 |
|  | Whole arena resting time (s) | 482 ± 6 (10) | 507 ± 5 (11) | 0.03 | 477 ± 5 (12) | 483 ± 6 (10) | 0.56 |
|  | Center distance travelled (m) | 7.0 ± 0.5 (10) | 4.5 ± 0.5 (11) | 0.001 | 7.0 ± 0.5 (12) | 6.2 ± 0.5 (10) | 0.43 |
|  | Number of center entries | 46.6 ± 3.1 (10) | 34.0 ± 2.9 (11) | 0.016 | 45.7 ± 2.8 (12) | 44.8 ± 3.1 (10) | 0.89 |
|  | Total distance travelled (m) | 22.8 ± 1.2 (10) | 17.8 ± 1.2 (11) | 0.04 | 23.8 ± 1.1 (12) | 24.2 ± 1.2 (10) | 0.86 |
|  | Center average speed (m/s) | 7.5 ± 0.5 (10) | 6.3 ± 0.5 (11) | 0.23 | 8.4 ± 0.5 (12) | 9.5 ± 0.5 (10) | 0.22 |
|  | Latency to center entry (s) | 16.8 ± 3.9 (10) | 22.9 ± 5.1 (11) | 0.43 | 12.5 ± 2.7 (10) | 33.8 ± 7.9 (10) | 0.04 |
| **IntelliCage system** | Activity | - | - | - | 51.6 ± 1.5 (9) | 51.8 ± 1.5 (8) | 0.93 |
|  | Place preference learning (%) | - | - | - | 41.3 ± 1.1 (9) | 40.3 ± 1.1 (8) | 0.44 |
|  | Place preference reversal (%) | - | - | - | 73.9 ± 0.9 (9) | 69.3 ± 0.9 (8) | 0.26 |
|  | Place preference extinction (%) | - | - | - | 27.2 ± 1.3 (9) | 24.6 ± 1.2 (8) | 0.11 |
|  | Patrolling (%) | - | - | - | 26.9 ± 0.6 (9) | 27.8 ± 0.6 (8) | 0.31 |
| **Freezing time** | Baseline (%) | 1.1 ± 0.4 (10) | 1.9 ± 0.8 (11) | 0.64 | 1.2 ± 0.5 (12) | 0.4 ± 0.3 (10) | 0.18 |
|  | Context (%) | 0.8 ± 0.2 (10) | 0.9 ± 0.2 (11) | 0.89 | 1.5 ± 0.3 (12) | 1.8 ± 0.4 (10) | 0.59 |
|  | Cue (%) | 7.5 ± 2.4 (10) | 10.8 ± 2.5 (11) | 0.34 | 15.7 ± 6.0 (12) | 12.7 ± 2.9 (10) | 0.49 |
| **PPI** | 70 dB (%) | 46.6 ± 1.9 (11) | 29.6 ± 2.4 (11) | <0.001 | 46.2 ± 1.9 (12) | 35.9 ± 2.2 (8) | < 0.001 |
|  | 78 dB (%) | 56.6 ± 1.8 (11) | 51.4 ± 1.8 (11) | >0.05 | 55.8 ± 1.5 (12) | 48.1 ± 1.6 (11) | < 0.001 |
|  | 82 dB (%) | 66 ± 2 (11) | 53 ± 2 (11) | <0.001 | 56 ± 2 (12) | 56 ± 2 (11) | >0.05 |
|  | 85 dB | 63 ± 2 (11) | 58 ± 2 (11) | >0.05 | 58 ± 2 (12) | 65 ± 1 (11) | < 0.001 |
| **Elevated plus maze (time** **spent in each zone)** | Open arms (s) | 18 ± 1 (15) | 26 ± 2 (11) | 0.09 | 31 ± 4 (16) | 35 ± 5 (12) | 0.51 |
|  | Closed arms (s) | 238 ± 11 (15) | 235 ± 12 (11) | 0.09 | 222 ± 21 (16) | 200 ± 22 (12) | 0.51 |
|  | Center zone (s) | 34.6 ± 1.9 (15) | 39.9 ± 2.7 (11) | 0.09 | 27.9 ± 3.3 (16) | 37.1 ± 4.9 (12) | 0.51 |
| **Elevated plus maze (visits)** | Open arms | 6.2 ± 0.6 (15) | 5.3 ± 0.7 (11) | 0.17 | 5.0 ± 0.5 (16) | 8.2 ± 0.8 (12) | 0.019 |
|  | Closed arms | 12.9 ± 0.6 (15) | 11.4 ± 0.7 (11) | 0.17 | 11.6 ± 0.8 (16) | 13.1 ± 0.9 (12) | >0.05 |
| **Elevated plus maze (distance)** | Open arms (m) | 1.6 ± 0.3 (15) | 1.6 ± 0.4 (11) | 0.45 | 1.9 ± 0.3 (16) | 1.7 ± 0.3 (12) | 0.98 |
|  | Closed arms (m) | 12.2 ± 0.3 (15) | 11.5 ± 0.4 (11) | 0.45 | 11.4 ± 0.7 (16) | 12.1 ± 0.8 (12) | 0.98 |
| **Object recognition** | Duration familiar (s) | 6.7 ± 1.7 (15) | 3.6 ± 1.3 (11) | 0.28 | 18.3 ± 3.4 (16) | 10.2 ± 2.5 (12) | 0.15 |
|  | Duration novel (s) | 4.7 ± 0.9 (15) | 5.5 ± 1.2 (11) | 0.74 | 5.7 ± 0.9 (16) | 4.9 ± 1.1 (12) | 0.71 |
|  | Discrimination index | 0.60 ± 0.04 (15) | 0.6 ±0.1 (11) | 0.32 | 0.80 ± 0.03 (16) | 0.60 ± 0.04 (12) | 0.02 |
| **Social discrimination** | Duration juvenile (s) | 73.4 ± 7.8 (16) | 64.3 ± 9.4 (11) | 0.59 | 63.6 ± 7.8 (16) | 64.3 ± 9.4 (11) | 0.59 |
|  | Duration object (s) | 28.8 ± 4.2 (16) | 18.8 ± 4.2 (11) | 0.15 | 37.8 ± 4.7 (16) | 21.5 ± 4.5 (11) | 0.14 |
|  | Discrimination index | 0.70 ± 0.03 (16) | 0.80 ± 0.03 (11) | 0.22 | 0.60 ± 0.03 (16) | 0.80 ± 0.03 (11) | 0.01 |
| **Nestlet shredding** | % of shredded nestlet | 18.4 ± 2.2 (15) | 14.7 ± 2.2 (11) | 0.42 | 10.9 ± 1.5 (16) | 7.6 ± 1.4 (12) | 0.21 |

**Table S5 | Numerical data underlying Fig. 9–10 and Fig. S3–S6.** Data from 8–16 animals per group (*Grin2b*^+/+^ and *Grin2b*^+/Δ^ mice) are presented as mean ± SEM. Genotype comparisons within each sex (male and female) were made using ANOVA. Reported *p*-values indicate statistical differences between genotypes within each sex.

### Supplementary Figures

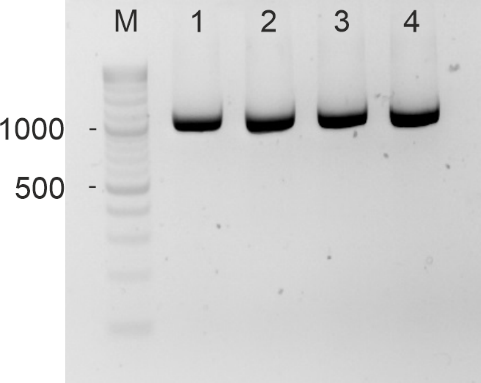

**Figure S1 | A representative agarose gel shows the amplification of a 1200 nt fragment of *Grin2b* mRNA.** The mRNA was obtained from hippocampal tissue of *Grin2b*^+/Δ^ female mice at 4 months of age. The forward primer was located in exon 7 and the reverse primer in exon 13 (see Table S1 for sequences). For direct Sanger sequencing of the PCR amplicons, an additional forward primer located in exon 12 was used (see Table S1). M denotes the molecular weight marker; lanes 1–4 represent samples from four individual mice.

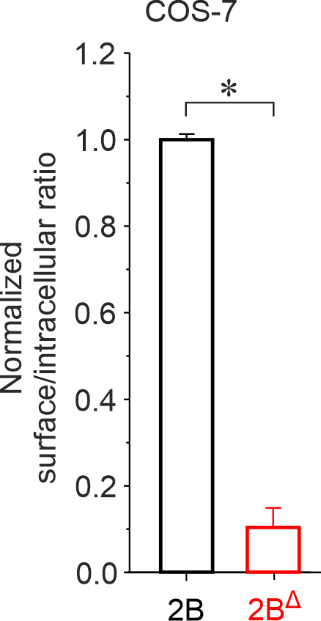

**Figure S2 | GluN2B^Δ^ subunit disrupts NMDAR surface expression in COS-7 cells.** COS˗7 cells were co-transfected with GluN1 and wild-type GluN2B-GFP (2B) or mutant GluN2B^Δ^-GFP (2B^Δ^) subunits. The graph shows the relative surface expression of GluN1/GFP-GluN2B and GluN1/GluN2B^Δ^-GFP receptors, quantified using a colorimetric assay. Data are presented as mean ± SEM. Asterisk (*) indicates a statistically significant difference (Student's t-test; *n* = 4 independent experiments, with the relative surface expression measured in quadruplicate per experiment).

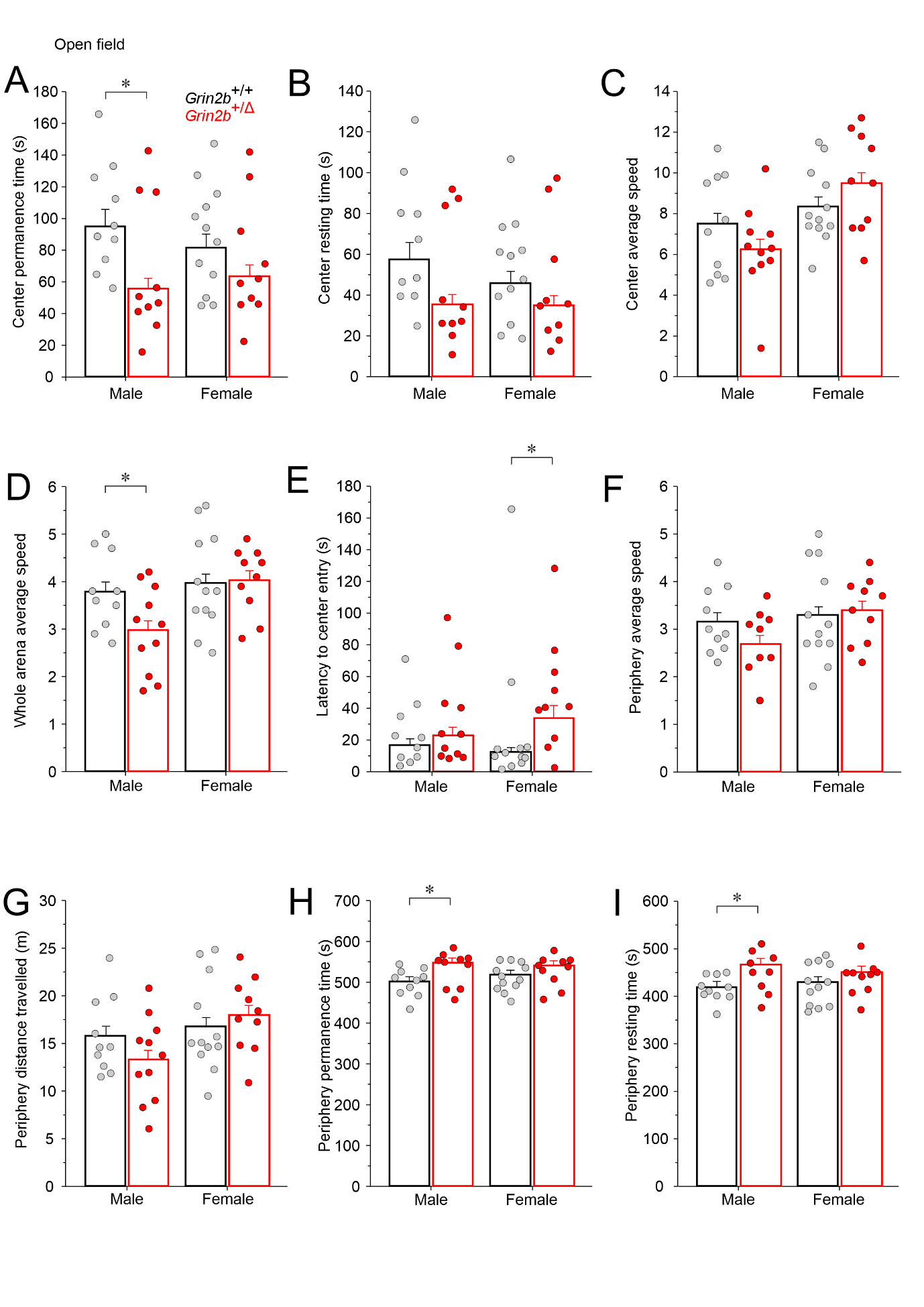

**Figure S3 | *Grin2b*^+/Δ^ mice display hypoactivity and increased anxiety. Supplementary data from the open field test.** (A–I) Center permanence time, center resting time, center average speed, whole arena average speed, latency to first center entry, periphery average speed, periphery distance traveled, periphery permanence time, and periphery resting time were measured in male and female *Grin2b*^+/Δ^ and *Grin2b*^+/+^ mice. Data are presented as mean ± SEM. Number of animals *Grin2b*^+/+^/*Grin2b*^+/Δ^: *n* = 10/11 males and *n* = 12/10 females. Asterisks (*) indicate significant genotype differences (Student’s t-test). The numerical data underlying this figure are shown in Table S5.

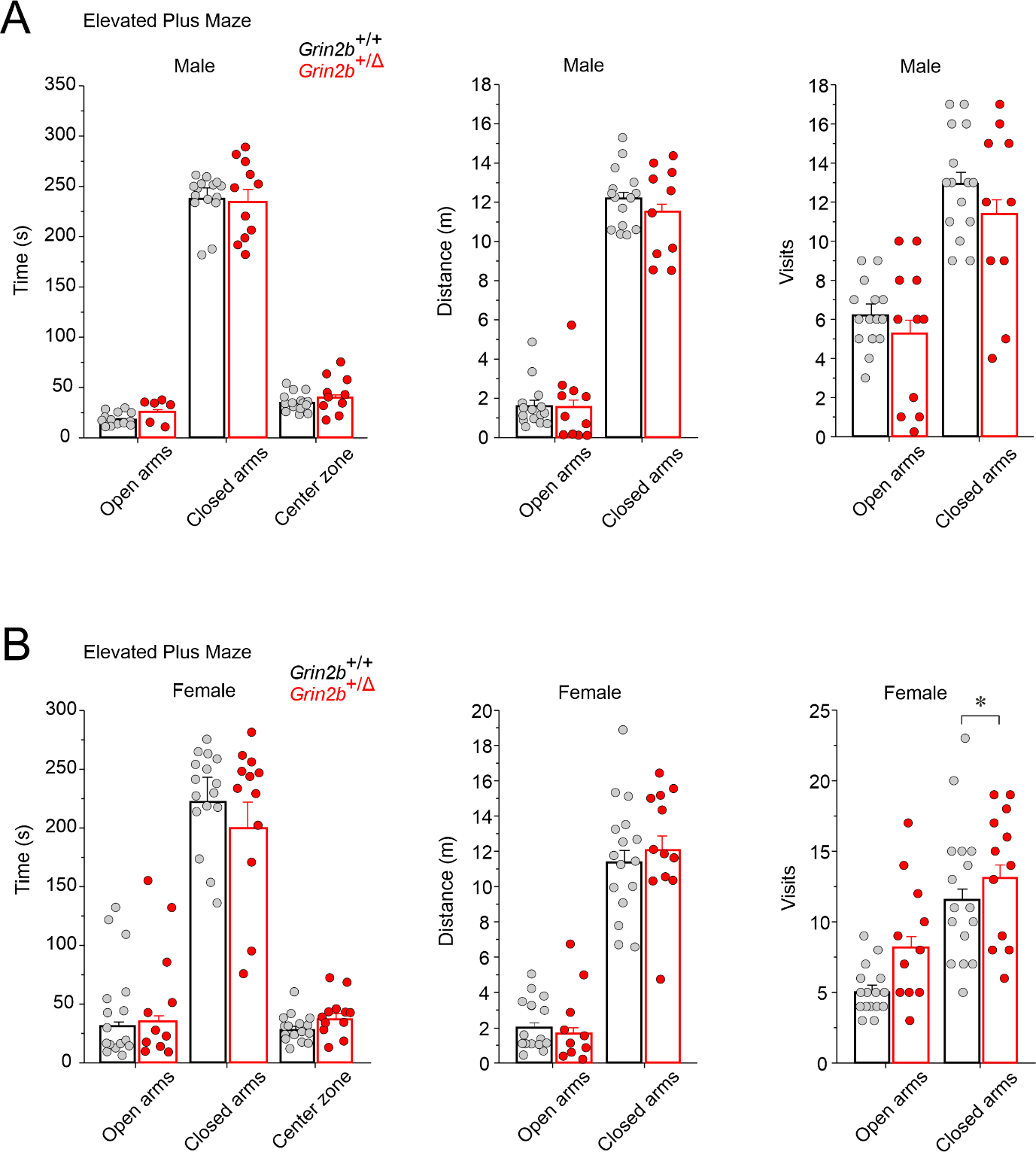

**Figure S4 | Elevated plus maze analysis of anxiety-like behavior in *Grin2b*^+/Δ^ mice.** Bar graphs show measures of anxiety for male (A) and female (B) *Grin2b*^+/Δ^ mice, including time spent, distance traveled, and number of entries into the closed and open arms. Data are presented as mean ± SEM and were analyzed by ANOVA followed by post hoc LSD test. Number of animals *Grin2b*^+/+^/*Grin2b*^+/Δ^: *n* = 15/11 males and *n* = 16/12 females. Asterisk (*) indicates a significant genotype difference. The numerical data underlying this figure are shown in Table S5.

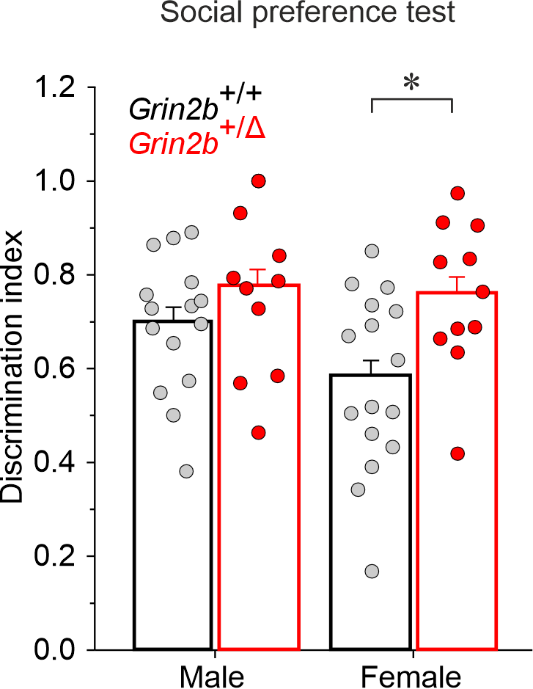

**Figure S5 | Social behavior in *Grin2b*^+/Δ^ mice.** Social preference was assessed, with the discrimination index representing the time spent exploring a juvenile target relative to an object. Data are presented as mean ± SEM and were analyzed by Student’s t-test. Number of animals *Grin2b*^+/+^/*Grin2b*^+/Δ^: 16/11 males, 16/11 females. Asterisk (*) indicates a significant genotype difference. The numerical data underlying this figure are shown in Table S5.

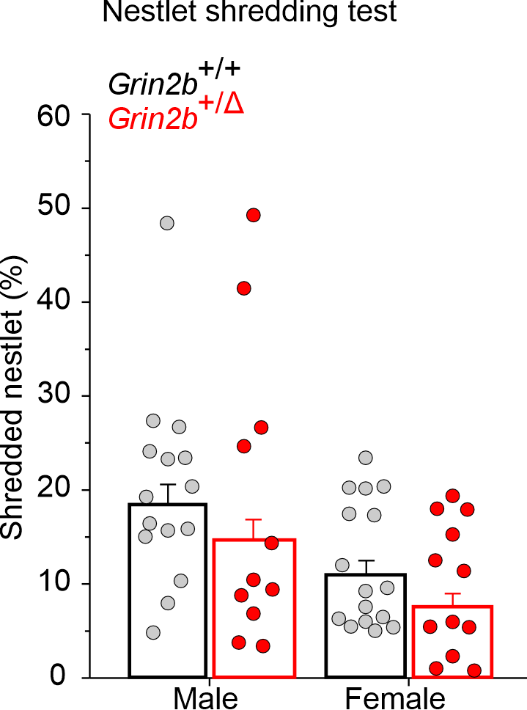

**Figure S6 | Repetitive behavior in *Grin2b*^+/Δ^ mice.** Performance in the nestlet shredding test, measured as the % weight of unused nesting material (initial weight = 10 g). Data are presented as mean ± SEM and were analyzed by Student's *t-test*. Number of animals *Grin2b*^+/+^/*Grin2b*^+/Δ^: *n* = 15/11 males, *n* = 16/12 females. The numerical data underlying this figure are provided in Table S5.
